# Plant defense synergies and antagonisms affect performance of specialist herbivores of common milkweed

**DOI:** 10.1101/2021.09.13.460116

**Authors:** Collin Edwards, Stephen Ellner, Anurag Agrawal

**Affiliations:** Department of Ecology and Evolutionary Biology, Cornell University, Ithaca NY, USA; Department of Biology, Tufts University, Medford MA, USA

**Keywords:** Defense synergy hypothesis, Random Forest, common milkweed *(Asclepias syriaca)*, swamp milkweed beetle *(Labidomera clivicollis)*, monarch butterfly *(Danaus plexippus)*, plant-insect interactions, chemical ecology, synergy

## Abstract

As a general rule, plants defend against herbivores with multiple traits. The defense synergy hypothesis posits that some traits are more effective when co-expressed with others compared to their independent efficacy. However, this hypothesis has rarely been tested outside of phytochemical mixtures, and seldom under field conditions. We tested for synergies between multiple defense traits of common milkweed *(Asclepias syriaca)* by assaying the performance of two specialist chewing herbivores on plants in natural populations. We employed regression and a novel application of Random Forests to identify synergies and antagonisms between defense traits. We found the first direct empirical evidence for two previously hypothesized defense synergies in milkweed (latex by secondary metabolites, latex by trichomes), and identified numerous other potential synergies and antagonisms. Our strongest evidence for a defense synergy was between leaf mass per area and low nitrogen content; given that these “leaf economic” traits typically covary in milkweed, a defense synergy could reinforce their co-expression. We report that each of the plant defense traits showed context-dependent effects on herbivores, and increased trait expression could well be beneficial to herbivores for some ranges of observed expression. The novel methods and findings presented here complement more mechanistic approaches to the study of plant defense diversity, and provide some of the best evidence to date that multiple classes of plant defense synergize in their impact on insects. Plant defense synergies against highly specialized herbivores, as shown here, are consistent with ongoing reciprocal evolution between these antagonists.

## Introduction

Plant defenses structure plant and insect communities, shape the way that primary productivity flows through to the rest of the food web, and can strongly impact the success of individual plants in the face of herbivory (Kursar et al. 2009, Kant et al. 2015, Coley et al. 2018). Consequently, ecologists have spent decades developing and evaluating theories to explain when and how plants should defend themselves (Agrawal 2011). One consistent finding is that individual plants generally have multiple, diverse defense traits (Duffey and Stout 1996, Agrawal and Fishbein 2006, Salazar et al. 2018). Explaining why plants invest in multiple defenses is an open question, especially since there is evidence that selection has favored specific combinations of defense traits (“defense syndromes”: see Agrawal and Fishbein 2006, da Silva and Batalha 2011, Sanczuk et al. 2020, cf. Moles et al. 2013). Several non-exclusive hypotheses have been proposed to explain both the prevalence of multi-trait defense strategies and the specific combinations of defenses that are observed (Agrawal 2011).

Jones and Firn (1991) articulated the problem of multiple phytochemical defenses clearly: individual plants often express many different types of secondary metabolites, and it is difficult to explain why. One of the leading explanations for multiple secondary metabolites is synergistic interactions between different compounds (Duffey and Stout 1996, Richards et al. 2016, Wetzel and Whitehead 2020). For example, the furanocoumarin Xanthotoxin is more toxic when mixed with other furanocoumarins than expected from its effects alone (Berenbaum et al. 1991); similarly, amides in *Piper* plants are more effective in combination than separately (Scott et al. 2002, Dyer et al. 2003, Richards et al. 2010, Whitehead and Bowers 2014). Synergies have also been detected in phytochemical defenses against microbes (e.g., Stermitz et al. 2000, Muroi and Kubo 1993) and fungi (Cipollini and Stiles 1992). Conversely, ecologists have also detected antagonistic relationships, where compounds reduce the efficacy of each other, including among furanocoumarins (Diawara et al. 1993), amides (Whitehead and Bowers 2014), and between pyrrolizidine alkaloids and chlorogenic acid (Liu et al. 2017). Tests for synergies and antagonisms between secondary metabolites are still relatively rare (Richards et al. 2016), but there have been almost no tests for synergies between classes of defenses (e.g., physical and chemical defenses), even though there is every reason to expect them to exist (Rasmann and Agrawal 2009, Agrawal 2011, Richards et al. 2016, but see Green et al. 2001, Steppuhn and Baldwin 2007).

Studies of synergies between phytochemicals generally measure the responses of herbivores to artificial diets with manipulated concentrations of phytochemicals in an ANOVA design (e.g. Dyer et al. 2003, Berenbaum et al. 1991, Stamp and Yang 1996, Scott et al. 2002). Accordingly, synergy or antagonism is detected based on the interaction term of the two (or more) treatments (for a discussion on the appropriate null model for this experimental design, see Pennings 1996, Hay 1996, Nelson and Kursar 1999). For experimental designs with continuously varying defense traits, the logical extension of this approach is regression of herbivore performance (growth, survival, etc.) on levels of defense traits (e.g., concentrations of different phytochemicals) with a term for the interaction between two traits which captures potential synergies or antagonisms (Richards et al. 2016). However, implicit in this regression modeling approach is the assumption of a *bilinear* interaction: that the synergistic benefit (or antagonism) of one additional unit of defense trait A for each unit of defense trait B is the same regardless of the initial levels of A or B (e.g. Fig. 2a, 2d). However, other relationships are biologically reasonable. The synergistic benefit of one trait might be a threshold effect (e.g. Fig. 2b). This could occur if sufficient quantities of a defense causes a switch in herbivore behavior (e.g. *Trichopplusia ni* exhibit trenching behavior on latex-bearing plant species and not on others) and the effect of a second defense is impacted by herbivore behavior (Dussourd and Denno 1994). Alternately, a synergy might have diminishing marginal returns (Fig. 2c), with the benefit of the synergy most prominent when increasing from low to moderate levels of trait expression.

To identify complex relationships like these in experimental data, in this paper we introduce tests for synergies and antagonisms using Random Forest regression. Random Forests are a popular form of machine learning that fit data using an ensemble (effectively, model averaging) of many decision trees (James et al. 2021). Because of this structure, Random Forest models are very flexible and can capture non-linear trends in data. Recent innovations have provided a framework for harnessing the flexible nature of Random Forests for inference about interactions, as we do here (Hooker 2007, Hooker and Mentch 2019). To our knowledge, this approach for identifying interactions with Random Forests has not been previously used in ecology, so we include thoroughly annotated code on Figshare.

Our experiments involved herbivores feeding on common milkweed, *Asclepias syriaca,* a well-characterized system for studying plant defenses. Milkweed produces toxic phytochemicals -- cardenolides -- that are potent defenses and severely limit which species of herbivores that can feed on it, leading to a simplified herbivore community (Agrawal 2017). In addition to cardenolides, *A. syriaca* also employs other defenses: trichomes on the leaf surfaces may reduce traction and insect access to feeding (Agrawal et al. 2009), latex exuded from channels (laticifers) in damaged tissues can cement insect mouth parts shut (Agrawal and Konno 2009), and leaf toughness and low nutrient content may reduce insect preference or performance (Agrawal and Fishbein 2006). Furthermore, expression of these defensive traits varies substantially in natural conditions within species, allowing experiments to capture broad ranges of defense traits in a field setting.

Two putative defense synergies have been proposed for milkweed plants (Agrawal 2011): latex with trichomes, and latex with cardenolides. First, freshly hatched monarch butterfly caterpillars were observed to first carefully chew off trichomes in a small feeding region, and then to bite into the leaf tissue (Zalucki et al. 2002). At this point latex exudes, and the caterpillar must struggle free before it can consume the leaf tissue. Many caterpillars fail to survive this first exposure to latex (Zalucki et al. 2001). Trichomes and latex may thus act synergistically, with higher densities of trichomes leaving neonate caterpillars more exhausted when they encounter the first-bite latex, and thus less able to struggle free. There is indirect evidence for a synergy between these traits, as, across *Asclepias* species, these two traits also show correlated evolution (Agrawal and Fishbein 2006). Second, *A. syriaca* latex contains cardenolides that mirror the concentrations in leaves (Agrawal et al. 2014, Züst et al. 2019), so latex may be a vehicle to deliver toxins directly into the mouth of feeding herbivores in addition to physically gumming up mouthparts. In this way, latex acts as both a chemical and physical barrier when cardenolide expression is high, suggesting a synergy between latex and cardenolide expression. There are indications that this mechanism may be present in other plants with latex and secondary metabolites, as reduced levels of secondary metabolites in latex led to increased herbivory in the common dandelion *(Taraxacum officinale)* (Huber et al. 2016). While latex by trichome and latex by cardenolide synergies in milkweeds were proposed over the last two decades, there has been no direct empirical evidence for either. Both synergies include latex, which is most prominently a defense against chewing herbivores; the latex by trichome synergy is specifically hypothesized from observations of monarch caterpillar behavior, and may not be relevant for other herbivores with different feeding behaviors.

There are not specific antagonisms hypothesized for *A. Syriaca,* or even widely discussed in the broader literature outside of antagonisms between individual secondary metabolites (e.g. Diawara et al. 1993, Calcagno et al. 2002, Whitehead and Bowers 2014, Liu et al. 2017). For specialist insects that are able to sequester plant toxins, we might expect an antagonistic relationship between secondary metabolites and traits that increase predation -- indirect defenses that attract natural predators. Indeed, any plants traits that are somewhat effective against herbivores, but also have negative effects on natural enemies of herbivores could be antagonistic with indirect defenses.

In this study we evaluate plant defense synergies and antagonisms using herbivore performance on common milkweed *A. syriaca* under field conditions, employing both well-established regression approaches and a novel application of Random Forests. While the majority of existing studies of synergies have used artificial diets, we measure herbivore performance under field conditions, relying on natural variation in defense traits in wild plants. Using observational field studies sacrifices controlled conditions, but provides a realistic context for plant defenses, and allows us to study defenses that are not easily manipulated in the lab. Using two specialist chewing herbivores (the swamp milkweed beetle *Labidomera clivicollis* and the monarch butterfly *Danaus plexippus)* we tested the defense synergy hypothesis for two predicted synergies: latex by cardenolide and latex by trichome. We also explored other potential synergies and antagonisms between cardenolides, trichome density, latex, leaf toughness, carbon, nitrogen, and leaf mass per area. Where synergies and antagonisms have not been previously hypothesized, we sought evidence for trait interactions that are worthy of more targeted experiments. Our methods provide a road map for future tests of the synergy hypothesis between classes of plant defenses, and for other evaluations of trait synergies in ecology.

## Material and Methods

### Study Species

*A. syriaca* is a long-lived herbaceous plant native throughout eastern North America, often found in recently disturbed habitats including roadsides and fields. Larvae of *D. plexippus* and *L. clivicollis* are both specialist herbivores that commonly feed on the leaf tissues of *A. syriaca* in our study area. Both of these herbivores have modifications to their NA, K-ATPase (sodium pumps) that increase their tolerance of cardenolides (Dobler et al. 2012), but do not provide complete immunity (Zalucki et al. 2001, Agrawal 2005, Tao et al. 2016, Jones et al. 2019).

### Field experiments

In September 2015, we conducted an experiment to assay the growth and survival of neonate monarch caterpillars, *D. plexippus,* on 117 *A. syriaca* plants near Ithaca, NY (42.39 N, 76.39 W). Caterpillars were obtained from a laboratory colony (see Jones and Agrawal 2019). We selected plants from a natural population haphazardly, using distance and visual phenotype to avoid selecting ramets from the same plant; we also avoided ramets that showed high levels of damage or early senescence. We placed two freshly hatched monarch caterpillars on one of the top pair of fully expanded leaves and enclosed them in a fine mesh bag. We used the other leaf in that pair to measure plant traits, including taking latex and toughness measurements, and collecting a leaf disk for measuring trichome density and leaf mass per area (LMA; the inverse of specific leaf area). We then removed that paired leaf to be dried for measurement of cardenolides, carbon, and nitrogen.

Latex, leaf toughness, LMA, and cardenolides were quantified following standardized methods (for a detailed description, see Hahn et al. (2019)). In brief, latex was measured by removing 2-3 mm of the leaf tip and measuring the weight of the resulting latex exudate; leaf toughness was measured using a penetrometer (Chatillon type 5/6) once on either side of the leaf and averaging the results; LMA (kg dry mass/m^2^) was measured by drying and weighing a 31.67 mm^2^ leaf punch; cardenolides were separated and measured from dried and ground leaf tissue using High Performance Liquid Chromatography (HPLC) following the methods described in Züst et al. (2019). Observed cardenolide compounds did not match currently identified compounds, so we report their identity based on the retention time (e.g. during HPLC analysis, cardenolide 10.7 eluted at 10.7 minutes). Trichome density was measured by first imaging the underside of leaf discs with the Ziess SteREO Discovery.V20, then using ImageJ to draw vertical and horizontal transects through the middle of the image and count all hairs that crossed the transects (Schindelin et al., 2012). These transect counts were highly correlated with densities obtained by counting all hairs in a quadrant of the leaf disc, but were faster and more robust to overlapping hairs (CBE, unpublished). Ground and dried leaf samples were analyzed at the Cornell Isotope Laboratory (COIL) for concentrations of carbon and nitrogen.

Monarch caterpillars generally remained on the plants for six days, and then were removed and weighed. In a few cases, field conditions prevented collection on day six, and caterpillars were instead collected on day seven; where appropriate, we account for this discrepancy with random effects. Missing caterpillars were presumed dead. In several cases, plants died during the experiment, necessitating removing them from the study. In total, we obtained 165 weighed larvae and 183 measures of survivorship for 92 plants.

In September 2016, we conducted an experiment to assay growth and survival of neonate swamp milkweed beetles, *L. clivicollis,* on 129 *A. syriaca* plants with the same methods as above, with the following exceptions. First, we used a separate field site approximately 300 meters away, at Dunlop Meadow (a Cornell University Natural Area). Second, because *L. clivicollis* aggregate in their early life stages, we placed 5 larvae in each enclosure rather than two. In a few cases, loss of larvae during placement left us with only 4 larvae in an enclosure. Third, due to unusual weather patterns, many of the plants experienced accelerated senescence during the experiment; we did not include these samples. Fourth, due to contamination of tissue samples, we were unable to measure LMA or carbon and nitrogen concentrations. In total, we obtained 133 weighed larvae and 464 records of survivorship for 95 plants.

### Analysis

We took two approaches to analyzing our data. First we looked for bilinear synergies and antagonisms using a traditional linear regression approach, where synergies and antagonisms were represented with a statistical interaction term. Second, we implemented a novel (for ecology) extension of Random Forests to identify synergies and antagonisms that were not bilinear. In all cases, our metric of herbivore growth was log final mass. For regression analyses, we rescaled plant traits so that their mean was zero and their standard deviations was 1; for Random Forest analyses we rescaled plant traits so that their maximum was 1.

In total we had two data sets, each with two measures of herbivore response (survival and growth). For simplicity, in the rest of this paper we will refer to these data sets as “Monarchs”, and “Beetles”. These two data sets have some overlap in the defense traits measured, but the cardenolide compounds expressed in each year were different, and due to sample contamination we were unable to measure LMA, carbon, or nitrogen in the Beetles data. To facilitate interpretation, we re-contextualize percent nitrogen as “non-nitrogen” (100 – percent nitrogen), so that all analyzed traits are expected *a priori* to have a negative effect on herbivores. Although carbon and nitrogen concentrations are sometimes combined into the carbon-to-nitrogen ratio (C:N) as a proxy for nutritional quality, here we treat carbon and (non-)nitrogen as independent, as we and others have done in the past (e.g. Agrawal 2004a, Agrawal 2004b). This decision was driven by three factors. First, the purpose of our study was to quantify trait interactions, and pre-combining two traits in a fixed fashion as in the C:N ratio instead pre-suppose a specific trait interaction. Second, herbivores sometimes respond to carbon and nitrogen separately; analyzing them as separate traits allowed us to detect this. Third, we found that variation in C:N was driven almost exclusively by variation in (non-)nitrogen (correlation coefficient of 0.95, p<0.001), and was uncorrelated with carbon (correlation coefficient of −0.077, p = 0.3) (Fig. S1).

### Trait Correlations

We computed pairwise across-plant trait (phenotypic) correlations for each of our data sets, using Pearson correlation coefficients. Each of our data sets contained several cardenolide compounds that were highly correlated (Monarchs: cardenolides 17.4, 17.7, 18.4; Beetles: cardenolides 10.7, 18.1). For these suites of covarying cardenolides, we used principal component analysis (PCA) to identify a single measure -- the first PC -- that represented the majority of the variation in those cardenolides. All further analysis used these first PCs in place of the individual cardenolides in question, labeled “cardenolide suite 2015” and “cardenolide suite 2016”. For the remaining cardenolides which were not highly correlated (Monarchs: cardenolides 10.5, 18.6, Beetle: cardenolides 8.4, 10.2), we treated each compound as a separate defense trait.

### Linear regression

For each trait pair (e.g., latex by trichomes, latex by cardenolide for each of the independent compounds (10.5, 18.6, 10.2, 8.4) and the two cardenolide suites, etc), we fit a regression model that included main effects for the two focal traits, and an interaction between those two traits (as well as a random effect of plant to account for structure in the data). We then used marginal hypothesis testing to identify the significance of the interaction term. Because this analysis involves many comparisons, we discourage viewing P values as part of a Null Hypothesis Significance Test (NHST) to “definitively” prove (or find lack of proof for) specific synergies or antagonisms. Instead, we emphasize the value of P-values as evidence (or a lack of evidence) for synergies or antagonisms (Muff et al. 2022); synergies and antagonisms with the smallest P values are those that the data most strongly supports. Following Muff et al. (2022), we designate P < 0.01 as “strong evidence”, 0.01 < P < 0.05 as “moderate evidence”, and 0.05 < P < 0.1 as “weak evidence”. We note that focusing on P-values as evidence rather than a binary NHST is more consistent with the actual definition and statistical best practices for P-values, and is a good idea even without multiple comparisons.

We develop a technical basis for interpreting the regression interaction coefficient below, then describe the intuition this provides. For a pair of scaled traits *x*_1_ and *x*_2_ with main effect coefficients *β*_1_ and *β*_2_ (these represent the direct effect on the herbivore of variation in either trait when the other is at its average) and interaction coefficient *β*_12_, we can rewrite our expectation for the expected herbivore performance *y* from the standard regression form

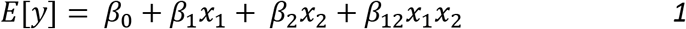

to

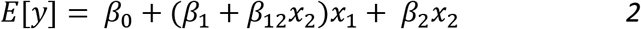

From Equation 2 we see that the expected effect of varying the level of trait 1 (*x*_1_) depends on both *β*_1_ (the direct effect of trait 1) and the product of the interaction coefficient and *x*_2_. Because we are working with scaled trait values, *x*_1_ and *x*_2_ represent deviations from the mean trait values across our experiment. When *β*_12_ is negative, if trait 2 is above the mean for that trait (*x*_2_ is positive), higher-than-average values of trait 1 (positive values of *x*_1_) are associated with reduced herbivore performance relative to when trait 2 is at its mean. Because equation 1 can just as easily be rewritten to focus on variation in trait 2, in the form

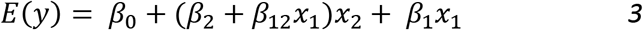

we see the interpretation is symmetrical, and *β*_12_ is modifying the effect of *each* trait in the context of the other. When herbivore survival is the response, *y* is the logit transformation of survival, but this interpretation still applies to survival because the logit and its inverse are monotonic.

To identify an interaction as a synergy or an antagonism, we look to the sign of the interaction coefficient, *β*_12_ above. Because the response is insect performance, a negative coefficient for the interaction term represents a potential synergy between traits; herbivore performance is lower when the traits are both above average expression levels, compared to what we would expect from the separate effect of each of the trait’s expression levels. Similarly, a positive interaction coefficient represents an antagonism between traits. Because each trait is rescaled to have mean 0 and standard deviation 1, the interaction term can be interpreted as modifying the effect of one trait based on how far the other trait is from the mean. Because the classification of a trait as a defense against herbivores could be context-dependent (see below), we took the inclusive approach of identifying synergies and antagonisms based solely on the sign of the interaction coefficient.

An underlying assumption of many plant-insect ecologists (including ourselves) is that for a given pair of plant and herbivore species, a plant trait generally is or is not a defense. However, a key insight from Equation 2 is that, depending on the strength of trait interactions, a focal trait can be harmful or helpful to herbivores depending on the expression levels of other traits. From Equation 2, we find that the net effect on herbivores of increasing trait 1 changes sign (e.g. switches from being defensive to beneficial to herbivores or vice versa) when *β*_1_ = –*β*_12_*x*_2_. This switching point depends on the coefficients of the main effect (*β*_1_) and interaction (*β*_12_), as well as how far trait 2 is from its mean (*x*_2_). For a linear model with a non-zero interaction term, a switching point will always exist, but it may be associated with extreme, biologically unreasonable trait values. We calculated the switching point for each trait pairing, treating a switching point as biologically reasonable if it occurred when the non-focal trait value was within one standard deviations of its mean. We designated a trait as “context-dependent” if, when paired with each other trait, at least one of the switching points was biologically likely. If no switching points were biologically reasonable, we designate the trait as “defensive” or “helpful” based on whether it was harmful or helpful to herbivore performance when other traits were at their mean.

Because the interaction coefficient is interpreted on the scale of the linear predictor rather than the response, its magnitude cannot be used to compare the strength of synergies between survival (logit-link) and growth (identity-link) responses, or to make comparisons with equation-free models like Random Forests. To provide a more interpretable and robust measure of the strength of synergies and antagonisms, for each trait pair and response (growth or survival) we developed two new metrics, which we introduce here. The first, the “Predictable Response Range” (PRR), captures the variation in response values that can be attributed to a model. The second, the percent of Predicted Response Variation explained by the interaction (%PRV), captures the relative role of non-additivity (e.g., synergy or antagonism) in explaining model predictions.

PRR is a metric for any model that describes the range of model predictions for the data (Fig 1). Much like the coefficient of determination, R^2^, PRR is a tool to understand the variation in the dependent variable that can be explained by the model. Unlike R^2^, however, PRR focuses on explaining how the dependent variable is expected to change (e.g. from modeled effects, after excluding the variation from random noise and un-modeled variables) if predictor values where changed from the least favorable to the most favorable in the data. For regression models, PRR was simply the range of ŷ, the model predictions of the data (Figure 1). For the Random Forest analysis, calculation was slightly more complex (see Appendix S1) but produced an analogous range. When applied to our models of two defense traits and their interaction, PRR captured how much the plant with the highest predicted herbivore performance could expect to benefit if it were able to modify its two modeled traits to those of the plant with the lowest predicted herbivore performance. PRR thus represented all explained effects of the trait combination on herbivores in our data: the combination of direct effects of the traits, bilinear synergies or antagonisms, and in the case of Random Forests, any other non-additive relationships.

**Figure 1:**
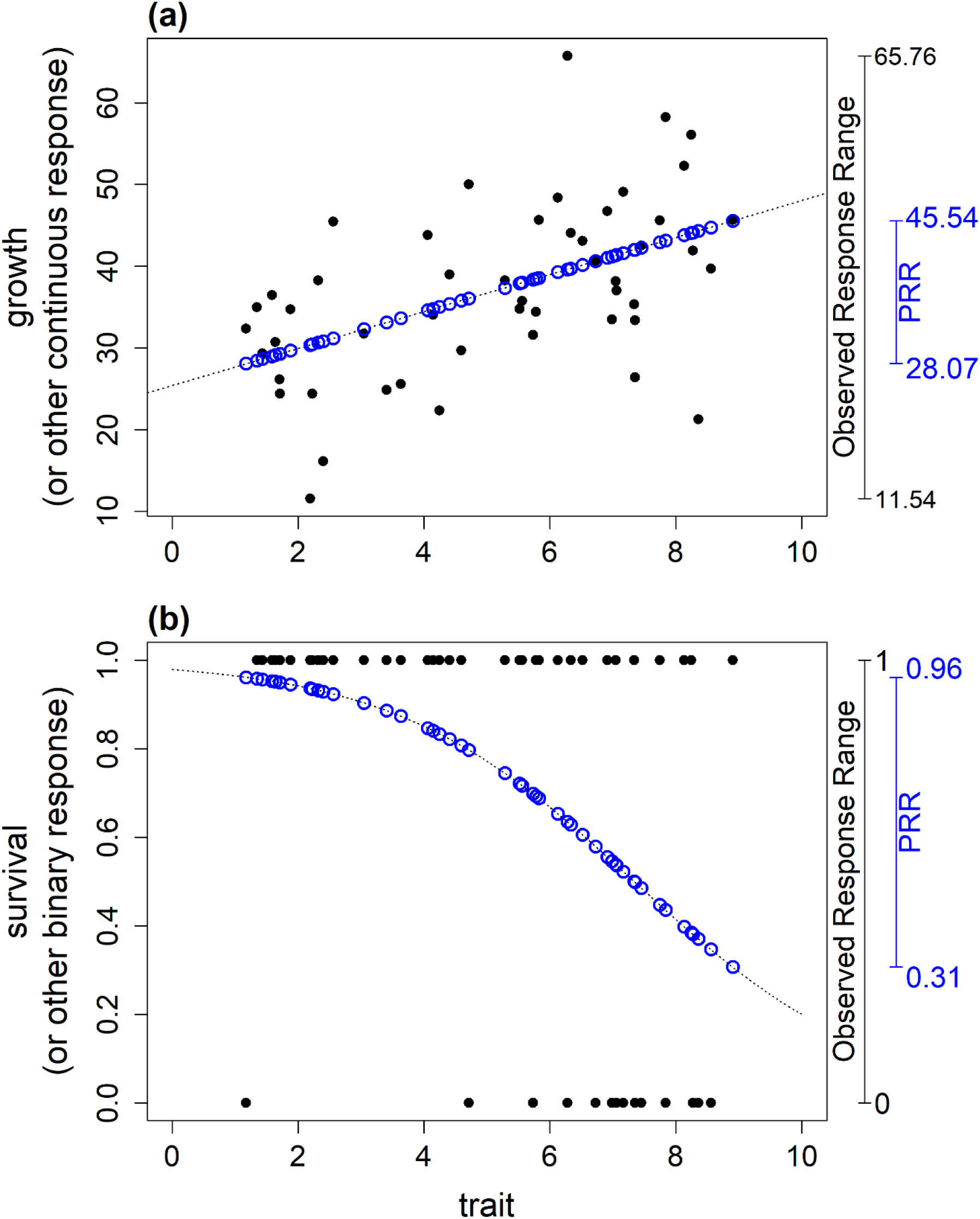
Example of the Predictable Response Range (PRR) for simplified simulated data. Black points represent simulated data; blue circles show the predictions for those points from a fitted model. PRR captures the range of response values (e.g., growth or survival) predicted for data points. Here 30 trait values were drawn from a uniform distribution ranging from 1 to 9, and (a) growth and (b) survival responses to traits were simulated. The dashed line shows models fit to the data of the form ‘response ~ trait’ using (a) linear regression or (b) logistic regression.

%PRV is a metric measuring how much of the variation explained by a model can be attributed to non-additive terms. We calculated this by first using the fitted model in question to predict response values (survival or log final mass) across a grid of the two focal trait values, then fitting an additive model to those predictions, with the trait values treated as factors. This new additive model should perfectly fit any variation in the response that can be attributed to additive effects (e.g. will exactly fit data predicted from a linear model without an interaction term); any residuals of this new additive model are therefore due to non-additivity. We calculate %PRV from a type III ANOVA of the additive model: %PRV is the ratio of the residual sum of squares to the total sum of squares, multiplied by 100 to convert to percentages. This makes %PRV somewhat analogous to the coefficient of determination (R^2^) converted to percentages: a value of 50% means that half of the variation in model predictions is due to non-additive model components. To avoid %PRV depending on regions of extrapolation, we used a weighted ANOVA with weights corresponding to nearness to actual data points, calculated using multivariate kernel density estimation.

PRR and %PRV have natural units that are comparable between models: range in the response (log weight or survival) and percentages, respectively. Because Random Forests are more flexible than regression models, we expect the non-interactive components of our Random Forest decomposition (see below) to explain a larger fraction of the variation in the data than would our regression models, resulting in a lower %PRV for Random Forest analyses than regression models.

### Random Forest

Like many machine learning algorithms, Random Forests can be very effective at prediction (e.g., “How fast do caterpillars grow on a plant with these specific defense traits?”), but do not directly provide information (analogous to regression coefficients) about the basis for those predictions (e.g. “How much does each plant trait affect caterpillar growth rate?”). Hooker (2007) provides methodology for interrogating a fitted Random Forest model to determine the presence or absence of interactions among variables, and to quantify the importance of interactions if they are present. For a given experiment and response (growth or survival), we first fit a Random Forest model that included all available plant traits (Random Forests capture interactions without needing explicit terms to represent them). Following Hooker (2007), we then treated this fitted model as if it were an experimental system, in which we could measure herbivore performance on hypothetical plants having any specified combination of defense trait values. We conducted simulated experiments, using the fitted Random Forest to generate the “response” (predicted herbivore performance) for a set of hypothetical plants with differing traits. The predicted herbivore performance was thus the result of any linear and non-linear patterns in the real data that the Random Forest model captured. We carried out simulated experiments for each pair of defense traits, and analyzed the resulting simulated data to determine if the response was additive (no interactions) or non-additive (synergy, antagonism or other relationship). For a detailed walkthrough of simulated experiments and their analysis, see Appendix S1 (annotated code for implementing this technique is available on Figshare).

### Simulated synergies and antagonisms

Because Random Forest models are equation-free, the method presented above for using random-forest models can capture non-bilinear synergies (or antagonisms), but the fitted model does not provide information for the functional form of that relationship, and neither do PRR or %PRV. To determine whether Random Forest model fits with non-additive relationships corresponded to synergies or antagonisms, we simulated data as if it had been generated by the Random Forest analysis procedure above, but with known functional forms of synergies or antagonisms (see Appendix S1 for details). As in the methodology above, we fit this simulated data with an additive linear model. Synergies and antagonisms represent non-additive processes, and might be identifiable from the residuals of this model; we plotted those residuals (the third column in Fig 2a-f). We explored many potential trait synergies and antagonisms, but here present results for three synergies (linear, threshold, diminishing marginal returns) and three antagonisms (linear, threshold, decaying).

**Figure 2:**
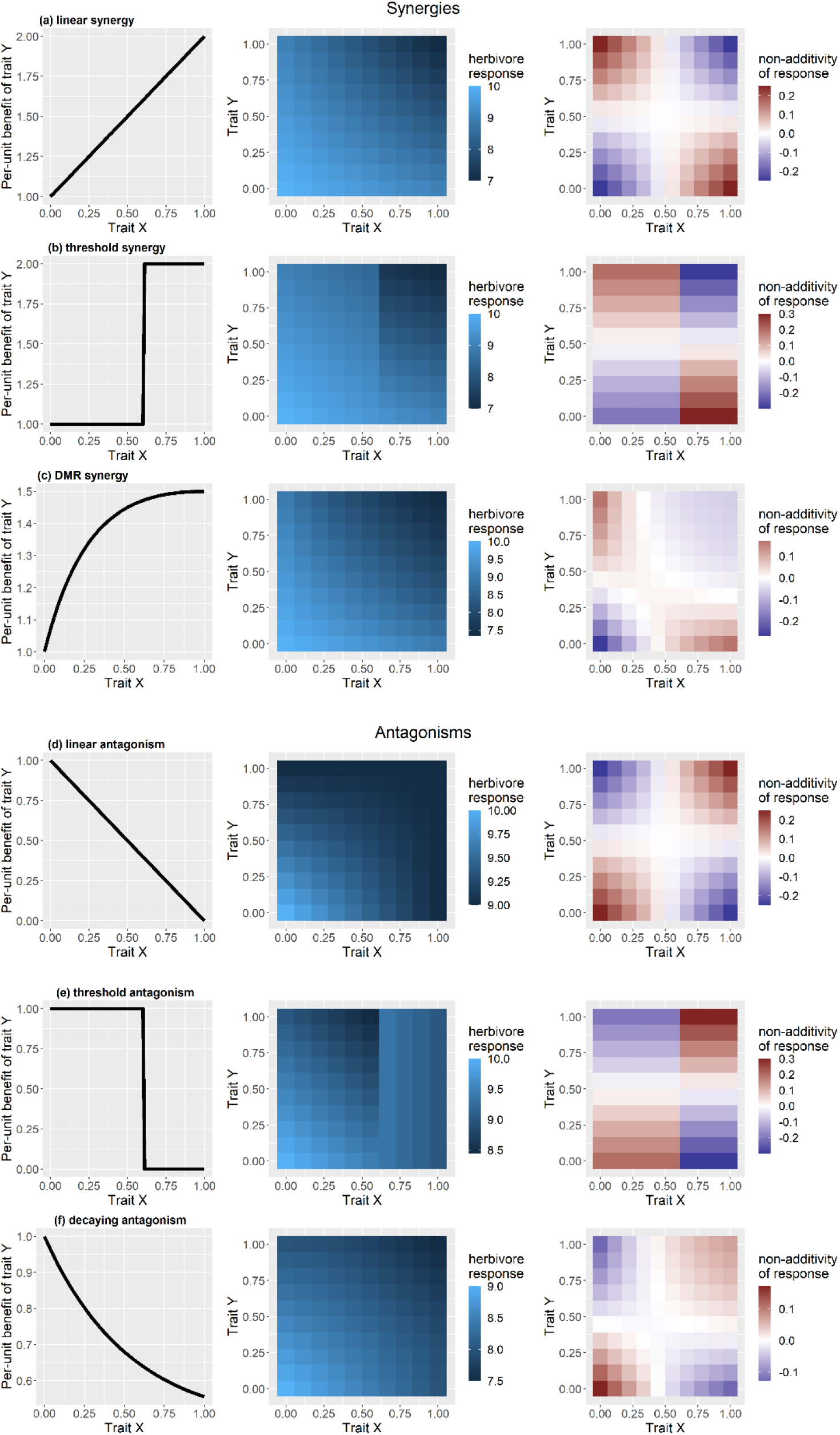
Data simulated from known synergies or antagonisms. For each relationship, we first show the per-unit benefit of investment in trait Y (when Y is held at 0.5) as trait X is varied (left); then herbivore performance, which could represent growth or survival depending on units and rescaling (middle); then the residuals of herbivore performance when fit with an additive model. In the residual plots, positive values show herbivores over-performing the additive model, and negative values show herbivores underperforming the additive model. (a-c) for synergistic relationships, herbivores under-perform the additive model when both defense traits are high or both are low, and over-perform when one trait is high and the other is low. (d-f) for antagonistic relationships, herbivores over-perform the additive model when both traits are high or both are low, and under-perform when one trait is high and the other is low.

### Classifying random forest results

Based on our simulations, we found that synergies and antagonisms had residuals that could be separated into four quadrants based on the sign of the residuals (see Fig. 2). For simplicity, we label the four quadrants Q1, Q2, Q3, and Q4 starting in the bottom left (nearest the origin) and moving clockwise. We found that synergies were represented by herbivores under-performing the additive model (negative residuals) in Q1 and Q3 (the quadrants on the diagonal), and over-performing the additive model (positive residuals) in Q2 and Q4 (the quadrants on the off-diagonal). The reverse was true for antagonisms. To quantify how much a given trait pair matched one of these qualitative patterns, we define a new test statistic, θ. θ was calculated as the sum of the signs of residuals in Q1 and Q3 minus the sum of the signs of residuals in Q2 and Q4 divided by the total number of predictions summed. Residuals of 0 were given a sign of 0. For any pair of breakpoints separating the trait space into four quadrants, θ represented the average tendency towards synergy (negative values) or antagonism (positive values), with θ = −1 if each prediction followed the described pattern for synergies, θ = 1 if each prediction followed the pattern for antagonisms, and 0 if residuals were evenly split between those patterns. For each fitted random forest model, we chose the breakpoints that maximized the absolute value of θ, e.g. the breakpoints that best represented a synergy or antagonism. To classify an interaction as a synergy or antagonism, we simulated 999 permutations of the Random Forest predictions, in which the predictions of each grid point were randomly re-assigned. For each of these simulated data sets, we then identified the new optimal breakpoints and calculated θ, generating the null distribution of θ for that trait pair. We designated an interaction as “unclear” if the observed θ was between the 0.025 and 0.975 quantiles of the null distribution, and “synergy” or “antagonism” if it was below or above those thresholds, respectively.

### Software and Packages

All analyses and simulations were carried out in the programming language R (R Core Team 2021). We used the following key packages: **mgcv** (Wood 2011) to fit mixed effects models; **randomForest** (Liaw and Wiener 2002) to fit Random Forest models; **car** (Fox and Weisberg 2019) for marginal hypothesis testing and calculating sum of squares; the **tidyverse** package suite (Wickham et al. 2019) for data cleaning and manipulation; **ggplot2** (Wickham 2016), **ggpubr** (Kassambara 2020), and **cowplot** (Wilke 2020) for creating figures. The scripts used to carry out all our analyses are available on Figshare (doi:10.6084/m9.figshare.20421633).

## Results

### Trait Correlations

Several patterns of trait co-expression emerged in our field studies (Figures S1, S2). In 2015, several of the physical traits showed positive phenotypic correlations – notably, leaf toughness was correlated with both latex exudation from leaves (r = 0.17, p < 0.021) and leaf trichome density (r = 0.49, p < 0.001). Percent leaf carbon was positively correlated with the two traits that clearly incorporated carbon, trichome density (r = 0.49, p < 0.001) and leaf toughness (r =0.21, p<0.01). In 2016, physical defense traits (trichomes, toughness, and latex) all had significant positive correlation with each other (latex and toughness, r = 0.38, p < 0.01; latex and trichomes, r = 0.13, p <0.04; toughness and trichomes, r = 0.30, p < 0.01), and were generally negatively correlated or essentially uncorrelated with cardenolides.

### Linear regression

For suppressing monarch growth, we found strong evidence for a bilinear synergy between non-nitrogen and LMA; moderate evidence for a bilinear synergy between leaf toughness and carbon, and between non-nitrogen and trichome density; and weak evidence for a bilinear synergy between leaf toughness and trichome density, and between leaf toughness and non-nitrogen (Table 1). For suppressing monarch growth, we also found moderate evidence for a bilinear antagonism between cardenolide 18.6 and non-nitrogen, between cardenolide 18.6 and leaf toughness, and between cardenolide 18.6 and LMA. For suppressing monarch survival, we found weak evidence for a synergy between cardenolide 18.6 and latex. For suppressing *L. clivicollis* growth, we found weak evidence for a synergy between latex and cardenolide 10.2, and between latex and leaf toughness. We also found weak evidence for an antagonism between cardenolide 8.4 and the covarying cardenolide suite 2016. For suppressing *L. clivollis* survival, we found weak evidence for a synergy between latex and cardenolide 10.2.

**Table 1:** Summary of regression analysis for bilinear interactions. LMA is leaf mass per area, the inverse of specific leaf area. All trait pairs with P≤0.1 are shown; lower P values represent synergies or antagonisms with stronger support. F-statistic and chi-square statistic are for a single model term with one degree of freedom; relationship is identified by the sign of the interaction coefficient (negative = synergy). %PRV measures the importance of non-additivity in explaining model fit (see methods); PRR (“Predictable Response Range”) measures the range of response values predicted by the model (see methods). For growth, PRR is in units of log(grams final weight), and for survival, PRR is the expected probabilities of herbivore survival.

| data set | trait 1 | trait 2 | relationship | F or chi-square | P | %PRV | PRR |
| --- | --- | --- | --- | --- | --- | --- | --- |
| Beetle growth | latex | cardenolide 10.2 | synergy | 3.9439 | 0.051 | 27 | -6.71 -- -5.92 |
|  | latex | leaf toughness | synergy | 3.836 | 0.054 | 96 | -7 -- -5.7 |
|  | cardenolide suite 2016 | cardenolide 8.4 | antagonism | 2.8265 | 0.096 | 60 | -6.69 -- -5.67 |
| Beetle survival | latex | cardenolide 10.2 | synergy | 2.6981 | 0.1 | 20 | 0 -- 0.84 |
| Monarch growth | non-nitrogen | LMA | synergy | 10.3651 | 0.002 | 53 | -5.77 -- -4.51 |
|  | non-nitrogen | cardenolide 18.6 | antagonism | 5.7922 | 0.018 | 84 | -5.62 -- -4.58 |
|  | leaf toughness | carbon | synergy | 5.2086 | 0.025 | 61 | -5.84 -- -4.57 |
|  | leaf toughness | cardenolide 18.6 | antagonism | 5.187 | 0.025 | 98 | -5.39 -- -4.58 |
|  | cardenolide 18.6 | LMA | antagonism | 4.9287 | 0.029 | 75 | -5.52 -- -4.58 |
|  | trichome density | non-nitrogen | synergy | 4.2549 | 0.042 | 34 | -5.37 -- -4.38 |
|  | leaf toughness | trichome density | synergy | 3.5435 | 0.063 | 34 | -5.46 -- -4.5 |
|  | leaf toughness | non-nitrogen | synergy | 3.0646 | 0.083 | 61 | -5.32 -- -4.7 |
| Monarch survival | latex | cardenolide 18.6 | synergy | 3.1335 | 0.077 | 82 | 0.66 -- 1 |

All of our presumed defense traits showed context-dependent effects. Of the 204 interaction pairs, 93 (46%) had switching points that were within one standard deviation of the mean of the second trait (e.g., Figure 3) (Table S1). Only 75 of the pairs (37%) had switching points more than two standard deviation from the mean. Moreover, of all the traits measured, each of them were context-dependent rather than consistently defensive (or consistently beneficial) to herbivores in at least one trait pairing for either growth or survival (Table 2).

**Figure 3:**
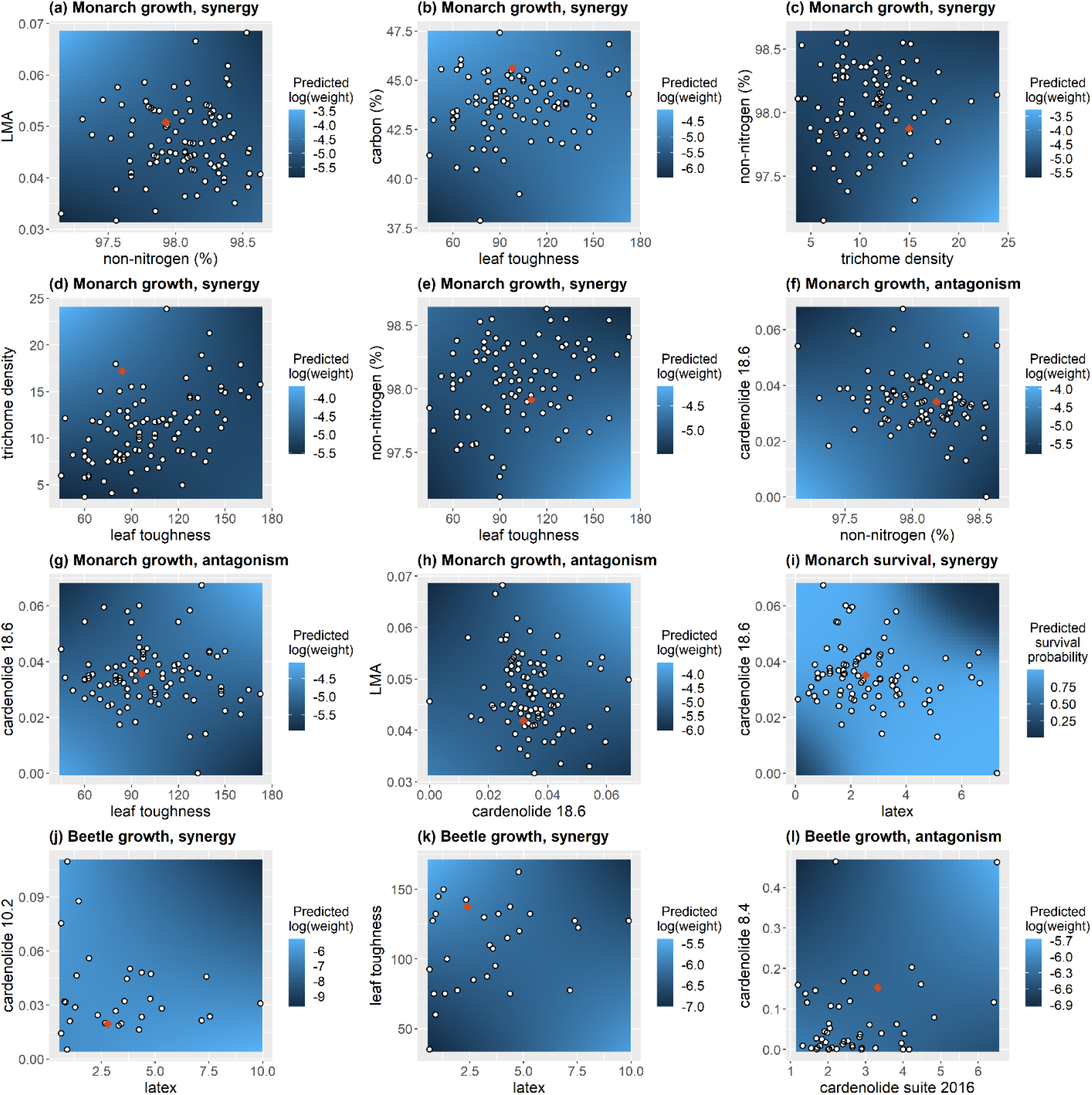
Bilinear synergies fit to observed data using regression models, for all interaction terms with p<0.1 (see Table 1). Heat maps show predicted survival or growth from fitted models (analogous to central column of Figure 2); points correspond to trait values of observed plants. Orange diamonds show the “switching points” where the effect on herbivores of increasing a plant trait changes from being helpful to harmful or vice versa. Note that because the regression models are parametric, the values of the surface in regions without data are unlikely to represent meaningful patterns (i.e., extrapolation away from the data should not be trusted). Units: LMA is shown in g/10cm^2^ dry mass; carbon and non-nitrogen in percent of dry mass; leaf toughness in grams of force, trichomes in counts per millimeter of transect; cardenolides in mg/g dry mass, latex in mg fresh mass, cardenolide suites are unitless.

**Table 2:** Traits were identified as context-dependent if their effect on herbivores switched sign (positive to negative) in response to another trait changing values within one standard deviation of its mean.

| Trait | Response | Context-dependent on: |
| --- | --- | --- |
| carbon | Monarch growth | - |
| carbon | Monarch survival | cardenolide 18.6, leaf toughness, cardenolide suite 2015 |
| non-nitrogen | Monarch growth | LMA, cardenolide 10.5, cardenolide 18.6, carbon, trichome density, leaf toughness, cardenolide suite 2015 |
| non-nitrogen | Monarch survival | LMA, cardenolide 18.6 |
| cardenolide 10.2 | Beetle growth | cardenolide 8.4, leaf toughness, trichome density, latex |
| cardenolide 10.2 | Beetle survival | - |
| cardenolide 10.5 | Monarch growth | LMA, cardenolide 18.6, non-nitrogen, carbon, latex, leaf toughness, cardenolide suite 2015 |
| cardenolide 10.5 | Monarch survival | cardenolide suite 2015 |
| cardenolide 18.6 | Monarch growth | LMA, cardenolide 10.5, non-nitrogen, carbon, latex, leaf toughness, cardenolide suite 2015 |
| cardenolide 18.6 | Monarch survival | LMA, cardenolide 10.5, non- nitrogen, carbon, latex, trichome density, cardenolide suite 2015 |
| cardenolide 8.4 | Beetle growth | cardenolide 10.2, leaf toughness, trichome density |
| cardenolide 8.4 | Beetle survival | - |
| cardenolide suite 2015 | Monarch growth | non-nitrogen, leaf toughness |
| cardenolide suite 2015 | Monarch survival | cardenolide 10.5, cardenolide 18.6, carbon, latex, trichome density |
| cardenolide suite 2015 | Beetle growth | cardenolide 8.4, cardenolide 10.2, trichome density |
| cardenolide suite 2015 | Beetle survival | cardenolide 10.2 |
| latex | Monarch growth | - |
| latex | Monarch survival | LMA, cardenolide 10.5, cardenolide 18.6, carbon, trichome density, leaf toughness, cardenolide suite 2015 |
| latex | Beetle growth | cardenolide 10.2, leaf toughness, trichome density |
| latex | Beetle survival | trichome density |
| LMA | Monarch growth | cardenolide 18.6, non-nitrogen, trichome density, cardenolide suite 2015 |
| LMA | Monarch survival | cardenolide 10.5, cardenolide 18.6, non-nitrogen, carbon, latex |
| leaf toughness | Monarch growth | cardenolide 10.5, cardenolide 18.6, non-nitrogen, carbon, latex, trichome density, cardenolide suite 2015 |
| leaf toughness | Monarch survival | carbon |
| leaf toughness | Beetle growth | latex |
| leaf toughness | Beetle survival | cardenolide 10.2, cardenolide suite 2016, trichome density |
| trichome density | Monarch growth | - |
| trichome density | Monarch survival | cardenolide suite 2015 |
| trichome density | Beetle growth | cardenolide 8.4, cardenolide 10.2, cardenolide suite 2015, latex |
| trichome density | Beetle survival | cardenolide 10.2, cardenolide suite 2016, leaf toughness, latex |

### Simulations to classify Random Forests

Data simulated from synergies always produced residual plots in which the bottom left and top right regions showed herbivores underperforming compared to predictions from an additive model (an ANOVA model with no interaction terms, fit to the simulated data), and the top left and bottom right showed herbivores over-performing compared to predictions from an additive model (e.g. Fig. 2a-c). The reverse was true for antagonisms: herbivores always over-performed compared to predictions from an additive model in the bottom left and top right, and underperformed compared to predictions from an additive the top left and bottom right regions (e.g. Fig. 2d-f). Depending on the functional form of the synergy or antagonism, patterns of residuals did not always separate perfectly into four quadrants (e.g. Fig. 2c, right panel), but the general pattern held true for all synergies and antagonisms we evaluated (see code on Figshare to explore additional synergies and antagonisms).

### Random Forests

Random Forests identified many additional synergies and antagonisms that were not detected using regression approaches (recall, differences in results are not unexpected, because the regression analysis specifically tests for bilinear departures from linearity, while the Random Forests analysis tests for arbitrary deviations from additivity). Because our metrics for evaluating the non-additivity of the modeled relationship are new, there are no clear guidelines as to what constitutes strong or weak evidence. We choose to emphasize synergies and antagonisms with %PRV of 20 or higher (i.e., 20% or more of the sum of squares of the random forest model predictions cannot be explained by additive models) for which the relationship was clear (i.e., θ outside the 95% range of the null distribution). However, we provide readers with all results with clear relationships and %PRV larger than 5 in Table 3 (and all results are available in Table S2). All 10 synergies and antagonisms with %PRV of 20 or greater were detected for Monarch survival. The relationship with the highest %PRV (33%) was a synergy between latex and trichome density, with expected probability of survival ranging between 0.68 and 0.93 across observed trait values (PRR). Of the 10 interactions with %PRV 20 or higher, physical traits of leaves – trichome density, leaf toughness, and leaf mass area – were present in all but one, half of the interactions involved one of the two nutritional quality traits (carbon or non-nitrogen), and three of the five synergies involved one cardenolide compound, cardenolide 10.5.

**Table 3:** Summary of Random Forest analysis, showing all trait pairs with %PRV of 5 or higher (after rounding) with clear synergies or antagonisms (θ outside the 95% limits of the null distribution from permutation testing). %PRV and PRR are the same metrics as in Table 1.

| data set | trait 1 | trait 2 | relationship | %PRV | PRR |
| --- | --- | --- | --- | --- | --- |
| Monarch survival | latex | trichome density | synergy | 33 | 0.68 -- 0.93 |
| Monarch survival | LMA | cardenolide 10.5 | synergy | 26 | 0.78 -- 0.94 |
| Monarch survival | leaf toughness | trichome density | antagonism | 26 | 0.7 -- 0.93 |
| Monarch survival | cardenolide 10.5 | carbon | synergy | 25 | 0.8 -- 0.93 |
| Monarch survival | trichome density | cardenolide 10.5 | synergy | 24 | 0.66 -- 0.93 |
| Monarch survival | trichome density | carbon | antagonism | 23 | 0.7 -- 0.92 |
| Monarch survival | latex | leaf toughness | antagonism | 22 | 0.69 -- 0.94 |
| Monarch survival | LMA | carbon | antagonism | 21 | 0.79 -- 0.92 |
| Monarch survival | LMA | non-nitrogen | antagonism | 20 | 0.7 -- 0.93 |
| Monarch survival | trichome density | non-nitrogen | synergy | 20 | 0.66 -- 0.93 |
| Monarch survival | leaf toughness | carbon | antagonism | 19 | 0.76 -- 0.93 |
| Monarch survival | leaf toughness | cardenolide 10.5 | synergy | 18 | 0.78 -- 0.94 |
| Monarch growth | leaf toughness | carbon | synergy | 18 | -5.37 -- -4.9 |
| Monarch survival | non-nitrogen | carbon | synergy | 17 | 0.74 -- 0.93 |
| Monarch survival | LMA | leaf toughness | antagonism | 16 | 0.73 -- 0.93 |
| Monarch survival | latex | cardenolide suite 2015 | synergy | 16 | 0.69 -- 0.93 |
| Monarch survival | latex | cardenolide 18.6 | synergy | 16 | 0.7 -- 0.93 |
| Monarch survival | cardenolide 10.5 | cardenolide suite 2016 | antagonism | 15 | 0.77 -- 0.93 |
| Monarch survival | cardenolide 18.6 | carbon | antagonism | 13 | 0.8 -- 0.92 |
| Monarch growth | cardenolide suite 2015 | carbon | antagonism | 11 | -5.33 -- -4.9 |
| Monarch survival | cardenolide 18.6 | non-nitrogen | synergy | 9 | 0.71 -- 0.92 |
| Monarch growth | trichome density | carbon | antagonism | 9 | -5.39 -- -4.88 |
| Monarch growth | non-nitrogen | carbon | synergy | 8 | -5.3 -- -4.9 |
| Monarch growth | LMA | non-nitrogen | synergy | 8 | -5.31 -- -4.95 |
| Beetle growth | leaf toughness | cardenolide 10.2 | synergy | 8 | -6.51 -- -5.92 |
| Monarch growth | LMA | carbon | antagonism | 7 | -5.35 -- -4.88 |
| Beetle growth | leaf toughness | cardenolide suite 2016 | synergy | 7 | -6.57 -- -6.03 |
| Beetle growth | cardenolide 10.2 | cardenolide suite 2016 | synergy | 6 | -6.59 -- -6.06 |

### Correspondence between regression and Random Forest results

Eight of the thirteen bilinear synergies and antagonisms identified using regression were similarly classified as synergy or antagonism using Random Forests (Table S2). In four cases, regression identified bilinear interactions that our Random Forest method did not clearly identify as either synergy or antagonism. For the interaction between cardenolide 8.4 and the cardenolide 2016 suite on *L. clivicollis* growth, the “unclear” designation derives from the poor statistical power (only 4 of the 100 grid cells in trait space were usable) (Figure S3). The other three interactions (latex by leaf toughness on *L. clivicollis* growth, cardenolide 18.6 by leaf toughness on monarch growth, non-nitrogen by leaf toughness on monarch growth) likely reflect complicated functional forms that did not fall clearly into synergies or antagonisms using random forests, but were identifiably synergy or antagonism when simplified to bilinear interactions. Of the bilinear synergies identified using linear regression, only the leaf toughness by trichome density interaction on monarch growth conflicted with Random Forest results (regression: weak evidence for a synergy; Random Forest: antagonism with %PRV = 3). Of the synergies identified using Random Forests, none of the 10 interactions with %PRV 20 or higher had any evidence of bilinear synergy (P > 0.1 in all cases). In fact, of the 37 interactions Random Forest analyzed with a %PRV of 5 or greater, only four interactions (synergies) had P values less than 0.1. In general, %PRV was much higher for regression models than for Random Forests. This was true for all bilinear interactions with P < 0.1, 65% of interactions with Random Forest %PRV greater than or equal to 5, and 70% of all interactions.

## Discussion

Using surveys of herbivore performance in the field and both well-established and novel statistical tools, we found evidence for the plant defense synergy hypothesis. We found the first direct empirical support for previously hypothesized synergies between latex and cardenolides (Figure 3i, j), and between latex and trichomes (Figure 4a) on specialist monarch survival. In comparing the bilinear synergies and antagonisms identified using regression with the synergies and antagonisms identified using Random Forests, we found both overlapping and unique results, suggesting these tools are complimentary rather than redundant. When regression analysis identified a significant bilinear synergy or antagonism, the Random Forest analysis usually agreed with the type of interaction. However, the strongest synergies and antagonisms identified by the Random Forest analysis did not have evidence for bilinear interactions. Using only regression analysis, we would have identified a single synergy for monarch survival (latex by cardenolide 18.5, a previously hypothesized synergy). However, by applying our Random Forest methods, we identified an additional five notable synergies affecting monarch survival (%PRV greater than or equal to 20), including the latex by trichome density synergy previously hypothesized based on the natural history of *D. plexippus* caterpillars (Zalucki et al. 2002). In contrast, many of the bilinear relationships identified using regression did not have clear patterns or had small %PRV when examined using Random Forests. This reflects the different strengths of these two methods; the relative simplicity of regressions provides more power when relationships are largely bilinear, allowing us to detect such relationships more easily. In contrast, Random Forests are by their nature flexible, and require more data to confidently describe a relationship form; however, their flexibility allows them to detect relationships that do not conform to the bilinear forms assumed by regression models.

**Figure 4:**
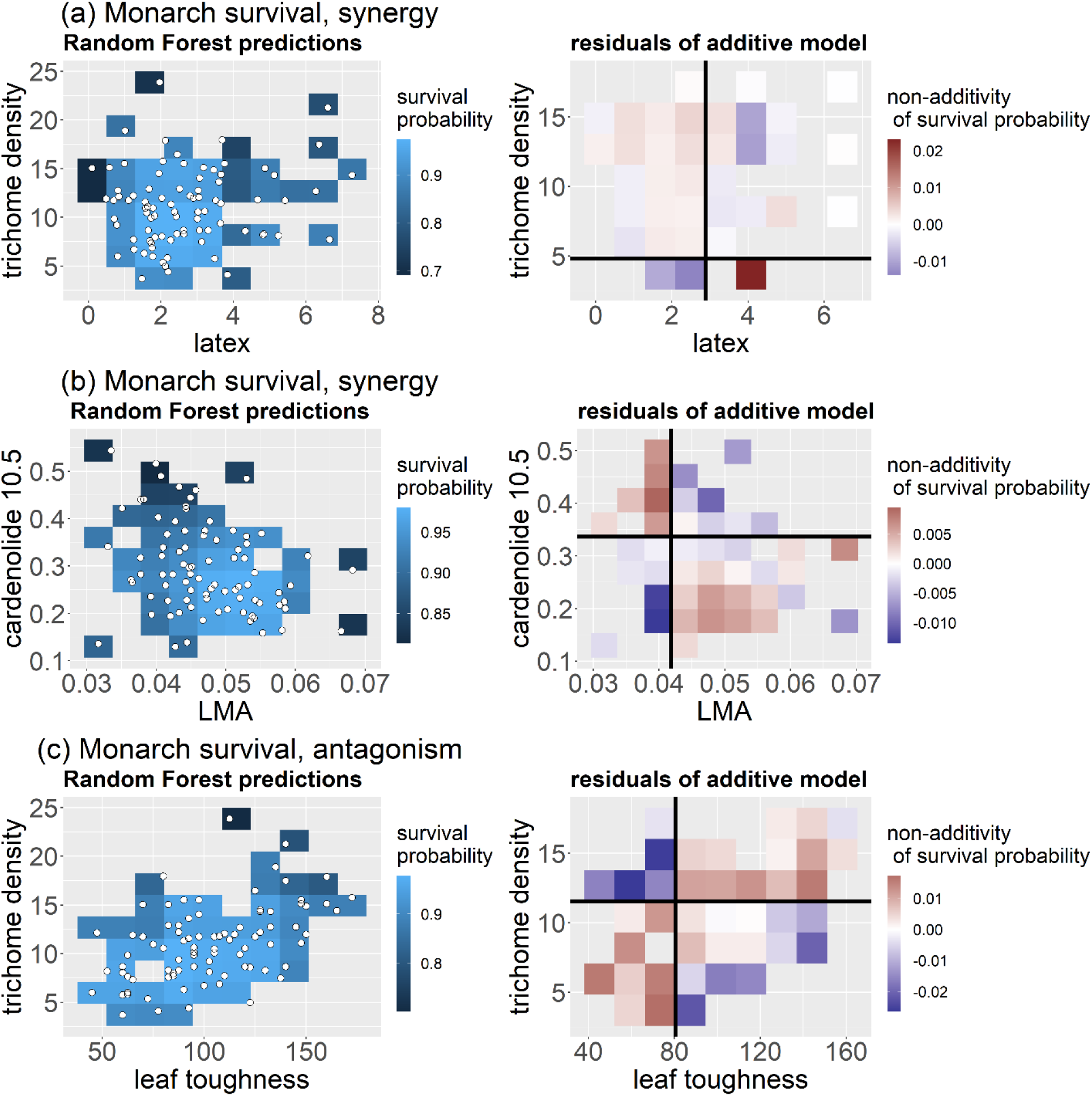
Plots of the three synergies and antagonisms identified from Random Forest analyses that showed the highest %PRV; all happen to be from the ‘Monarch survival data. On the left, predictions of the random forest for pairs of trait values calculated as described in the methods (analogous to the second column of Figure 2). White points represent actual data; only grid cells containing actual data are plotted. On the right, residuals from fitting an additive model to the predictions on the left (analogous to the third column of Figure 2). Black lines identify the quadrant separation that maximizes the difference in residual between Q1 + Q3 (bottom left and top right quadrants) vs Q2 and Q4 (top left and bottom right quadrants) (i.e. the separation that maximizes the absolute value of θ). Note that because additive models will perfectly fit grid cells that are alone in their row or column (necessarily leading to residuals of 0), we have removed these entries from the plots. In comparing to Figure 2, we can see that the residuals of (a) and (c) are consistent with synergies, and the residuals of (b) are consistent with antagonisms (permutation testing confirms this). Similar plots for every trait pair of each data set can be found in the supplements. (a) θ = −0.50, %PRV = 33, PRR = 0.683 -- 0.934, (b) θ = 0.7, %PRV = 25, PRR = 0.696 – 0.928, (c) θ = −0.54, %PRV = 25, PRR = 0.797 – 0.934. Units are as described in Figure 3.

We found evidence for a range of new potential synergies and antagonisms. The strongest evidence was for a bilinear synergy between non-nitrogen and leaf mass per area (LMA) on monarch growth (Figure 3a): monarch growth was worse on thick, low-nitrogen leaves than the leaf thickness or low nitrogen alone would imply. For low values of LMA (thin leaves), increased non-nitrogen led to *increased* herbivore growth; similarly, for low values of non-nitrogen, increased LMA increased herbivore growth. However at higher levels of either trait – past the switching point – increasing the level of the other trait led to decreased herbivore performance. Lack of nitrogen and high LMA may thus jointly contribute to low nutritional value for insect herbivores (Wright and Cannon 2001). Indeed, comparisons across milkweed species previously suggested low nutritional quality as a convergent defense strategy employed in this genus (Agrawal and Fishbein 2006). In addition to potentially shaping defense strategies on its own, this synergy between non-nitrogen and LMA reinforces a pattern of covariation between LMA and non-nitrogen associated with the leaf economics spectrum (Wright et al. 2004, Fajardo and Siefert 2018); these trait correlations have been explained by physiological costs and benefits associated with resource acquisition and leaf lifespan (Shipley et al. 2006). Previous work in *A. syriaca* shows this relationship among wild plants in the field, and among genotypes grown in a common environment (Agrawal 2020). These findings suggest that the co-expression of LMA and low leaf nitrogen identified as part of the leaf economic spectrum coincides with a defense synergy: nutrient poor plants suppress herbivore growth even more by increasing LMA than do plants with higher nitrogen content. This inference must be balanced, however, by our finding that there was generally an antagonism between LMA and non-nitrogen on monarch survival; a detailed model would be needed to integrate the combined effects of leaf economics, herbivore growth, and herbivore survival.

We were surprised to find that our analysis identified nearly every “defensive” trait as context-dependent (Table 2, Figure 3). Our regression models predicted that the effect of increasing any given trait could be helpful or harmful to herbivores depending on the expressed levels of other traits (i.e., the switching point in the LMA by non-nitrogen synergy described above). This context-dependence is consistent with the adaptive redundancy hypothesis (Rasmann and Agrawal 2009), and may help to explain why studies often find inconsistent benefits of defensive traits (Agrawal 2011). For example, while cardenolides and latex are generally correlated with resistance to caterpillars, sometimes they have no apparent effect (Agrawal 2005, Agrawal and Fishbein 2006). While there are many reasons that experiments might find different or even conflicting results, our study suggests that the effect of a focal “defense” trait could differ or even reverse if another key (potentially unmeasured) trait differed sufficiently in expression between two experiments. Such variation in interacting traits is likely, as studies have found that expression levels of individual defense traits can vary substantially based on environmental conditions. For example, latex can differ in expression based on water availability and plant ontogeny (Agrawal et al. 2014, Barton 2014), and prickle and trichome density can vary based on light and water availability (Ehleringer 1982, Barton 2014, Agrawal et al. 2012). Taken together, this suggests plants face a puzzle of interacting traits whose benefits all depend on one another, with environmental conditions impacting or constraining some of those traits.

The herbivores species used in our experiments were both *Asclepias* specialists, and have adaptations that partially mitigate many of the defenses of milkweed plants (e.g. Dobler et al. 2012, Zalucki et al. 2001, Agrawal 2005, Tao et al. 2016, Jones et al. 2019). As such, low nutrient availability is expected to play a large role in determining herbivore performance (Feeny 1976), and has been proposed as a strategy for *Asclepias* specifically (Agrawal and Fishbein 2006). Three of the bilinear synergies identified by regression analysis and two of the notable interactions identified by Random Forest analysis (one synergy, one antagonism) involve non-nitrogen. These synergies and antagonisms involving non-nitrogen may reflect the expected importance of low nutrition as a defensive trait. Equivalently, these interactions between non-nitrogen and other defense traits suggest that the effect of those other traits – trichome density, LMA, cardenolides, leaf toughness – vary based on the resources available to the herbivore. Just as the effects of environmental stresses on plants can vary substantially based on resources available to those plants (e.g. Eneji et al. 2008, Zahoor et al. 2017), we expect the effects of plant defense traits to be different if the herbivore is limited by nitrogen, or has it in relative excess.

Several of the identified synergies involved latex: latex by various cardenolides, latex by leaf toughness, latex by trichome density, and latex by LMA. Latex is present in nearly 10% of angiosperm species, and has no known function in plants other than defense (Agrawal and Konno, 2009). Latex can be quite harmful to herbivores directly (e.g. Zalucki & Brower 1992, van Zandt and Agrawal 2004), but its commonness across plant families is all the more understandable if it synergizes with other defenses. Here our measure of latex was the wet mass exuded upon leaf damage, which in *Asclepias* typically correlates negatively with insect performance. Particularly notable were the latex by cardenolide synergies identified in this study, which have been proposed but untested in the past (Agrawal 2011). The hypothesized mechanisms for latex by cardenolide synergies – that latex could serve as a delivery system for toxic phytochemicals, and toxic phytochemicals make latex even more dangerous to herbivores – is not a special feature of monarchs or *L. clivicollis* feeding on *A. syriaca.* The latex of most plants contains secondary metabolites, often at elevated levels compared to other plant tissues (Agrawal and Konno 2009). Further, a recent study of the common dandelion *(Taraxacum officinale)* found that herbivore damage increased when sesquiterpene levels in latex were reduced; while this study was not explicitly addressing synergies, their findings imply latex by sesquiterpene synergy (Huber et al. 2016). To our knowledge, our study is the first to find explicit evidence for latex by secondary metabolite synergies, but we speculate that such synergies may be common across plant taxa.

Secondary metabolites are often emphasized in studies of plant defenses because of their generally-unambiguous function in resistance to herbivores (Ehrlich and Raven 1964). We found numerous synergies and some antagonisms between cardenolides and other defensive traits in *A. Syriaca;* this likely reflects the importance of cardenolides in resisting milkweed herbivores (Zalucki 2001, Tao et al. 2016, Agrawal et al. 2021). However, a meta-analysis of plant traits and herbivore resistance found that physical resistance traits were more reliably correlated with herbivore resistance, particularly when focusing on specialist herbivores (Carmona et al. 2013). The three purely physical resistance traits represented in our study (LMA, leaf toughness, trichome density) were present in 60% of the bilinear interactions and 90% of the Random Forest interactions. The importance of physical resistance traits in plant defenses across species, and the prevalence of synergies and antagonisms between physical traits and other defenses in *A. Syriaca* suggests that the suites of traits employed by individual plants may be structured by the costs and indirect benefits of physical traits.

We find PRR to be a useful tool for examining the effects of trait variation on response, measured in meaningful units. We caution that this metric should only be used for data in which variation in traits comes from natural sources (e.g., observational data). In a controlled experiment in which we could manipulate the expression levels of traits, PRR could generally be inflated by increasing the range of trait values outside any biologically meaningful region. %PRV is a valuable tool for interpreting the role of non-additivity in model fits (and is a necessary step for our use of Random Forests), but model predictions should be weighed by their proximity to actual data points to avoid %PRV values being driven by extrapolation.

Multivariate plant defense strategies have been explored through observations of trait co-expression. This has been done at the level of between-species comparisons (e.g., Agrawal and Fishbein 2006, da Silva and Batalha 2011, Moles et al. 2013, Sanczuk et al. 2020), between-population comparisons (e.g., Barton 2014), or between individuals within a population (e.g., Agrawal et al. 2014). Trait co-expression can provide indirect support for synergies, but these same signals might represent other forms of advantage or disadvantage for specific combinations. Among *Asclepias* species, comparative work has found interspecific patterns of co-expression between latex and trichomes and between LMA and non-nitrogen, but not between latex and cardenolide expression (Agrawal and Fishbein 2006). The statistical framework we provide here helps explicitly identify traits that are associated with synergistic (or antagonistic) impacts on herbivores.

Tests of synergies between classes of defenses (rather than between secondary metabolites within a class) are rare, but ours is not the first. There have been a few studies in aquatic systems finding synergies between structural components (calcium carbonate, spicules) and secondary metabolites in artificial diets simulating seaweeds (Hay et al. 1994) and sponges (Hill et al. 2005, Jones et al. 2005). Simpson and Raubenheimer (2001) found that tannic acids had an effect on locust only for some levels of protein-to-carbohydrate ratios in the artificial diets, suggesting that plant digestion inhibitors might synergize with defense strategies of low nutrient availability. Perhaps the strongest test for cross-class synergies in terrestrial plants used gene knockouts in *Nicotiana attenuata* to show that protease inhibitors (TPI, a form of digestion inhibitor) synergized with toxic nicotine to defend against generalist *Spodoptera exigua* (Stepphun and Baldwin 2007). Follow-up work in the same system also found that TPI reduced caterpillar response to simulated predation, suggesting a potential defense synergy between parasitoid-attracting volatiles and TPI. However, field experiments found plants with higher TPI actually had *reduced* parasitism (Schuman et al. 2012). Similar to Stepphun and Baldwin (2007), Green et al. (2001) found phytic acid (an anti-nutritive defense) reduced insect metabolism of defensive xanthotoxins, implying a potential synergy with phytic acid interfering with specialist herbivore counter-defenses. However, this relationship was not extended to measure the ultimate effect of the phytic acid by xanthotoxin interaction on herbivore survival or growth.

The rarity of synergy tests between classes of defenses is likely due in part to the logistical challenges involved in such experiments. The overwhelming majority of synergy studies to date have used artificial diets (e.g., Whitehead et al. 2021), and non-metabolite defenses like latex, leaf toughness, and trichomes do not lend themselves to such controlled experiments. Similarly, the most elegant current approaches for testing phytochemical synergies identify an interaction index based on the dose necessary to cause 50% mortality (ED50) (Nelson and Kursar 1999, Tallarida 2000, Richards et al. 2016). These tools are well-suited to artificial diet experiments in which an ED50 is likely to exist, but are difficult to implement when defenses cannot be manipulated and even high levels of putatively defensive traits have no guarantee of causing mortality. The synergies and antagonisms between classes of traits identified here and in the few other studies described above suggest that such trait interactions are common, even if the study of them is not. We hope the analytical tools provided here will support a broader exploration of synergies in the study of plant defenses. The benefits of being able to conduct such rigorous tests under natural conditions make this approach an essential complement to more controlled and mechanistic studies.

## Acknowledgements

We thank Tobias Züst and Ron White for their assistance with liquid chromotography, Kim Sparks and the Cornell Stable Isotope Laboratory for running carbon and nitrogen samples, Eamon Patrick and Patty Jones for maintaining herbivore colonies and providing eggs, Amy Hastings for invaluable direction and assistance in the lab, Giles Hooker for guidance in implementing his innovative approaches with Random Forests, Bob Reed and Karin van der Burg for access to and assistance with their microscope, RoseBarb Farms for the use of their land in 2015, Cornell Natural Areas and the Cornell Botanical Gardens for use and maintenance of Dunlop Meadow, and Monica Geber and Wesley Hochachka for their helpful feedback on this work. This paper is based on a dissertation presented to Cornell University in partial fulfillment of the requirements for a PhD and was funded in part by NSF-IOS 1907491 and 2209762 to AAA and a Cornell Mellon grant to CBE. CBE was supported by the National Science Foundation Graduate Research Fellowship under Grant No. 14590.

## Supplemental Tables

**Table S1:** The switching point of all trait pairs. “Switching point” is distance (absolute value) in standard deviations from the mean of the “context trait” where, when the context trait is at that value, the “focal trait” changes from being harmful to helpful to herbivores (or vice versa). If the switching point is within one standard deviation (e.g. that column is less than 1), the trait pair is shown in Table 2.

**Table S2**: Analysis results for each trait pair of each experiment, including results from both the regression and Random Forest analyses. Test statistic (P or chi-square) and P value are for regression analysis only; for all other metrics (relationship, %PRV, PRR), results for regression and Random Forest analysis are shown side by side.

---

**Figure S1:**
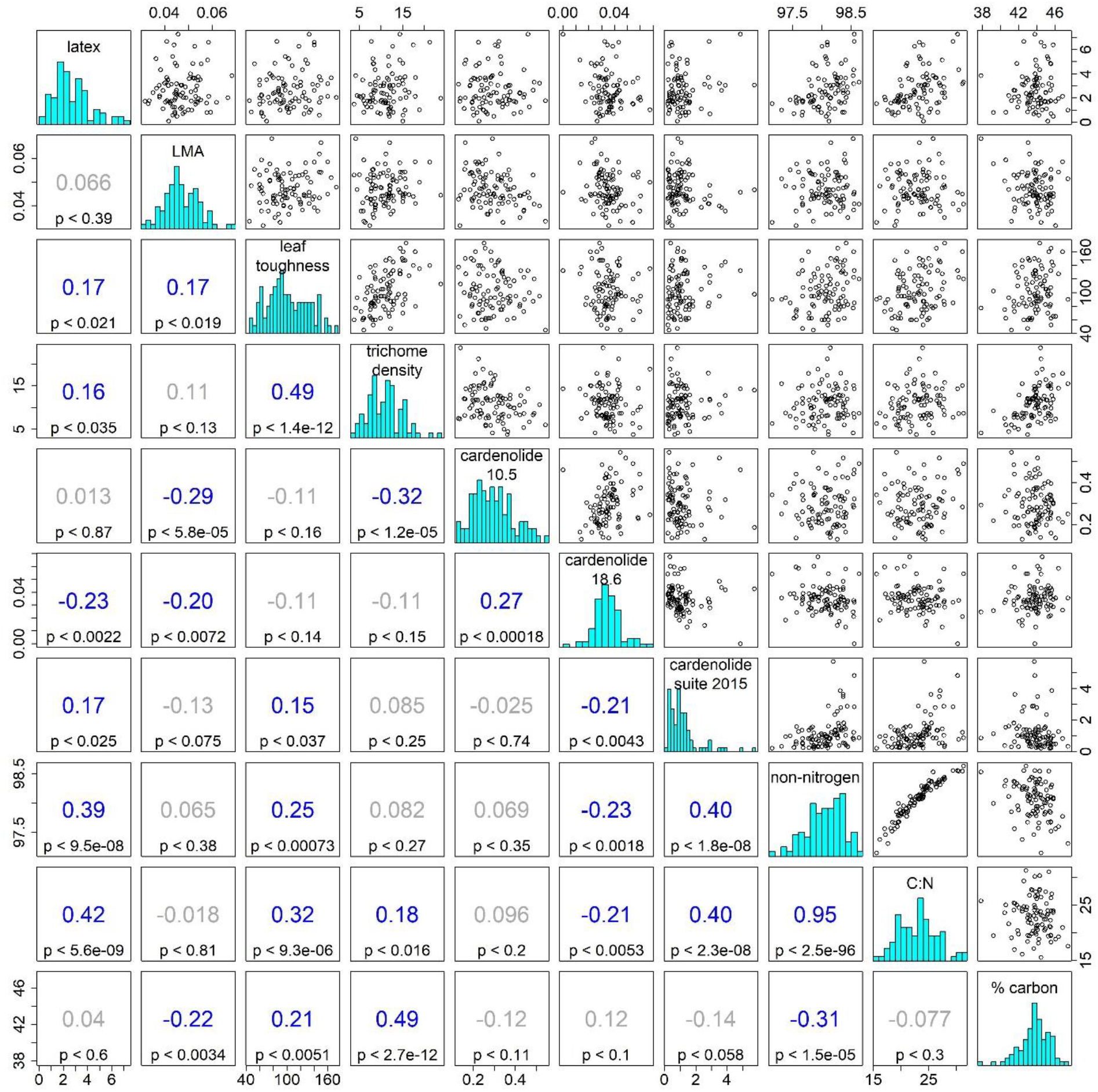
Pairwise correlations plots of plant traits for Monarch data. Upper triangle plots show observations, lower triangle shows Pearson correlation coefficients and associated p values; correlations are shown in blue when the p values are less than 0.05. On the diagonal, histograms of observed values.

**Figure S2:**
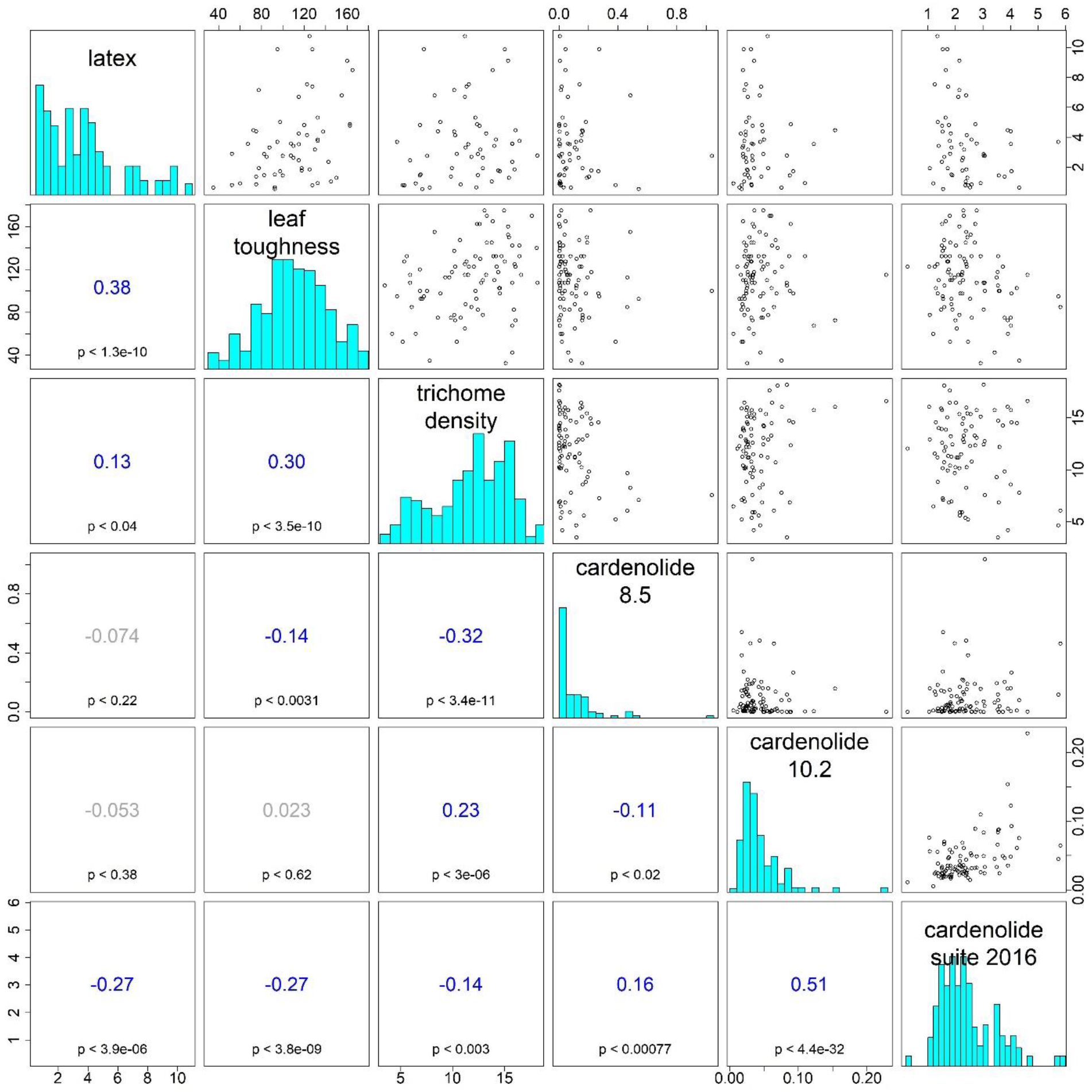
As Figure S1, but for Beetle data.

**Figure S3**: As Figure 4, but for all trait pairs. On right panels, classification was determined by permutation testing, with the more granular information on the next row: theta and the null 95% giving the test statistic and the 95% confidence limits of the simulated distribution of that test statistic under the null hypothesis.

## Notes

### Competing Interest Statement

The authors have declared no competing interest.

### Summary of Updates

We have substantially rewritten this manuscript and refined our statistical approaches. We have dropped the 2005 data and simplified our analyses into just "regression" and "random forest" rather than distinguishing between hypothesis testing and exploration. In the process of revising this manuscript, we realized that our regression model structure contained redundant random effects, as plant ID was nested within date. Further, we were correcting for variation in duration (6 or 7 days) in three places: duration was captured by plant ID, but we were also including date, and also using as our growth response the log final weight divided by days (which implicitly accounts for duration under the assumption of exponential growth). We have revised our analysis to only include the single random effect of plant ID, which can capture un-modeled variation between plants in addition to the variation associated with date and the effects of variable duration. We also now simply use log final weight as our response variable for herbivore growth, avoiding an assumed functional form of growth rate in favor of allowing our data to identify this through random effects. This new model structure is cleaner, removing some assumptions and model complexity compared to the original version. We recognized that given the construction of our analysis, traits could in fact switch from being defensive to being beneficial to herbivores if the interaction term was strong relative to the main effects. We have included this as a new part of our analysis, as we found that all of our traits were in fact context-dependent, and we believe this provides some explanation for the inconsistency of defense traits that has been previously reported in the plant defense literature. For consistency and ease of comparisons, we decided our traits should all be scaled such that higher values were expected a priori to confer resistance to herbivores. Accordingly, we have rescaled specific leaf area (SLA) into its inverse, Leaf Mass per Area (LMA), and % Nitrogen to % non-nitrogen. In revising this manuscript, we have also revised our calculations of PRR to range only across observed trait values, and have updated our calculations of %PRV (previously called %PRR) for survival analyses to calculate model fit based on predictions on the logit scale, rather on the probability scale. Finally, we have systematized identification of Random Forest relationships as "synergy", "antagonism", or "unclear" using permutation testing.

## References

Agrawal, A. A. 2004a. “Plant Defense and Density Dependence in the Population Growth of Herbivores”. The American Naturalist 164(1): 113–120. https://doi.org/10.1086/420980

Agrawal, A. A. 2004b. “Resistance and Susceptibility of Milkweed: Competition, Root Herbivory, and Plant Genetic Variation”. Ecology 85(8): 2118–2133. https://doi.org/10.1890/03-4084

Agrawal, A. A. 2005. “Natural Selection on Common Milkweed (Asclepias Syriaca) by a Community of Specialized Insect Herbivores.” Evolutionary Ecology Research 7 (5): 651–67. http://www.evolutionary-ecology.com/abstracts/v07/1801.html.

Agrawal, A. A. 2011. “Current Trends in the Evolutionary Ecology of Plant Defence: Plant Defence Theory.” Functional Ecology 25 (2): 420–32. https://doi.org/10.1111/j.1365-2435.2010.01796.x.

Agrawal, A. A. 2017. Monarchs and Milkweed: A Migrating Butterfly, a Poisonous Plant, and Their Remarkable Story of Coevolution. Princeton University Press.

Agrawal, A. A. 2020. “A scale-dependent framework for trade-offs, syndromes, and specialization in organismal biology”. Ecology 101, e02924. https://doi.org/10.1002/ecy.2924

Agrawal, A. A., and M. Fishbein. 2006. “Plant Defense Syndromes.” Ecology 87 (7 Suppl): S132–49. https://doi.org/10.1890/0012-9658(2006)87[132:pds]2.0.co;2.

Agrawal, A. A., M. Fishbein, R. Jetter, J.-P. Salminen, J. B. Goldstein, A. E. Freitag, and J. P. Sparks. 2009. “Phylogenetic Ecology of Leaf Surface Traits in the Milkweeds (Asclepias Spp.): Chemistry, Ecophysiology, and Insect Behavior.” The New Phytologist 183 (3): 848–67. https://doi.org/10.1111/j.1469-8137.2009.02897.x.

Agrawal, A. A., K. Böröczky, M. Haribal, A. P. Hastings, R. A. White, R.W. Jiang, and C. Duplais. 2021. “Cardenolides, Toxicity, and the Costs of Sequestration in the Coevolutionary Interaction Between Monarchs and Milkweeds.” Proceedings of the National Academy of Sciences of the United States of America 118 (16). https://doi.org/10.1073/pnas.2024463118

Agrawal, A. A., E. E. Kearney, A. P. Hastings, and T. E. Ramsey. 2012. “Attenuation of the jasmonate burst, plant defensive traits, and resistance to specialist monarch caterpillars on shaded common milkweed (Asclepias syriaca)”. Journal of Chemical Ecology, 38 (7): 893–901. https://doi.org/10.1007/s10886-012-0145-3

Agrawal, A. A., and K. Konno. 2009. “Latex: A Model for Understanding Mechanisms, Ecology, and Evolution of Plant Defense Against Herbivory.” Annual Review of Ecology, Evolution, and Systematics 40 (1): 311–31. https://doi.org/10.1146/annurev.ecolsys.110308.120307.

Agrawal, A. A., E. T. Patrick, and A. P. Hastings. 2014. “Tests of the Coupled Expression of Latex and Cardenolide Plant Defense in Common Milkweed (Asclepias Syriaca).” Ecosphere 5 (10): art126. https://doi.org/10.1890/ES14-00161.1.

Aljbory, Z., and M.S. Chen. 2018. “Indirect Plant Defense Against Insect Herbivores: a Review.” Insect Science 25:2–23. https://doi.org/10.1111/1744-7917.12436

Barton, K. E. 2014. “Prickles, Latex, and Tolerance in the Endemic Hawaiian Prickly Poppy (Argemone Glauca): Variation between Populations, across Ontogeny, and in Response to Abiotic Factors.” Oecologia 174 (4): 1273–81. https://doi.org/10.1007/s00442-013-2836-z.

Benrey, B., and R. F. Denno. 1997. “The Slow-Growth--High-Mortality Hypothesis: a Test Using the Cabbage Butterfly.” Ecology 78:987–999. https://doi.org/10.1890/0012-9658(1997)078[0987:TSGHMH]2.0.CO;2

Berenbaum, M. R., J. K. Nitao, and A. R. Zangerl. 1991. “Adaptive Significance of Furanocoumarin Diversity in Pastinaca Sativa (Apiaceae).” Journal of Chemical Ecology 17 (1): 207–15. https://doi.org/10.1007/BF00994434.

Calcagno, M. P., J. Coll, J. Lloria, F. Faini, and M. E. Alonso-Amelot. 2002. “Evaluation of Synergism in the Feeding Deterrence of Some Furanocoumarins on Spodoptera littoralis.” Journal of Chemical Ecology 28:175–191. https://doi.org/10.1023/A:1013575121691

Carmona, D., M. J. Lajeunesse, and M. T. J. Johnson. 2013. “Plant Traits That Predict Resistance to Herbivores.” Functional Ecology 358–367. https://doi.org/10.1111/j.1365-2435.2010.01794.x

Cipollini, M. L., and E. W. Stiles. 1992. “Antifungal Activity of Ripe Ericaceous Fruits: Phenoliz-Acid Interactions and Palatability for Dispersers.” Biochemical Systematics and Ecology 20 (6): 501–14. https://doi.org/10.1016/0305-1978(92)90004-W.

Coley, P. D., M.-J. Endara, and T. A. Kursar. 2018. “Consequences of Interspecific Variation in Defenses and Herbivore Host Choice for the Ecology and Evolution of Inga, a Speciose Rainforest Tree.” Oecologia 187 (2): 361–76. https://doi.org/10.1007/s00442-018-4080-z.

Diawara, M. M., J. T. Trumble, K. K. White, W. G. Carson, and L. A. Martinez. 1993. “Toxicity of Linear Furanocoumarins to Spodoptera Exigua: Evidence for Antagonistic Interactions.” Journal of Chemical Ecology 19 (11): 2473–84. https://doi.org/10.1007/BF00980684.

Dobler, S., S. Dalla, V. Wagschal, and A. A. Agrawal. 2012. “Community-Wide Convergent Evolution in Insect Adaptation to Toxic Cardenolides by Substitutions in the Na,K-ATPase.” PNAS 109 (32): 13040–45. https://doi.org/10.1073/pnas.1202111109.

Duffey, S. S., and M. J. Stout. 1996. “Antinutritive and Toxic Components of Plant Defense against Insects.” Archives of Insect Biochemistry and Physiology 32 (1): 3–37. https://doi.org/10.1002/(sici)1520-6327(1996)32:1<3::aid-arch2>3.0.co;2-1.

Dussourd, D. E., and R. F. Denno. 1994. “Host Range of Generalist Caterpillars: Trenching Permits Feeding on Plants with Secretory Canals.” Ecology 75:69–78. https://doi.org/10.2307/1939383

Dyer, L. A., C. D. Dodson, J. O. Stireman 3rd, M. A. Tobler, A. M. Smilanich, R. M. Fincher, and D. K. Letourneau. 2003. “Synergistic Effects of Three Piper Amides on Generalist and Specialist Herbivores.” Journal of Chemical Ecology 29 (11): 2499–2514. https://doi.org/10.1023/a:1026310001958.

Ehleringer, J. 1982. “The Influence of Water Stress and Temperature on Leaf Pubescence Development in Encelia farinose.” American Journal of Botany 69:670–675. https://doi.org/10.1002/j.1537-2197.1982.tb13306.x

Eneji, A. E., S. Inanaga, S. Muranaka, J. Li, T. Hattori, P. An, and W. Tsuji. 2008. “Growth and Nutrient Use in Four Grasses Under Drought Stress as Mediated by Silicon Fertilizers.” Journal of Plant Nutrition 31:355–365. https://doi.org/10.1080/01904160801894913

Fajardo, A., and A. Siefert. 2018. “Intraspecific trait variation and the leaf economics spectrum across resource gradients and levels of organization.” Ecology 99:1024–1030. https://doi.org/10.1002/ecy.2194

Feeny, P. 1976. “Plant Apparency and Chemical Defense.” Pages 1–40 in J. W. Wallace and R. L. Mansell, eds. Biochemical Interaction Between Plants and Insects, Recent Advances in Phytochemistry. Springer US, Boston, MA. https://doi.org/10.1007/978-1-4684-2646-5_1

Fox, J. and S. Weisberg. 2019. “An R Companion to Applied Regression, Third edition.” Sage, Thousand Oaks CA.

Green, E. S., A. R. Zangerl, and M. R. Berenbaum. 2001. “Effects of Phytic Acid and Xanthotoxin on Growth and Detoxification in Caterpillars.” Journal of Chemical Ecology 27 (9): 1763–73. https://doi.org/10.1023/a:1010452507718.

Hay, M. E., Q. E. Kappel, and W. Fenical. 1994. “Synergisms in Plant Defenses against Herbivores: Interactions of Chemistry, Calcification, and Plant Quality.” Ecology 75 (6): 1714–26. https://doi.org/10.2307/1939631.

Hay, Mark E. 1996. “Defensive Synergisms? Reply to Pennings.” Ecology 77 (6): 1950–52. https://doi.org/10.2307/2265799.

Herms, D. A., and W. J. Mattson. 1992. “The Dilemma of Plants: To Grow or Defend.” The Quarterly Review of Biology 67 (3): 283–335. https://doi.org/10.1086/417659

Hill, M. S., N. A. Lopez, and K. A. Young. 2005. “Anti-Predator Defenses in Western North Atlantic Sponges with Evidence of Enhanced Defense through Interactions between Spicules and Chemicals.” Marine Ecology Progress Series 291: 93–102. https://doi.org/10.3354/meps291093.

Hooker, G. 2007. “Generalized Functional Anova Diagnostics for High-Dimensional Functions of Dependent Variables.” Journal of Computational and Graphical Statistics 16 (3): 709–32. https://doi.org/10.1198/106186007X237892

Hahn, P. G., A. A. Agrawal, K. I. Sussman, and J. L. Maron. 2019. “Population Variation, Environmental Gradients, and the Evolutionary Ecology of Plant Defense against Herbivory.” The American Naturalist 193 (1): 20–34. https://doi.org/10.1086/700838

Hooker, G., and L. Mentch. 2019. “Please Stop Permuting Features: An Explanation and Alternatives.” arXiv[stat.ME]. arXiv. http://arxiv.org/abs/1905.03151.

Huber, M., J. Epping, C. S. Gronover, J. Fricke, Z. Aziz, T. Brillatz, M. Swyers, et al. 2016. “A Latex Metabolite Benefits Plant Fitness under Root Herbivore Attack.” PLoS Biology 14 (1): e1002332. https://doi.org/10.1371/journal.pbio.1002332.

James, G., D. Witten, T. Hastie, and R. Tibshirani. 2021. An Introduction to Statistical Learning: with Applications in R. Springer, New York, NY.

Jones, A. C., J. E. Blum, and J. R. Pawlik. 2005. “Testing for Defensive Synergy in Caribbean Sponges: Bad Taste or Glass Spicules?” Journal of Experimental Marine Biology and Ecology 322 (1): 67–81. https://doi.org/10.1016/j.jembe.2005.02.009.

Jones, C. G., R. D. Firn. 1991. “On the Evolution of Plant Secondary Chemical Diversity.” Philosophical Transactions of the Royal Society of London. Series B, Biological Sciences 333 (1267): 273–80. https://doi.org/10.1098/rstb.1991.0077.

Jones, P. L., and A. A. Agrawal. 2019. “Beyond Preference and Performance: Host Plant Selection by Monarch Butterflies, Danaus Plexippus.” Oikos 128 (8): 1092-1102 https://doi.org/10.1111/oik.06001.

Jones, P. L., G. Petschenka, L. Flacht, and A. A. Agrawal. 2019. “Cardenolide Intake, Sequestration, and Excretion by the Monarch Butterfly along Gradients of Plant Toxicity and Larval Ontogeny.” Journal of Chemical Ecology 45 (3): 264–77. https://doi.org/10.1007/s10886-019-01055-7.

Kant, M. R., W. Jonckheere, B. Knegt, F. Lemos, J. Liu, B. C. J. Schimmel, C. A. Villarroel, et al. 2015. “Mechanisms and Ecological Consequences of Plant Defence Induction and Suppression in Herbivore Communities.” Annals of Botany 115 (7): 1015–51. https://doi.org/10.1093/aob/mcv054.

Kassambara, A. 2020. “ggpubr: ‘ggplot2’ Based Publication Ready Plots”. R package version 0.4.0.

Kursar, T. A., K. G. Dexter, J. Lokvam, R. T. Pennington, J. E. Richardson, M. G. Weber, E. T. Murakami, C. Drake, R. McGregor, and P. D. Coley. 2009. “The Evolution of Antiherbivore Defenses and Their Contribution to Species Coexistence in the Tropical Tree Genus Inga.” PNAS 106 (43): 18073–78. https://doi.org/10.1073/pnas.0904786106

Liaw, A. and M. Wiener. 2002. Classification and Regression by randomForest. R News 2(3), 18–-22.

Liu, X., K. Vrieling, and P. G. L. Klinkhamer. 2017. “Interactions between Plant Metabolites Affect Herbivores: A Study with Pyrrolizidine Alkaloids and Chlorogenic Acid.” Frontiers in Plant Science 8: 903. https://doi.org/10.3389/fpls.2017.00903.

Moles, A. T., B. Peco, I. R. Wallis, W. J. Foley, A. G. B. Poore, E. W. Seabloom, P. A. Vesk, et al. 2013. “Correlations between Physical and Chemical Defences in Plants: Tradeoffs, Syndromes, or Just Many Different Ways to Skin a Herbivorous Cat?” The New Phytologist 198 (1): 252–63. https://doi.org/10.1111/nph.12116.

Muff, S., E. B. Nilsen, R. B. O’Hara, and C. R. Nater. 2022. “Rewriting Results Sections in the Language of Evidence”. Trends in Ecology & Evolution. https://doi.org/10.1016/j.tree.2021.10.009

Muroi, H., and I. Kubo. 1993. “Combination Effects of Antibacterial Compounds in Green Tea Flavor against Streptococcus Mutans.” Journal of Agricultural and Food Chemistry 41 (7): 1102–5. https://doi.org/10.1021/jf00031a017.

Nelson, A. C., and T. A. Kursar. 1999. “Interactions among Plant Defense Compounds: A Method for Analysis.” Chemoecology 9 (2): 81–92. https://doi.org/10.1007/s000490050037.

Pennings, S. C. 1996. “Testing for Synergisms between Chemical and Mineral Defenses--A Comment.” Ecology 77 (6): 1948–50. https://doi.org/10.2307/2265798.

R Core Team. 2021. R: A language and environment for statistical computing. R Foundation for Statistical Computing, Vienna, Austria. https://www.R-project.org/.

Rasmann, S., and A. A. Agrawal. 2009. “Plant Defense against Herbivory: Progress in Identifying Synergism, Redundancy, and Antagonism between Resistance Traits.” Current Opinion in Plant Biology 12 (4): 473–78. https://doi.org/10.1016/j.pbi.2009.05.005.

Richards, L. A., A. E. Glassmire, K. M. Ochsenrider, A. M. Smilanich, C. D. Dodson, C. S. Jeffrey, and L. A. Dyer. 2016. “Phytochemical Diversity and Synergistic Effects on Herbivores.” Phytochemistry Reviews 15 (6): 1153–66. https://doi.org/10.1007/s11101-016-9479-8.

Richards, L. A., L. A. Dyer, A. M. Smilanich, and C. D. Dodson. 2010. “Synergistic Effects of Amides from Two Piper Species on Generalist and Specialist Herbivores.” Journal of Chemical Ecology 36 (10): 1105–13. https://doi.org/10.1007/s10886-010-9852-9.

Salazar, D., J. Lokvam, I. Mesones, M. V. Pilco, J. M. A. Zuñiga, P. de Valpine, and P. V. Fine. 2018. “Origin and maintenance of chemical diversity in a species-rich tropical tree lineage.” Nature Ecology & Evolution, 2(6), 983–990. https://doi.org/10.1038/s41559-018-0552-0

Sanczuk, P., S. Govaert, C. Meeussen, K. De Pauw, T. Vanneste, L. Depauw, X. Moreira, et al. 2020. “Small Scale Environmental Variation Modulates Plant Defence Syndromes of Understory Plants in Deciduous Forests of Europe.” Global Ecology and Biogeography: A Journal of Macroecology 30(1): 205–2019. https://doi.org/10.1111/geb.13216.

Schindelin, J., I. Arganda-Carreras, E. Frise, V. Kaynig, M. Longair, T. Pietzsch, S. Preibisch, et al. 2012. “Fiji: An Open-Source Platform for Biological-Image Analysis.” Nature Methods 9 (7): 676. https://doi.org/10.1038/nmeth.2019

Schuman, M. C., K. Barthel, and I. T. Baldwin. 2012. “Herbivory-Induced Volatiles Function as Defenses Increasing Fitness of the Native Plant Nicotiana Attenuata in Nature.” eLife 1: e00007. https://doi.org/10.7554/eLife.00007.

Scott, I. M., E. Puniani, T. Durst, D. Phelps, S. Merali, R. A. Assabgui, P. Sánchez-Vindas, L. Poveda, B. J. R. Philogene, and J. T. Arnason. 2002. “Insecticidal Activity of Piper Tuberculatum Jacq. Extracts: Synergistic Interaction of Piperamides.” Agricultural and Forest Entomology 4 (2): 137–44.

Shipley, B., M. J. Lechowicz, I. Wright, and P. B. Reich. 2006. “Fundamental Trade-Offs Generating the Worldwide Leaf Economic Spectrum”. Ecology 87: 535-541 https://doi.org/10.1890/05-1051

da Silva, D. M., and M. A. Batalha. 2011. “Defense Syndromes against Herbivory in a Cerrado Plant Community.” Plant Ecology 212 (2): 181–93. https://doi.org/10.1007/s11258-010-9813-y.

Simpson, S. J., and D. Raubenheimer. 2001. “The Geometric Analysis of Nutrient–allelochemical Interactions: A Case Study Using Locusts.” Ecology 82 (2): 422–39. https://doi.org/10.1890/0012-9658(2001)082[0422:tgaona]2.0.co;2.

Stamp, N. E., and Y. Yang. 1996. “Response of Insect Herbivores to Multiple Allelochemicals under Different Thermal Regimes.” Ecology 77 (4): 1088–1102. https://doi.org/10.2307/2265578.

Steppuhn, A., and I. T. Baldwin. 2007. “Resistance Management in a Native Plant: Nicotine Prevents Herbivores from Compensating for Plant Protease Inhibitors.” Ecology Letters 10 (6): 499–511. https://doi.org/10.1111/j.1461-0248.2007.01045.x.

Stermitz, F. R., P. Lorenz, J. N. Tawara, L. A. Zenewicz, and K. Lewis. 2000. “Synergy in a Medicinal Plant: Antimicrobial Action of Berberine Potentiated by 5’-Methoxyhydnocarpin, a Multidrug Pump Inhibitor.” Proceedings of the National Academy of Sciences 97 (4): 1433–37. https://www.pnas.org/content/97/4/1433.short.

Tallarida, R. J. 2000. Drug Synergism and Dose-Effect Data Analysis. CRC Press. https://doi.org/10.1201/9781420036107

Tao, L., K. M. Hoang, M. D. Hunter, and J. C. de Roode. 2016. “Fitness Costs of Animal Medication: Antiparasitic Plant Chemicals Reduce Fitness of Monarch Butterfly Hosts.” The Journal of Animal Ecology 85 (5): 1246–54. https://doi.org/10.1111/1365-2656.12558.

Van Zandt, P. A., and A. A. Agrawal. 2004. “Specificity of Induced Plant Responses to Specialist Herbivores of the Common Milkweed Asclepias syriaca.” Oikos 104: 401–409. https://doi.org/10.1111/j.0030-1299.2004.12964.x

Wetzel, W. C., and S. R. Whitehead. 2020. “The Many Dimensions of Phytochemical Diversity: Linking Theory to Practice.” Ecology letters 23:16–32. https://doi.org/10.1111/ele.13422

Whitehead, S. R., and M. D. Bowers. 2014. “Chemical Ecology of Fruit Defence: Synergistic and Antagonistic Interactions among Amides from Piper.” Functional Ecology 28 (5): 1094–1106. https://doi.org/10.1111/1365-2435.12250.

Whitehead, S. R., Bass, E., Corrigan, A., Kessler, A. & Poveda, K. 2021. “Interaction Diversity Explains the Maintenance of Phytochemical Diversity.” Ecology Letters https://doi.org/10.1111/ele.13736

Wickham, H. 2016. “ggplot2: Elegant Graphics for Data Analysis”. Springer-Verlag New York.

Wickham, H., M. Averick, J. Bryan, W. Chang, L. D. McGowan, R. François, G. Grolemund, A. Hayes et al. 2019. “Welcome to the tidyverse.” Journal of Open Source Software, 4(43): 1686, https://doi.org/10.21105/joss.01686

Wilke, C. 2020. “cowplot: Streamlined Plot Theme and Plot Annotations for ‘ggplot2.’“ R package version 1.1.1.

Wood, S.N. 2011. “Fast Stable Restricted Maximum Likelihood and Marginal Likelihood Estimation of Semiparametric Generalized Linear Models”. Journal of the Royal Statistical Society (B) 73 (1): 3–36 https://doi.org/10.1111/j.1467-9868.2010.00749.x

Wright, I. J., and K. Cannon. 2001. “Relationships between leaf lifespan and structural defences in a low-nutrient, sclerophyll flora.” Functional Ecology 15:351–359. https://doi.org/10.1046/j.1365-2435.2001.00522.x

Wright, I. J., Reich P. B., Westoby, M., Ackerly, D. D., Baruch, Z., Bongers, F. Cavender-Bares, J., et al. 2004. “The Worldwide Leaf Economics Spectrum.” Nature 428:821–827. https://doi.org/10.1038/nature02403

Zahoor, R., H. Dong, M. Abid, W. Zhao, Y. Wang, and Z. Zhou. 2017. “Potassium Fertilizer Improves Drought Stress Alleviation Potential in Cotton by Enhancing Photosynthesis and Carbohydrate Metabolism.” Environmental and Experimental Botany 137:73–83. https://doi.org/10.1016/j.envexpbot.2017.02.002

Zalucki, M. P., and L. P. Brower. 1992. “Survival of First Instar Larvae of Danaus plexippus (Lepidoptera: Danainae) in Relation to Cardiac Glycoside and Latex Content of Asclepias humistrata (Asclepiadaceae)”. Chemoecology 3:81–93. https://doi.org/10.1007/BF01245886

Zalucki, M. P., Li. P. Brower, and A. Alonso-M. 2001. “Detrimental Effects of Latex and Cardiac Glycosides on Survival and Growth of First-Instar Monarch Butterfly Larvae Danaus Plexippus Feeding on the Sandhill Milkweed Asclepias Humistrata.” Ecological Entomology 26 (2): 212–24. https://doi.org/10.1046/j.1365-2311.2001.00313.x

Zalucki, Myron P., Anthony R. Clarke, and Stephen B. Malcolm. 2002. “Ecology and Behavior of First Instar Larval Lepidoptera.” Annual Review of Entomology 47 (1): 361–93. https://doi.org/10.1146/annurev.ento.47.091201.145220.

Züst, Tobias, Georg Petschenka, Amy P. Hastings, and Anurag A. Agrawal. 2019. “Toxicity of Milkweed Leaves and Latex: Chromatographic Quantification Versus Biological Activity of Cardenolides in 16 Asclepias Species.” Journal of Chemical Ecology 45 (1): 50–60. https://doi.ogikrg/10.1007/s10886-018-1040-3.

