## Appendix S1 for "Plant defense synergies and antagonisms affect performance of specialist herbivores of common milkweed"

**Process for PRR and %PRV calculations**

1. (For random forests) The Random Forest model was necessarily fit to all plant defense traits. In order to estimate how varying only the two traits of interest would affect herbivore performance, we had to identify the grid of values of those two traits we want to predict on, calculate predictions for each of those grid points *for each of the plants we observed* (e.g. with other traits set to the values of each of the observed plants), and then average the predictions across the observed plants. For each real plant measured, we generated 100 hypothetical plants, in which all trait values were the same as the real plant, except that values of the two traits of interest were varied across a 10 x 10 grid ranging from the smallest to largest values observed in any real plant. Because the outputs of Random Forests are piecewise constant at fine scales, it was important to avoid partitioning the trait space too finely (Giles Hooker, personal communication). For an empirical data set with N plants, this gave us 100N hypothetical plants.

(For regression models) Since our regression models do not include traits aside from the two traits of interest, we generated hypothetical plants defined only by the two traits of interest, spanning a grid 50x50 grid of grid ranging from the smallest to the largest values observed for each of the two traits. Regression models aren’t piecewise constant, so finer partitioning was not problematic, and we found that using a 50x50 grid of traits produced results similar to higher resolution grids, and could be computed in a reasonable amount of time (CBE, unpublished simulations).

2. Using the fitted model (Random Forest or regression), we predicted herbivore performance on the hypothetical plants. These predicted performances were kept on appropriate link scales (for log final weight, predictions were estimated log final weight; for survival, predictions were the logit of estimated survival probability).

3.	We fit a two-way ANOVA model to the predicted response of the form

response ~ trait1 + trait2

where the trait values of trait 1 and trait 2 were treated as factors. This new model captured all additive variation in the predictions of the original model; any residuals of the new model represented non-additive relationships in the regression or random forest model, including synergies or antagonisms. In the case of Random Forests, this captured non-additivity even if the independent response to each trait was nonlinear, or if synergies or antagonisms were not consistent across different levels of the traits. For this model we used a weighted ANOVA, where each observation from the grid of predicted values was weighted by how close it was (in terms of defense traits) to an actual empirical observation, using kernel density estimation to generate appropriate weights. This focused our results on meaningful patterns driven by the data, rather than spurious patterns driven by extrapolation of Random Forest or regression models outside the range of the data (Hooker and Mentch 2019).

4.	From the fitted ANOVA model, we divided the residual sum of squares by the total sum of squares, and multiplied by 100 to put our results on a percent scale. This was our %PRV, and represented the relative role of non-additive patterns in explaining our regression or Random Forest model predictions. In the case of Random Forests, for which there is not a P value equivalent, we used this percent as our metric for identifying important synergies and antagonisms.

5.	To calculate PRR,

(For Random Forests) We take a similar approach to step (1), except predicting observed values of the two traits of interest, rather than a 10 x 10 grid. For each of the n plants in the data, we took the values of our two traits of interest x_1_ and x_2_, and to estimate herbivore performance for those two trait values, we averaged the predicted herbivore performance on hypothetical plants identical to the observed plants, except that the two traits of interest were set to x_1_ and x_2_. This produced n estimated herbivore performance measures (each one an average of n predictions), one associated with each pair of observed trait values; PRR was the range of these estimates.

(For regression models): using the coefficients of our fitted regression model, we predicted herbivore performance on each plant (ignoring the fitted random effects of plant ID, as that term captured other variation). PRR is the range of these predicted performance values on the response scale (log final weight or probability of survival).

Because Random Forest regression struggles with missing data, we removed observations with one or more missing plant trait measurement. Random Forests also do not have appropriate ways to account for random effects, so we do not include the plant identity terms that were present in the regression analyses.

**Simulating interactions**

To simulate various functional forms of synergy and antagonism, we defined defense efficacy as a specified function of two hypothetical traits X and Y, where larger values indicate better defense (e.g. the first column in Fig 2a-f). We then used that function to simulate herbivore performance at a 10 X 10 grid of trait values, analogous to our fitted Random Forest predictions above (e.g. the second column in Fig 2a-f). For this set of simulated herbivore performance data, we fit an additive regression model of the form

efficacy ~ trait X + trait Y

with traits X and Y treated as factors (this is the same as the model fitted in *Random Forest* step 5 above). We assumed that traits X and Y were scaled to have values between 0 and 1, and defined herbivore performance as 10 minus the defense efficacy; as we were looking for qualitative patterns, these specific values do not qualitatively change our results (CBE, unpublished simulations). Code to simulate any arbitrary functional form of synergies or antagonism is available on Figshare.
