## Supplementary material for "Plant defense synergies and antagonisms affect performance of specialist herbivores of common milkweed": Figure S3

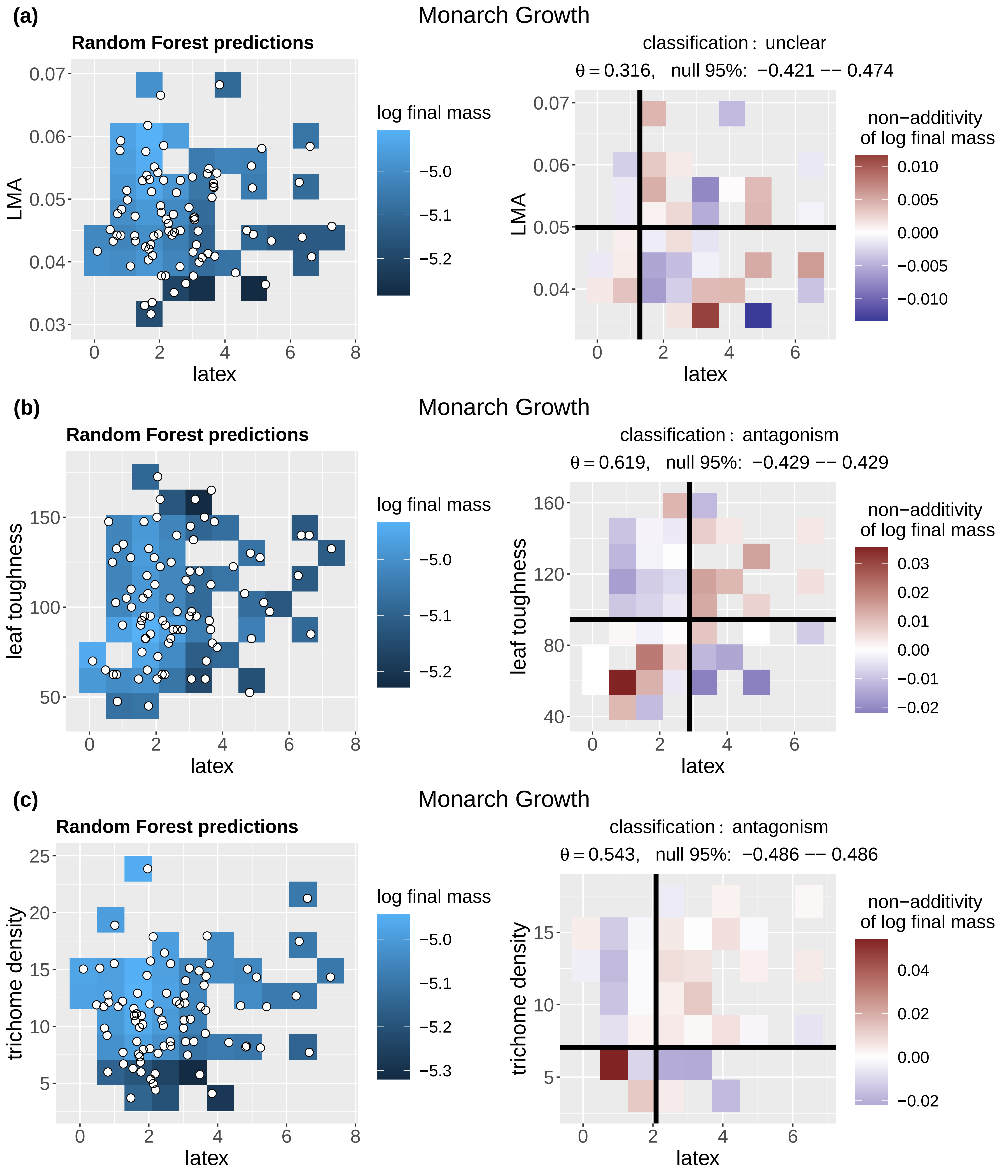

(a)
Monarch Growth
Random Forest predictions
LMA
latex
log final mass
classification : unclear
θ = 0.316, null 95%: −0.421 −− 0.474
non−additivity of log final mass
(b)
Monarch Growth
Random Forest predictions
leaf toughness
latex
log final mass
classification : antagonism
θ = 0.619, null 95%: −0.429 −− 0.429
non−additivity of log final mass
(c)
Monarch Growth
Random Forest predictions
trichome density
latex
log final mass
classification : antagonism
θ = 0.543, null 95%: −0.486 −− 0.486
non−additivity of log final mass

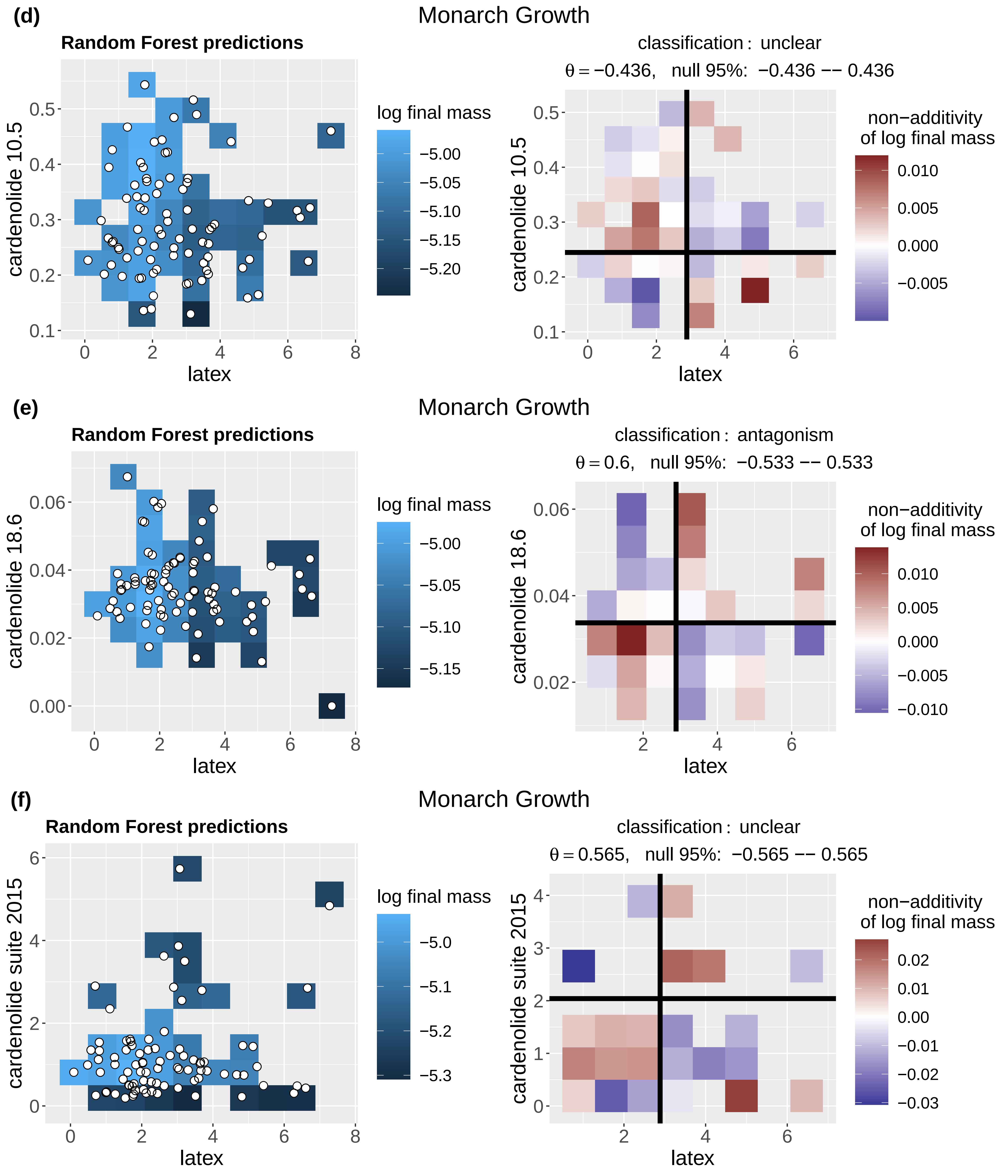

(d)
Monarch Growth
Random Forest predictions
log final mass
cardenolide 10.5
latex
classification : unclear
θ = −0.436, null 95%: −0.436 −− 0.436
non−additivity of log final mass
(e)
Monarch Growth
Random Forest predictions
log final mass
cardenolide 18.6
latex
classification : antagonism
θ = 0.6, null 95%: −0.533 −− 0.533
non−additivity of log final mass
(f)
Monarch Growth
Random Forest predictions
log final mass
cardenolide suite 2015
latex
classification : unclear
θ = 0.565, null 95%: −0.565 −− 0.565
non−additivity of log final mass

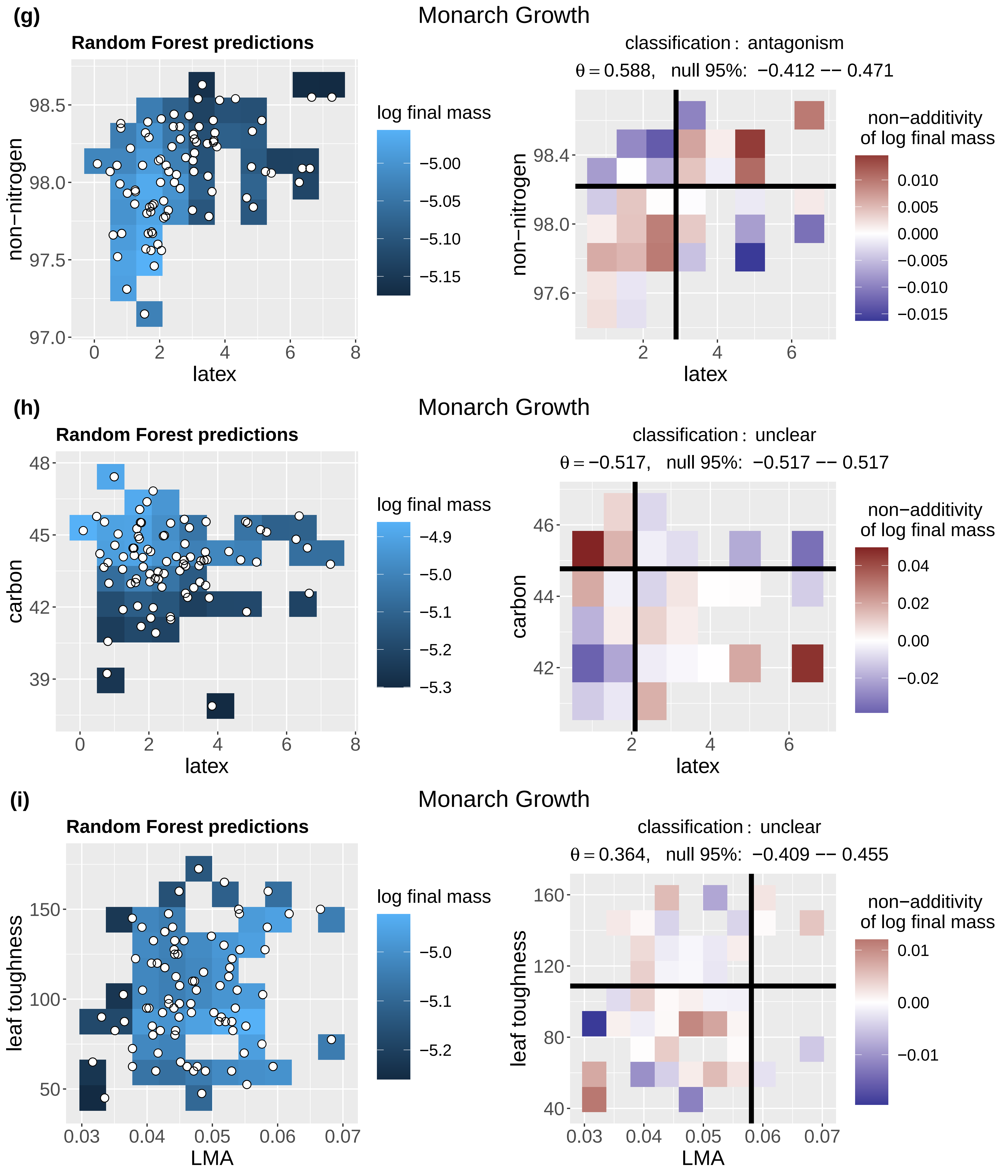
(g)
Monarch Growth
Random Forest predictions
log final mass
−5.00
−5.05
−5.10
−5.15
non−nitrogen
latex
classification : antagonism
θ = 0.588, null 95%: −0.412 −− 0.471
non−additivity of log final mass
0.010
0.005
0.000
−0.005
−0.010
−0.015
(h)
Monarch Growth
Random Forest predictions
log final mass
−4.9
−5.0
−5.1
−5.2
−5.3
carbon
latex
classification : unclear
θ = −0.517, null 95%: −0.517 −− 0.517
non−additivity of log final mass
0.04
0.02
0.00
−0.02
(i)
Monarch Growth
Random Forest predictions
log final mass
−5.0
−5.1
−5.2
leaf toughness
LMA
classification : unclear
θ = 0.364, null 95%: −0.409 −− 0.455
non−additivity of log final mass
0.01
0.00
−0.01

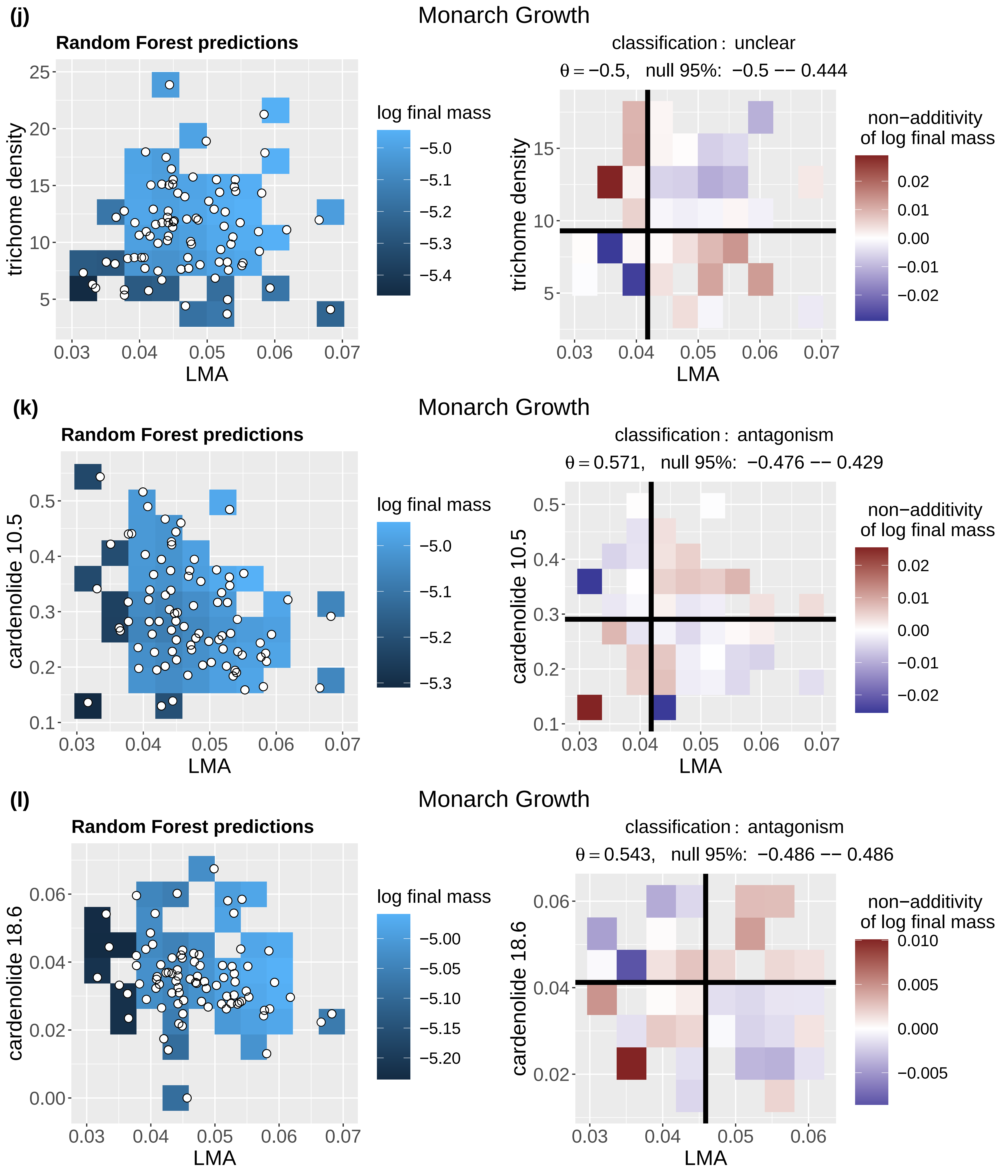

(j)
Monarch Growth
Random Forest predictions
trichome density
LMA
log final mass
−5.0
−5.1
−5.2
−5.3
−5.4
classification : unclear
θ = −0.5, null 95%: −0.5 −− 0.444
non−additivity of log final mass
0.02
0.01
0.00
−0.01
−0.02
(k)
Monarch Growth
Random Forest predictions
cardenolide 10.5
LMA
log final mass
−5.0
−5.1
−5.2
−5.3
classification : antagonism
θ = 0.571, null 95%: −0.476 −− 0.429
non−additivity of log final mass
0.02
0.01
0.00
−0.01
−0.02
(l)
Monarch Growth
Random Forest predictions
cardenolide 18.6
LMA
log final mass
−5.00
−5.05
−5.10
−5.15
−5.20
classification : antagonism
θ = 0.543, null 95%: −0.486 −− 0.486
non−additivity of log final mass
0.010
0.005
0.000
−0.005

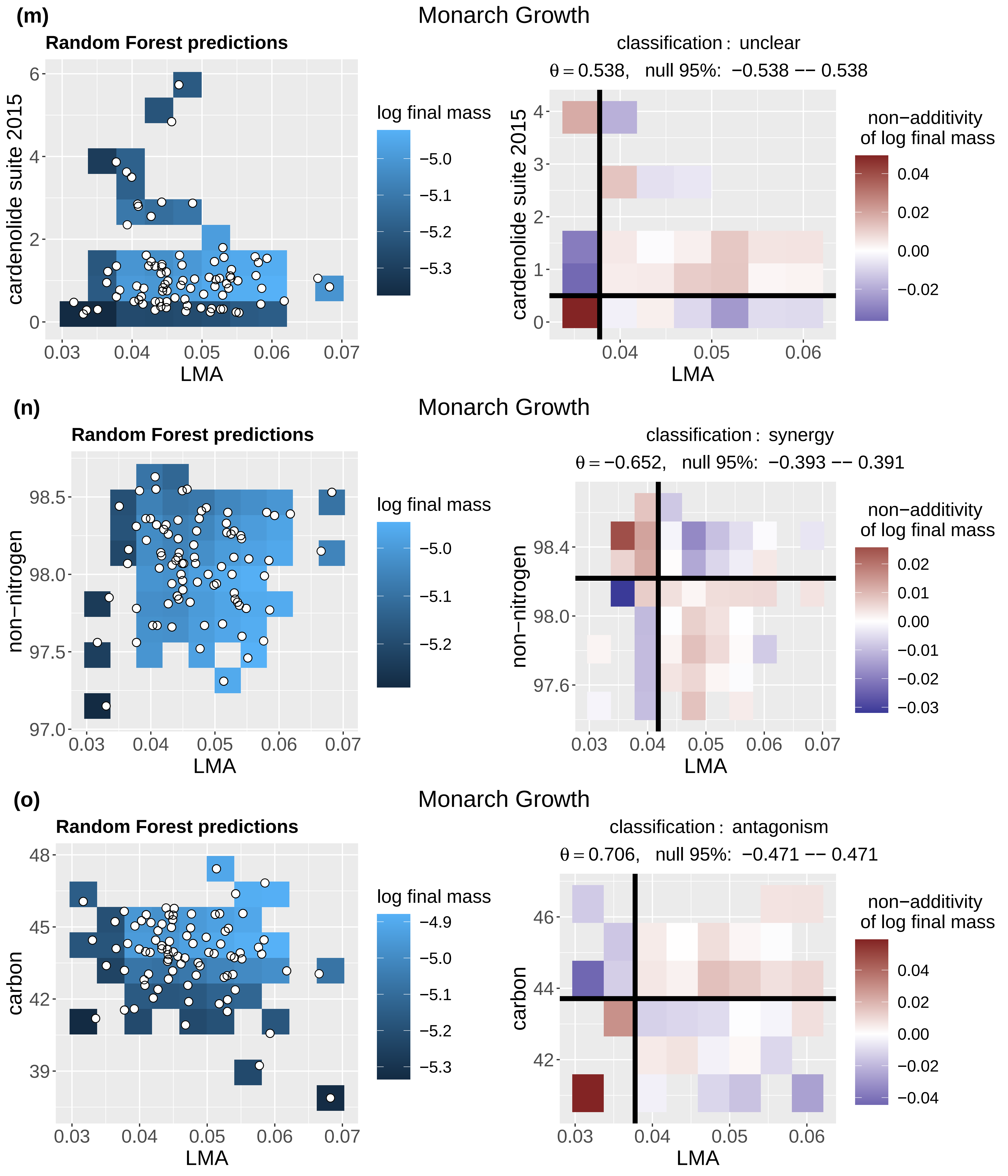

(m)
Monarch Growth
Random Forest predictions
cardenolide suite 2015
LMA
log final mass
classification : unclear
θ = 0.538, null 95%: −0.538 −− 0.538
non−additivity
of log final mass
(n)
Monarch Growth
Random Forest predictions
non−nitrogen
LMA
log final mass
classification : synergy
θ = −0.652, null 95%: −0.393 −− 0.391
non−additivity
of log final mass
(o)
Monarch Growth
Random Forest predictions
carbon
LMA
log final mass
classification : antagonism
θ = 0.706, null 95%: −0.471 −− 0.471
non−additivity
of log final mass

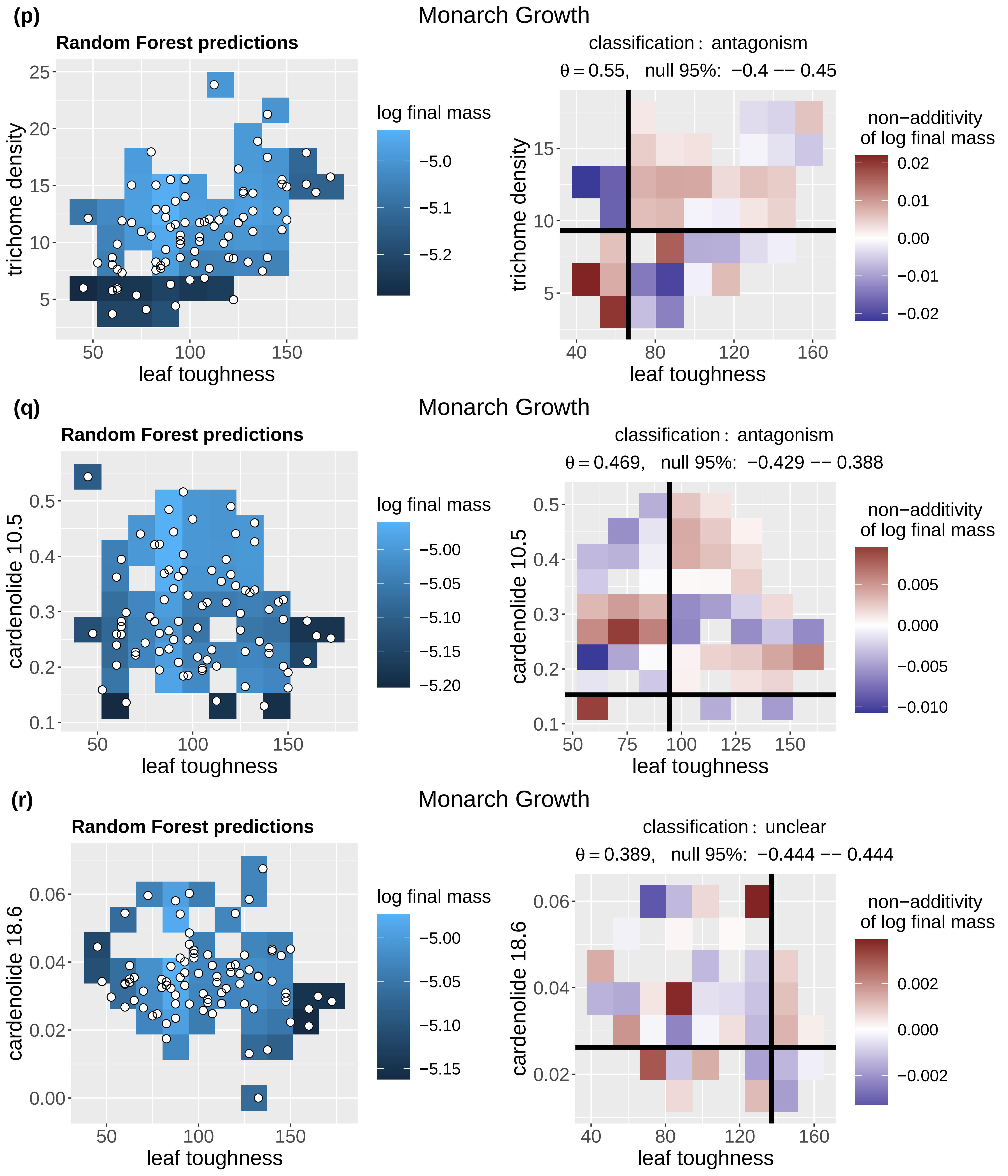
(p)
Monarch Growth
Random Forest predictions
trichome density
leaf toughness
log final mass
−5.0
−5.1
−5.2
classification : antagonism
θ = 0.55, null 95%: −0.4 −− 0.45
non−additivity
of log final mass
0.02
0.01
0.00
−0.01
−0.02
(q)
Monarch Growth
Random Forest predictions
cardenolide 10.5
leaf toughness
log final mass
−5.00
−5.05
−5.10
−5.15
−5.20
classification : antagonism
θ = 0.469, null 95%: −0.429 −− 0.388
non−additivity
of log final mass
0.005
0.000
−0.005
−0.010
(r)
Monarch Growth
Random Forest predictions
cardenolide 18.6
leaf toughness
log final mass
−5.00
−5.05
−5.10
−5.15
classification : unclear
θ = 0.389, null 95%: −0.444 −− 0.444
non−additivity
of log final mass
0.002
0.000
−0.002

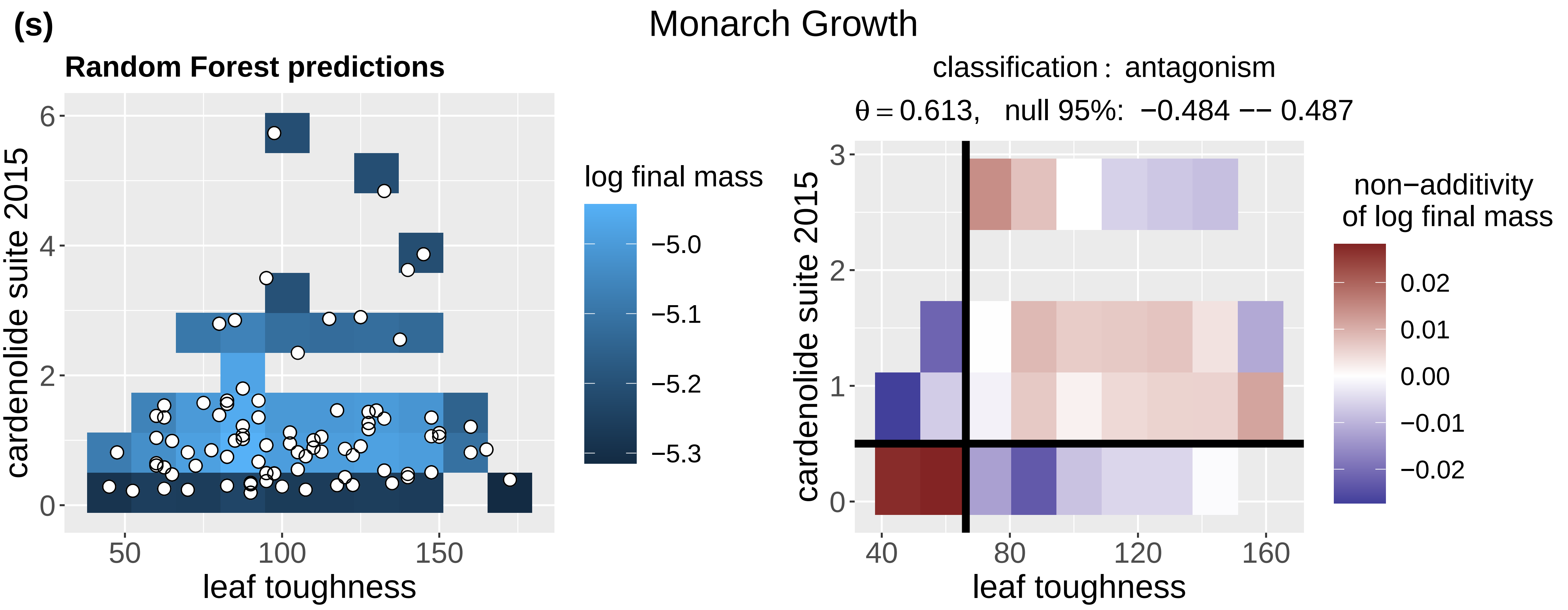
(s)
Monarch Growth
Random Forest predictions
log final mass
−5.0
−5.1
−5.2
−5.3
cardenolide suite 2015
leaf toughness
classification : antagonism
θ = 0.613, null 95%: −0.484 −− 0.487
non−additivity
of log final mass
0.02
0.01
0.00
−0.01
−0.02

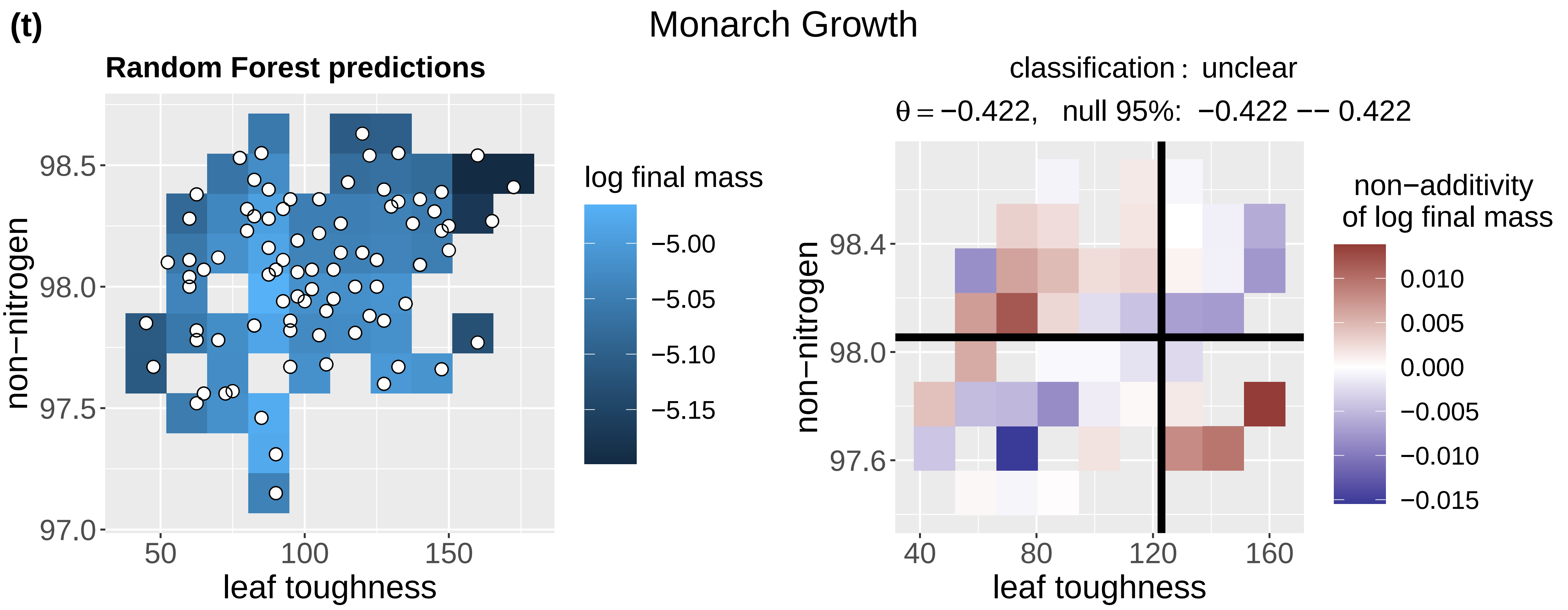
(t)
Monarch Growth
Random Forest predictions
log final mass
−5.00
−5.05
−5.10
−5.15
non−nitrogen
leaf toughness
classification : unclear
θ = −0.422, null 95%: −0.422 −− 0.422
non−additivity
of log final mass
0.010
0.005
0.000
−0.005
−0.010
−0.015

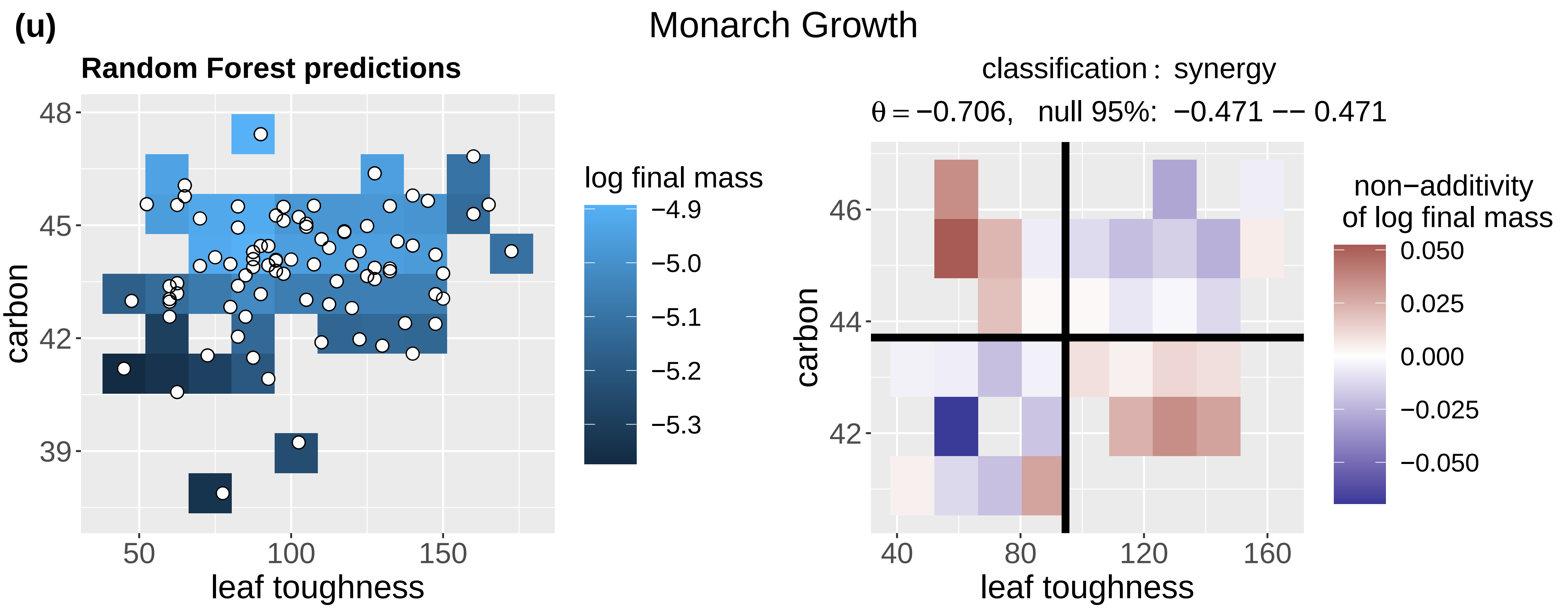
(u)
Monarch Growth
Random Forest predictions
log final mass
−4.9
−5.0
−5.1
−5.2
−5.3
carbon
leaf toughness
classification : synergy
θ = −0.706, null 95%: −0.471 −− 0.471
non−additivity
of log final mass
0.050
0.025
0.000
−0.025
−0.050

(v)

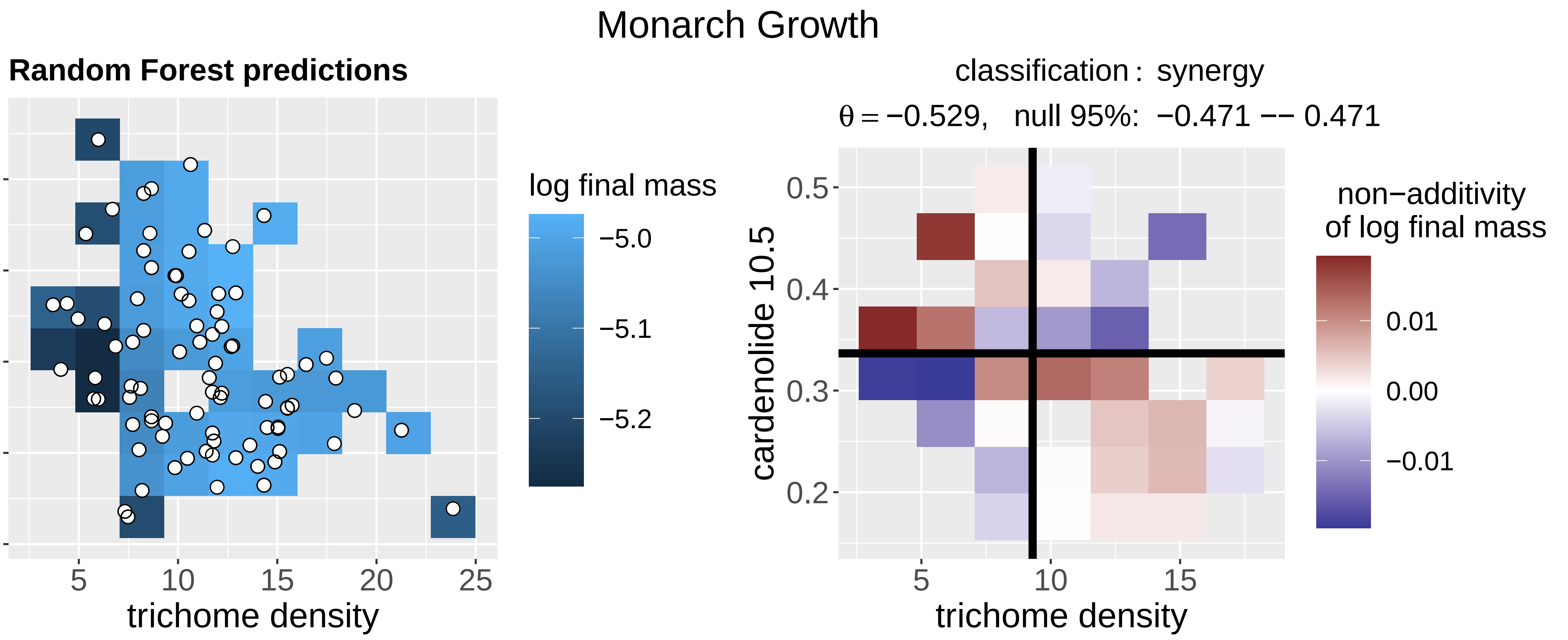
Monarch Growth
Random Forest predictions
cardenolide 10.5
trichome density
log final mass
−5.0
−5.1
−5.2
classification : synergy
θ = −0.529, null 95%: −0.471 −− 0.471
non−additivity
of log final mass
0.01
0.00
−0.01

(w)

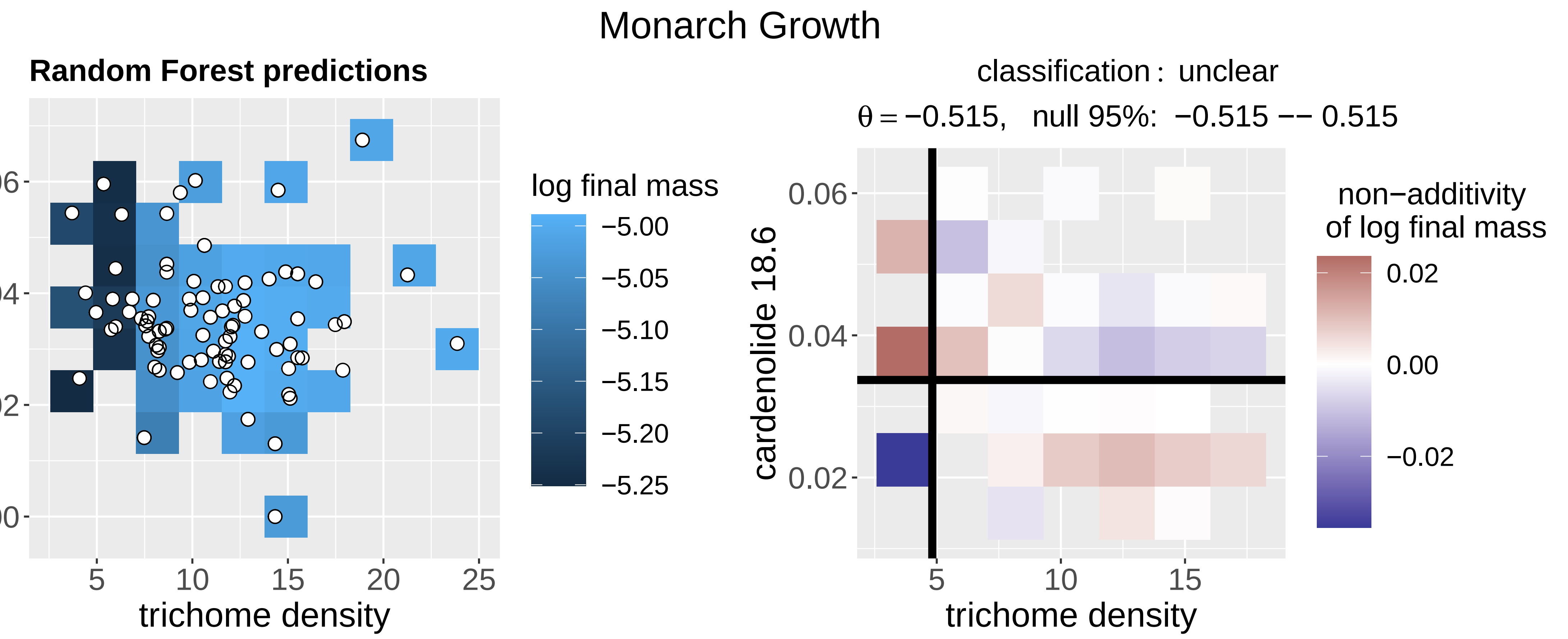
Monarch Growth
Random Forest predictions
cardenolide 18.6
trichome density
log final mass
−5.00
−5.05
−5.10
−5.15
−5.20
−5.25
classification : unclear
θ = −0.515, null 95%: −0.515 −− 0.515
non−additivity
of log final mass
0.02
0.00
−0.02

(x)

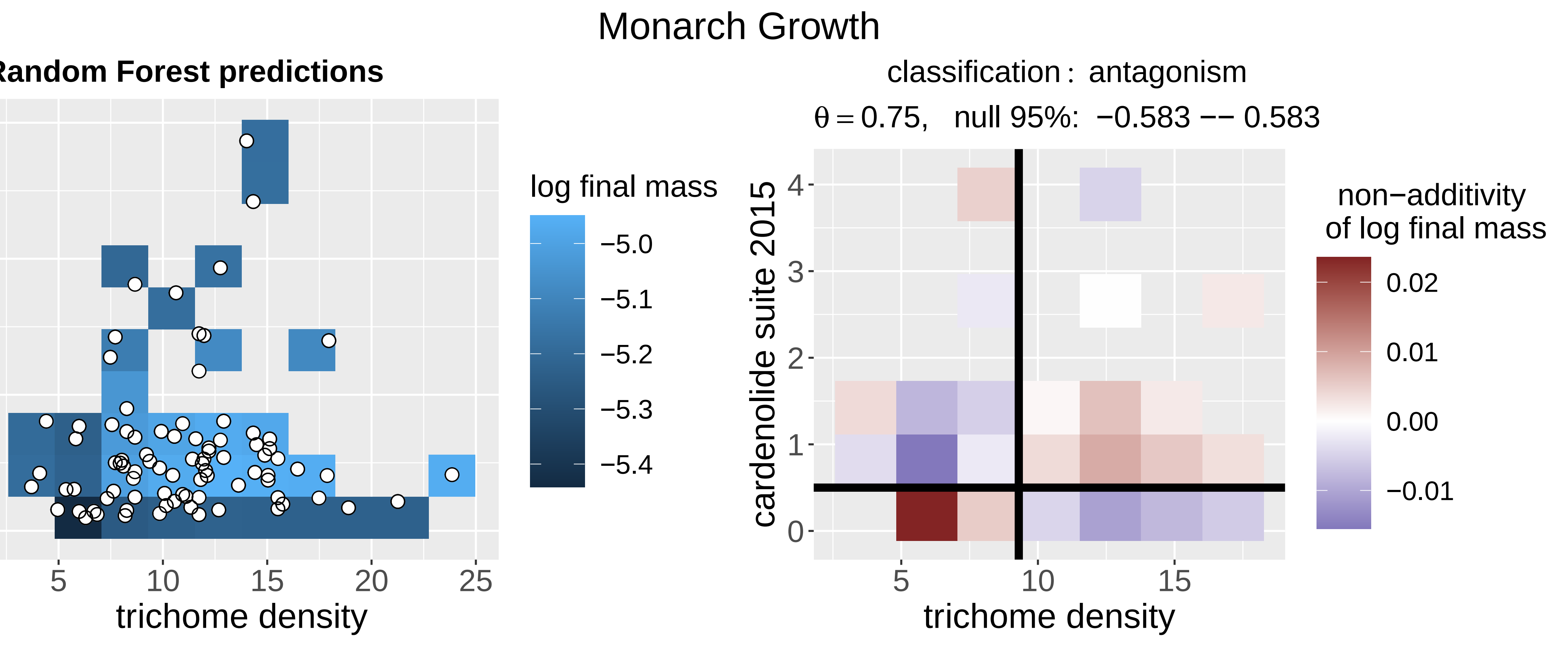
Monarch Growth
Random Forest predictions
cardenolide suite 2015
trichome density
log final mass
−5.0
−5.1
−5.2
−5.3
−5.4
classification : antagonism
θ = 0.75, null 95%: −0.583 −− 0.583
non−additivity
of log final mass
0.02
0.01
0.00
−0.01

(y)

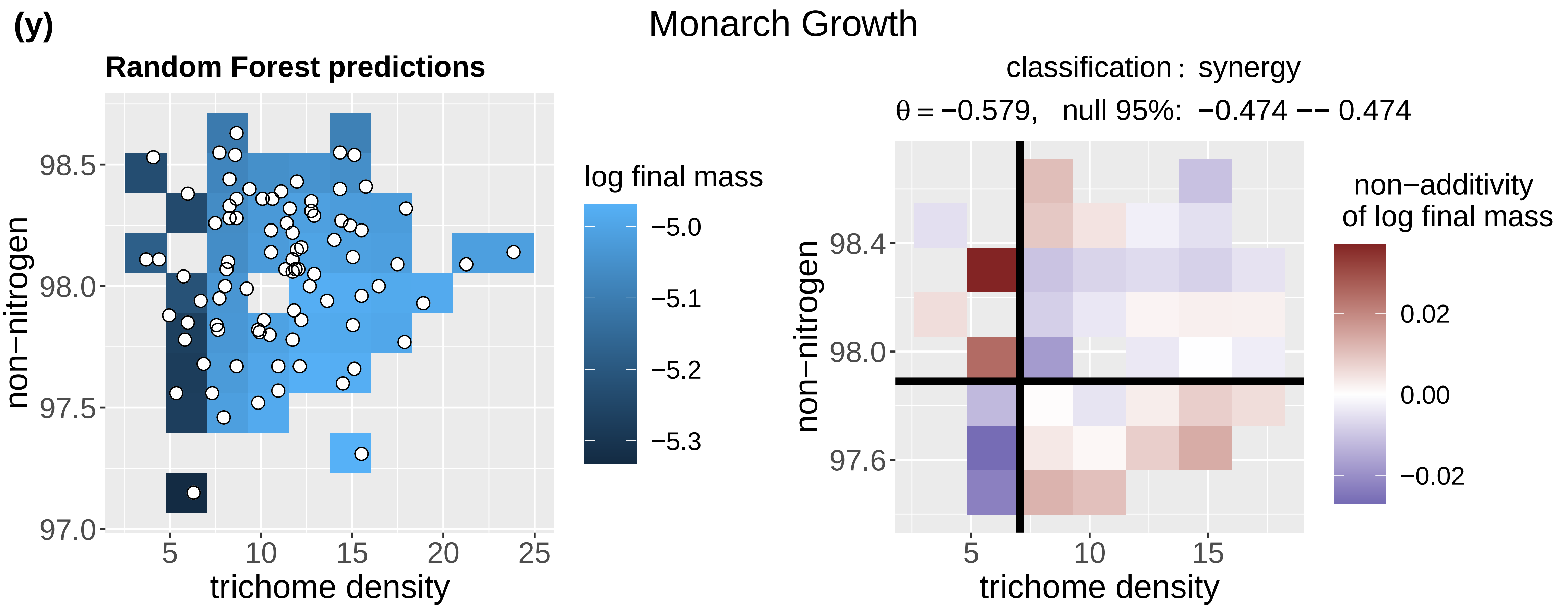
Monarch Growth
Random Forest predictions
log final mass
−5.0
−5.1
−5.2
−5.3
non−nitrogen
98.5
98.0
97.5
97.0
5
10
15
20
25
trichome density
classification : synergy
θ = −0.579, null 95%: −0.474 −− 0.474
non−additivity
of log final mass
0.02
0.00
−0.02
non−nitrogen
98.4
98.0
97.6
5
10
15
trichome density

(z)

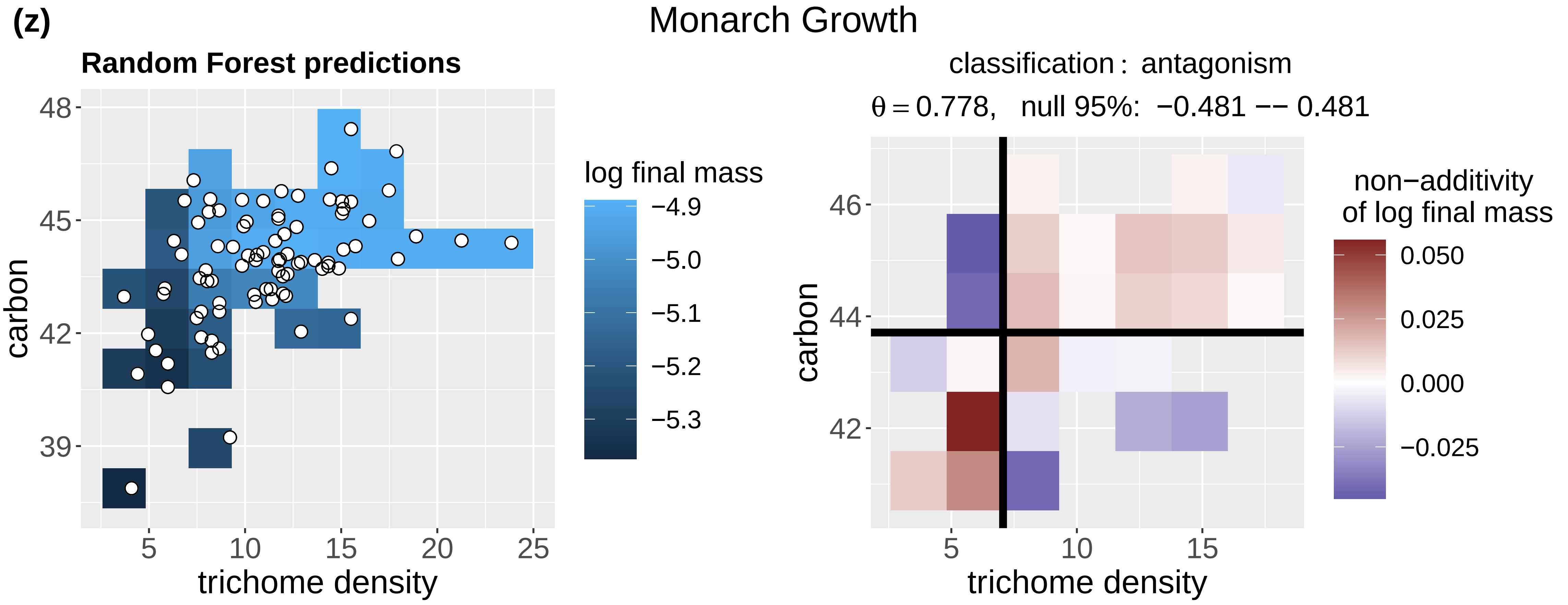
Monarch Growth
Random Forest predictions
log final mass
−4.9
−5.0
−5.1
−5.2
−5.3
carbon
48
45
42
39
5
10
15
20
25
trichome density
classification : antagonism
θ = 0.778, null 95%: −0.481 −− 0.481
non−additivity
of log final mass
0.050
0.025
0.000
−0.025
carbon
46
44
42
5
10
15
trichome density

(aa)

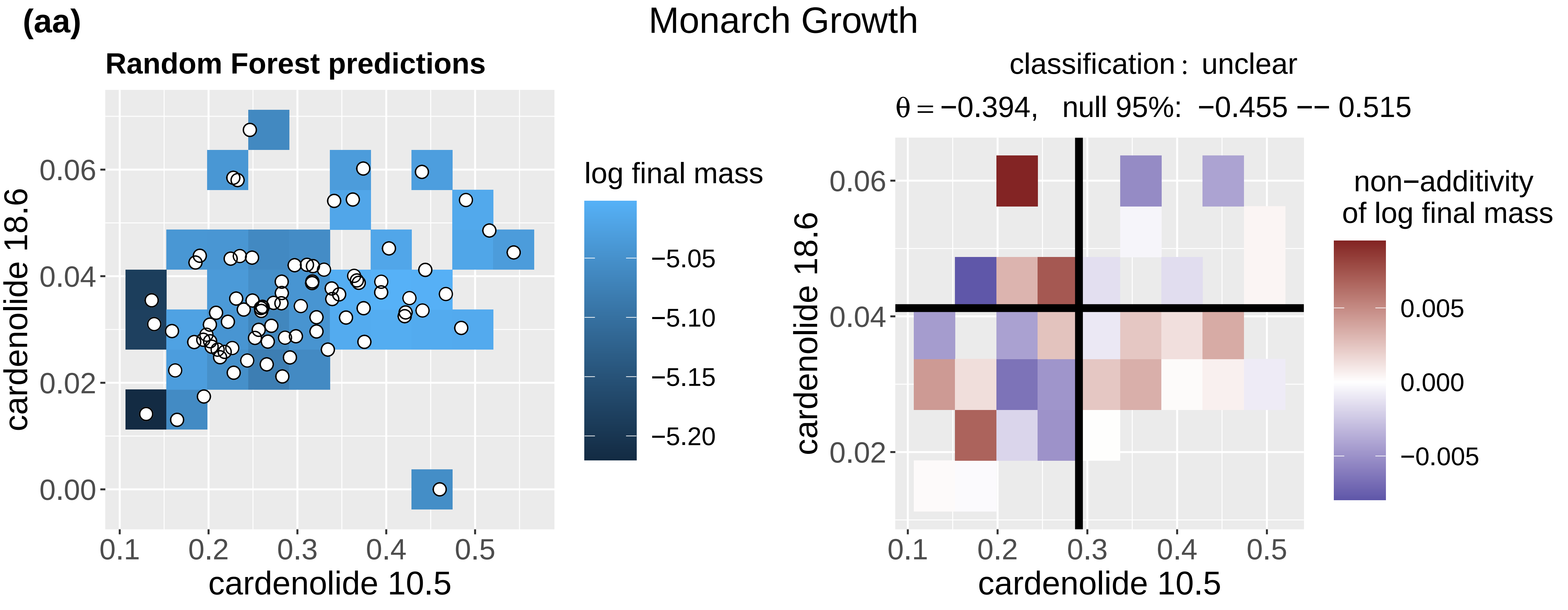
Monarch Growth
Random Forest predictions
log final mass
−5.05
−5.10
−5.15
−5.20
cardenolide 18.6
0.06
0.04
0.02
0.00
0.1
0.2
0.3
0.4
0.5
cardenolide 10.5
classification : unclear
θ = −0.394, null 95%: −0.455 −− 0.515
non−additivity
of log final mass
0.005
0.000
−0.005
cardenolide 18.6
0.06
0.04
0.02
0.1
0.2
0.3
0.4
0.5
cardenolide 10.5

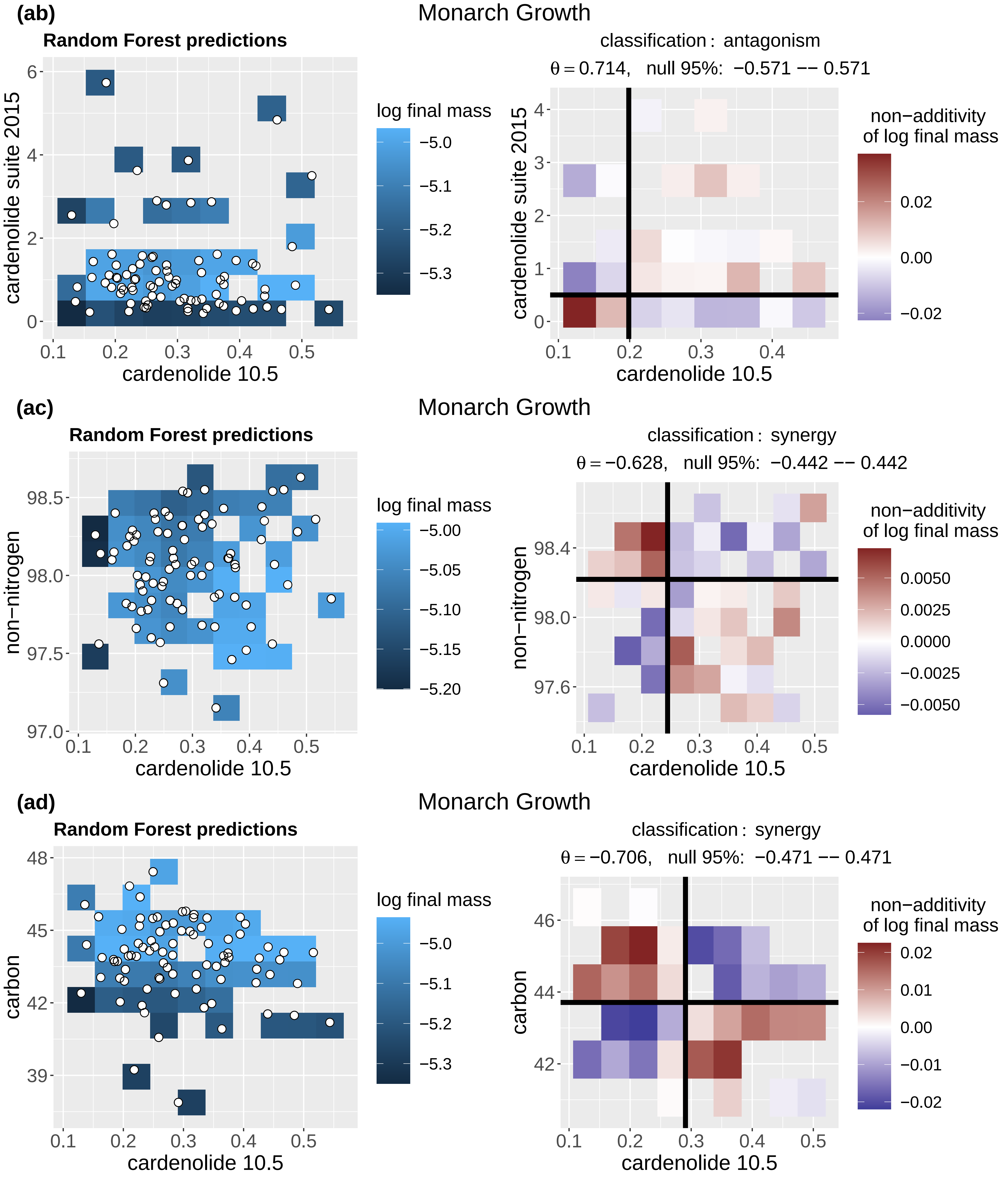
(ab)
Monarch Growth
Random Forest predictions
cardenolide suite 2015
cardenolide 10.5
log final mass
classification : antagonism
θ = 0.714, null 95%: −0.571 −− 0.571
non−additivity of log final mass
(ac)
Monarch Growth
Random Forest predictions
non−nitrogen
cardenolide 10.5
log final mass
classification : synergy
θ = −0.628, null 95%: −0.442 −− 0.442
non−additivity of log final mass
(ad)
Monarch Growth
Random Forest predictions
carbon
cardenolide 10.5
log final mass
classification : synergy
θ = −0.706, null 95%: −0.471 −− 0.471
non−additivity of log final mass

(ae)

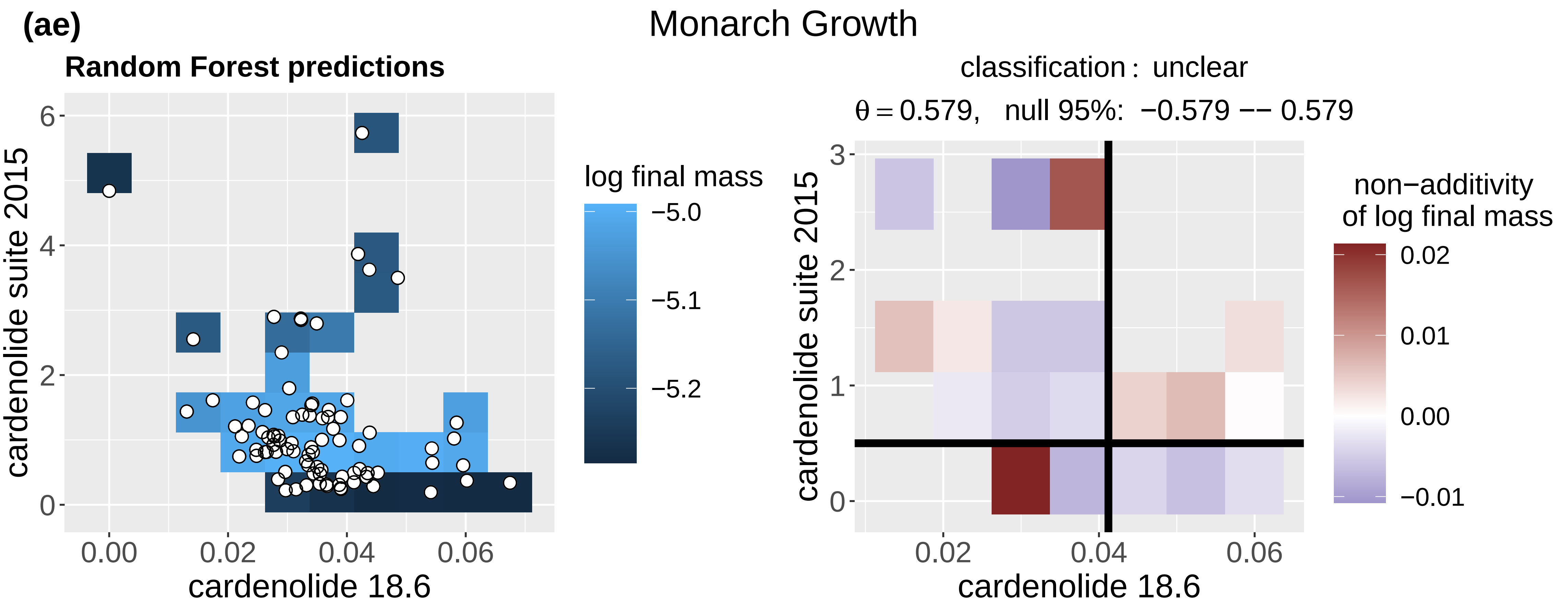
Monarch Growth
Random Forest predictions
cardenolide suite 2015
cardenolide 18.6
log final mass
−5.0
−5.1
−5.2
classification : unclear
θ = 0.579, null 95%: −0.579 −− 0.579
non−additivity of log final mass
0.02
0.01
0.00
−0.01

(af)

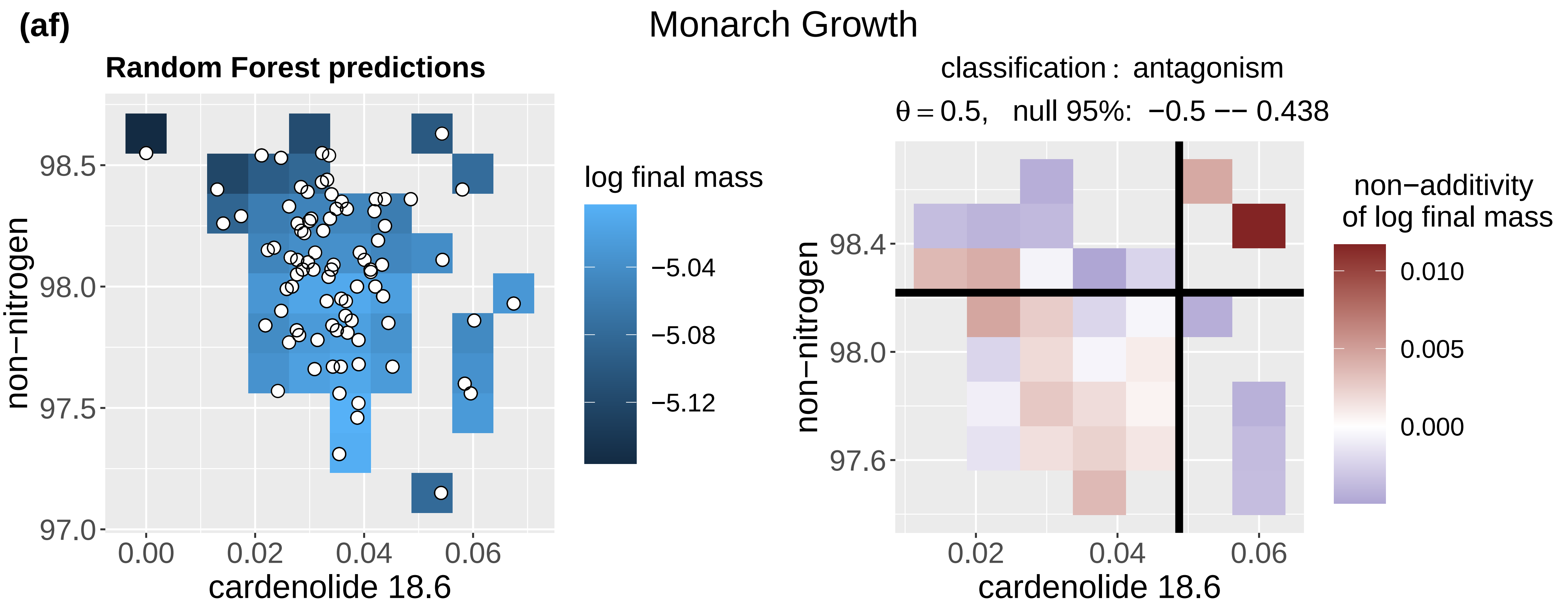
Monarch Growth
Random Forest predictions
non−nitrogen
cardenolide 18.6
log final mass
−5.04
−5.08
−5.12
classification : antagonism
θ = 0.5, null 95%: −0.5 −− 0.438
non−additivity of log final mass
0.010
0.005
0.000

(ag)

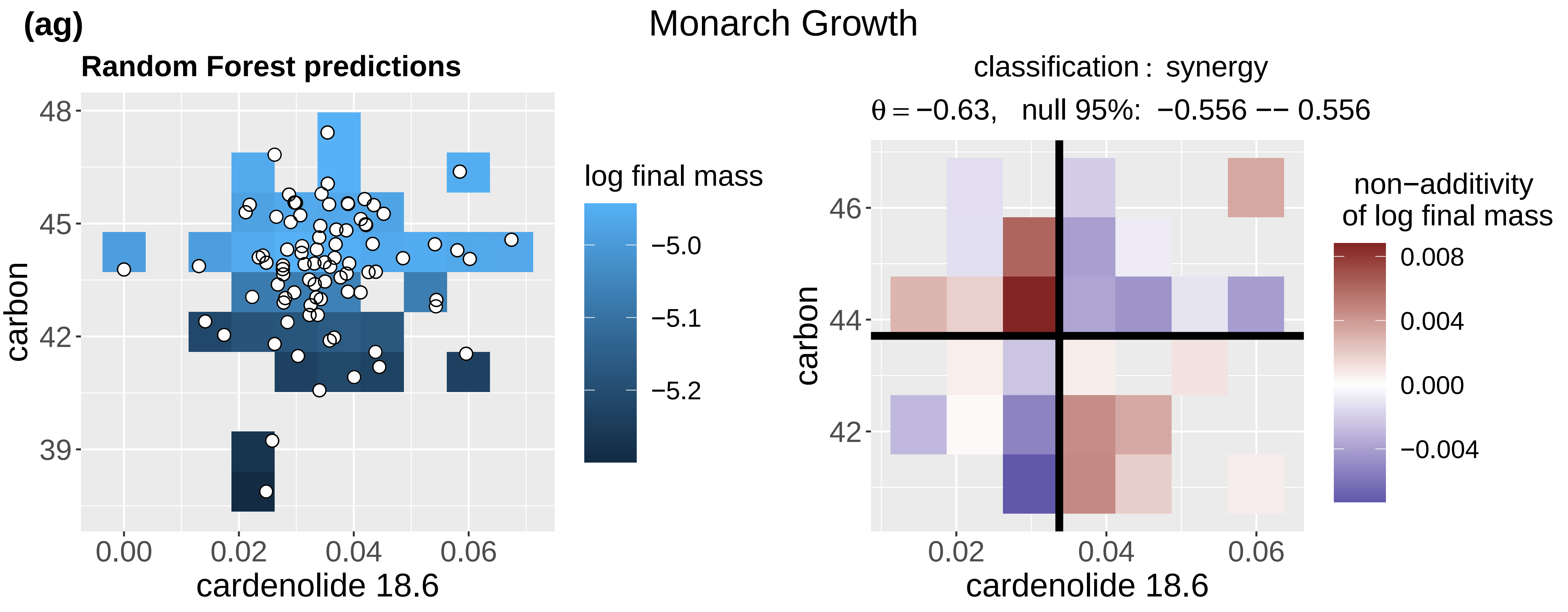
Monarch Growth
Random Forest predictions
carbon
cardenolide 18.6
log final mass
−5.0
−5.1
−5.2
classification : synergy
θ = −0.63, null 95%: −0.556 −− 0.556
non−additivity of log final mass
0.008
0.004
0.000
−0.004

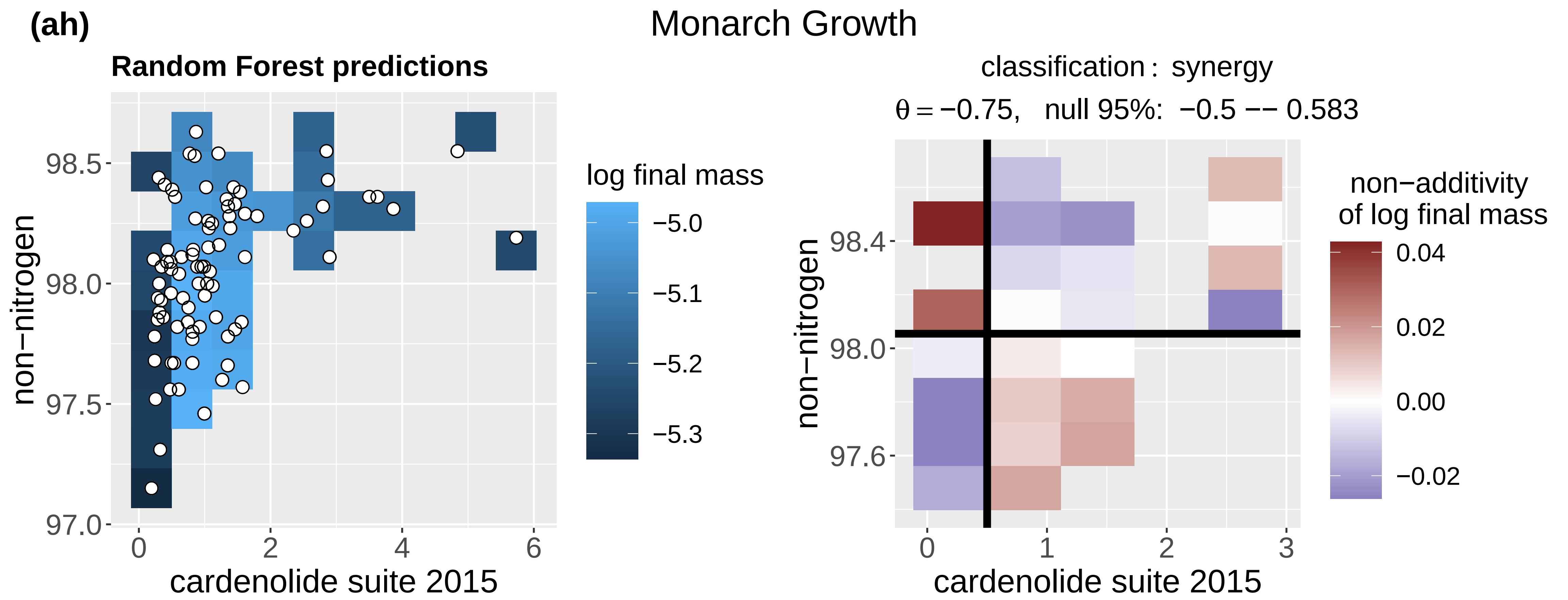
(ah)
Monarch Growth
Random Forest predictions
log final mass
non−nitrogen
cardenolide suite 2015
classification: synergy
θ = −0.75, null 95%: −0.5 −− 0.583
non−additivity
of log final mass

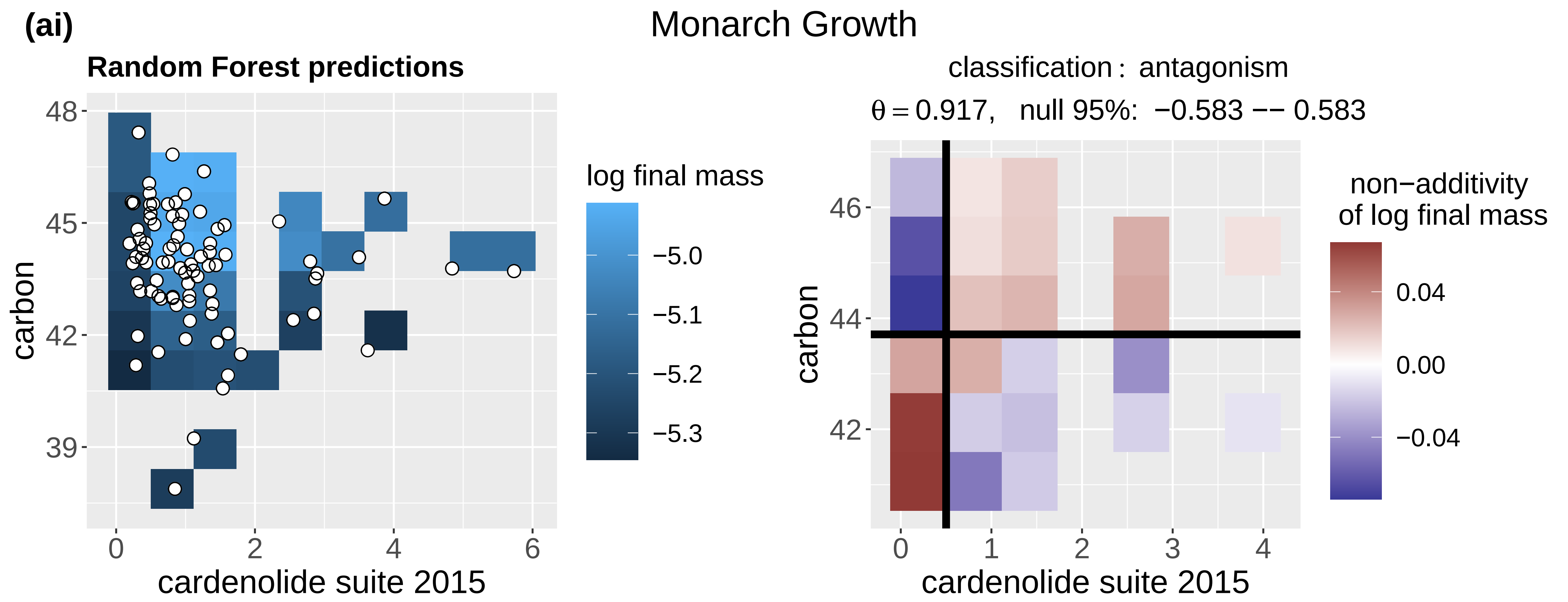
(ai)
Monarch Growth
Random Forest predictions
log final mass
carbon
cardenolide suite 2015
classification: antagonism
θ = 0.917, null 95%: −0.583 −− 0.583
non−additivity
of log final mass

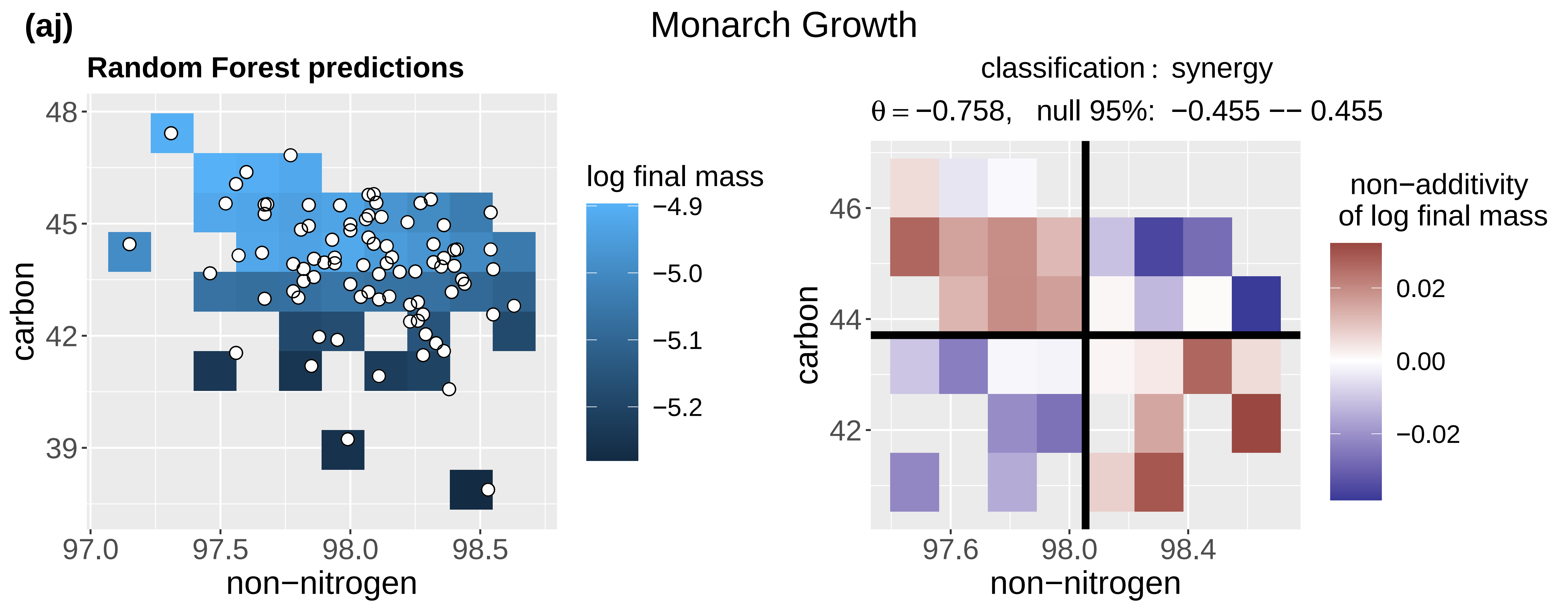
(aj)
Monarch Growth
Random Forest predictions
log final mass
carbon
non−nitrogen
classification: synergy
θ = −0.758, null 95%: −0.455 −− 0.455
non−additivity
of log final mass

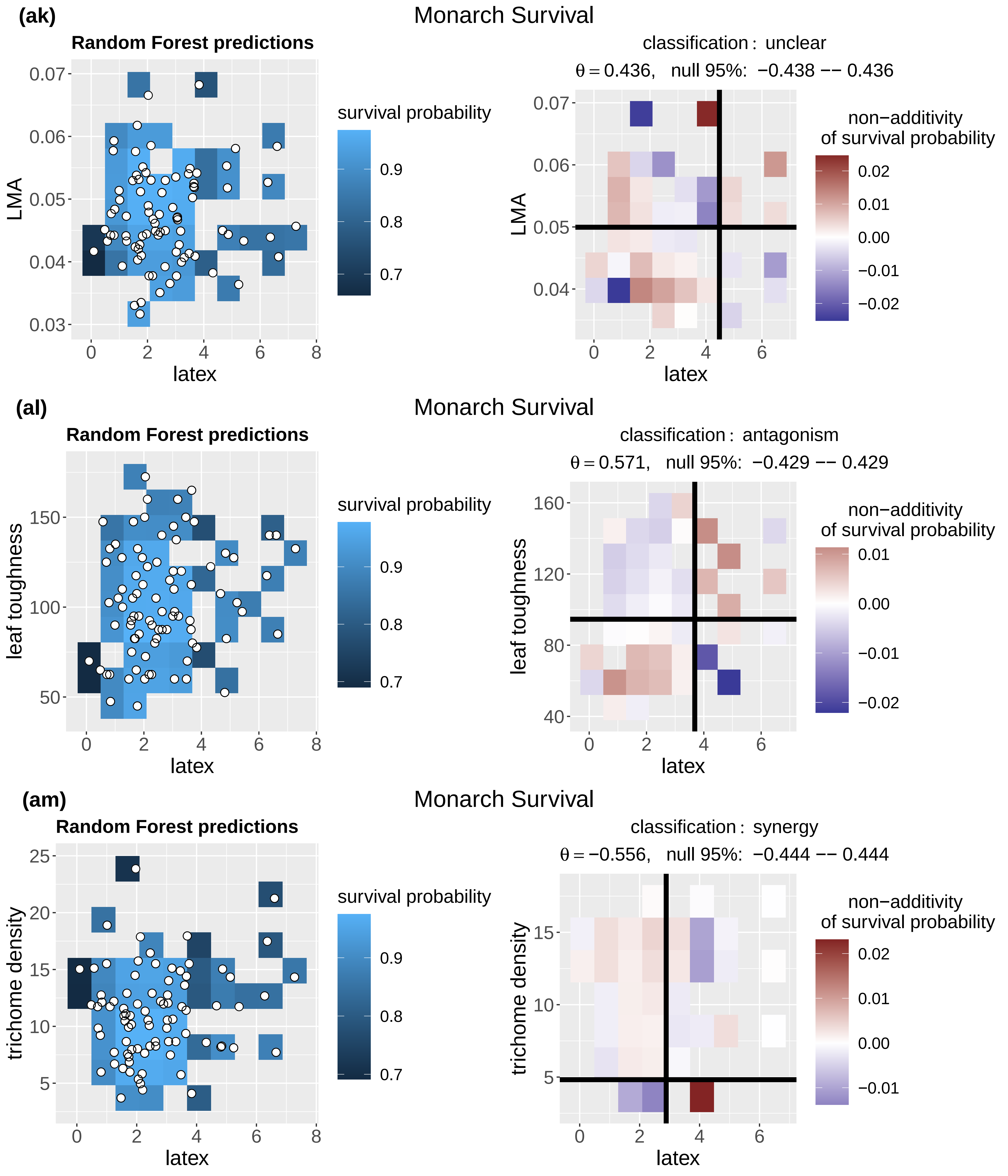

(ak)
Monarch Survival
Random Forest predictions
LMA
latex
survival probability
classification : unclear
θ = 0.436, null 95%: −0.438 −− 0.436
non−additivity
of survival probability
(al)
Monarch Survival
Random Forest predictions
leaf toughness
latex
survival probability
classification : antagonism
θ = 0.571, null 95%: −0.429 −− 0.429
non−additivity
of survival probability
(am)
Monarch Survival
Random Forest predictions
trichome density
latex
survival probability
classification : synergy
θ = −0.556, null 95%: −0.444 −− 0.444
non−additivity
of survival probability

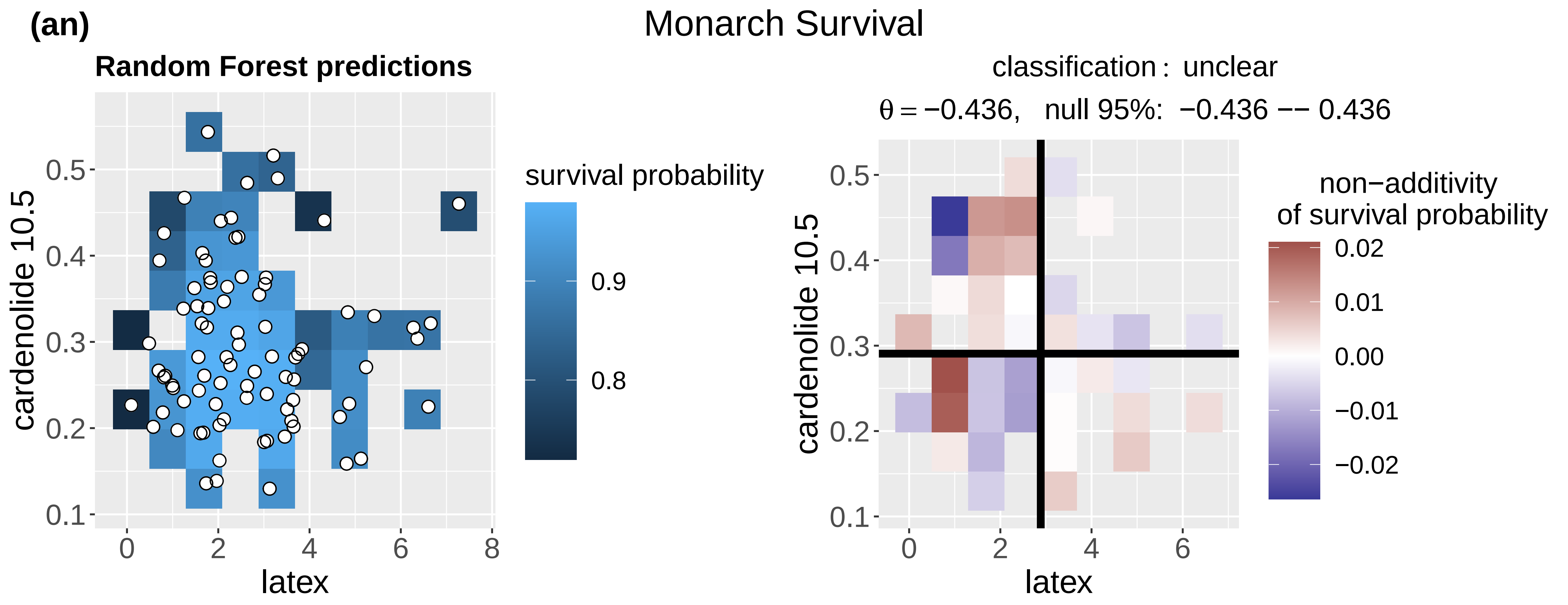
(an)
Monarch Survival
Random Forest predictions
cardenolide 10.5
latex
survival probability
0.9
0.8
classification : unclear
θ = −0.436, null 95%: −0.436 −− 0.436
non−additivity of survival probability
0.02
0.01
0.00
−0.01
−0.02

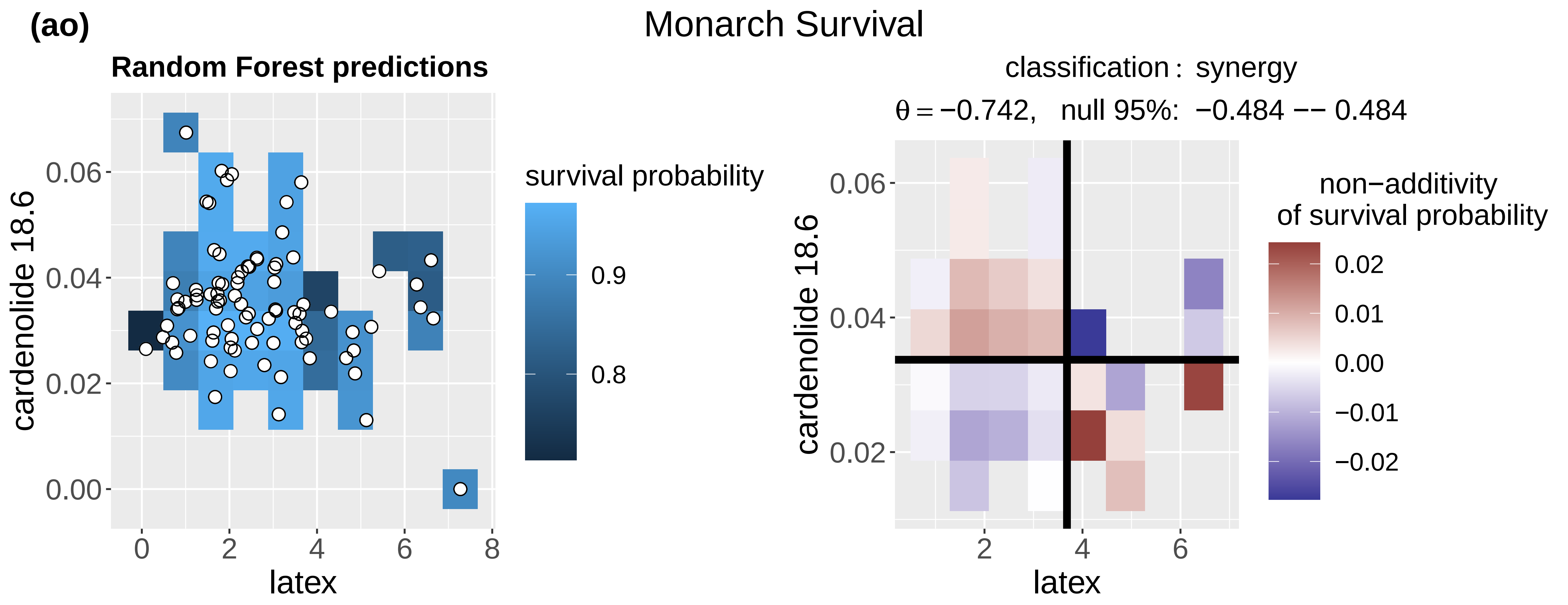
(ao)
Monarch Survival
Random Forest predictions
cardenolide 18.6
latex
survival probability
0.9
0.8
classification : synergy
θ = −0.742, null 95%: −0.484 −− 0.484
non−additivity of survival probability
0.02
0.01
0.00
−0.01
−0.02

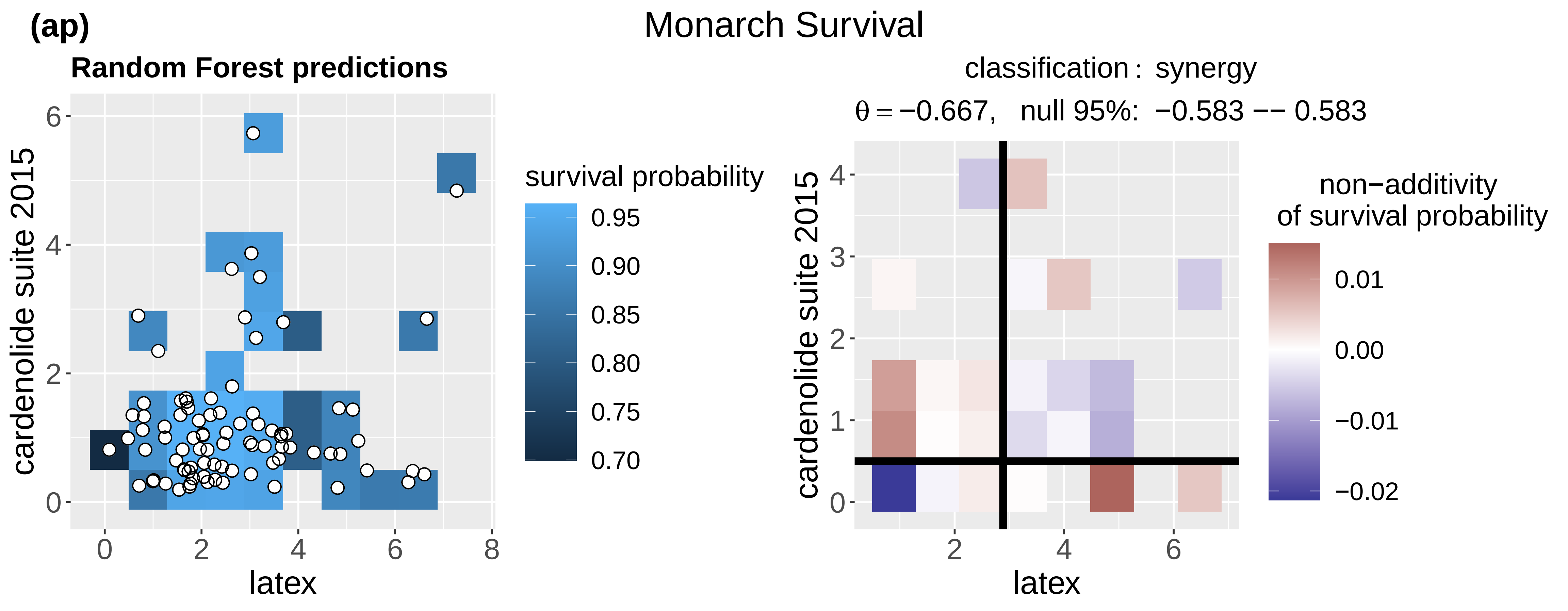
(ap)
Monarch Survival
Random Forest predictions
cardenolide suite 2015
latex
survival probability
0.95
0.90
0.85
0.80
0.75
0.70
classification : synergy
θ = −0.667, null 95%: −0.583 −− 0.583
non−additivity of survival probability
0.01
0.00
−0.01
−0.02

**(aq)**

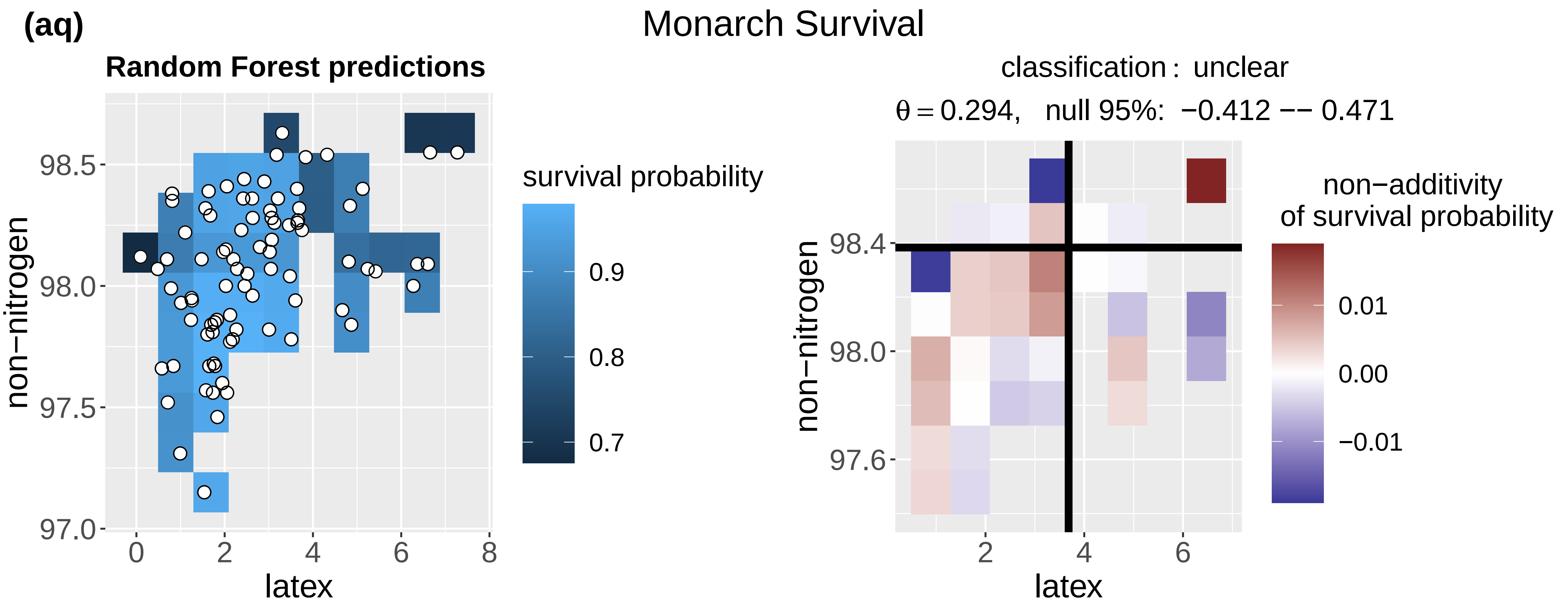
Monarch Survival
Random Forest predictions
non−nitrogen
98.5
98.0
97.5
97.0
0
2
4
6
8
latex
survival probability
0.9
0.8
0.7
classification : unclear
θ = 0.294, null 95%: −0.412 −− 0.471
non−nitrogen
98.4
98.0
97.6
2
4
6
latex
non−additivity
of survival probability
0.01
0.00
−0.01

**(ar)**

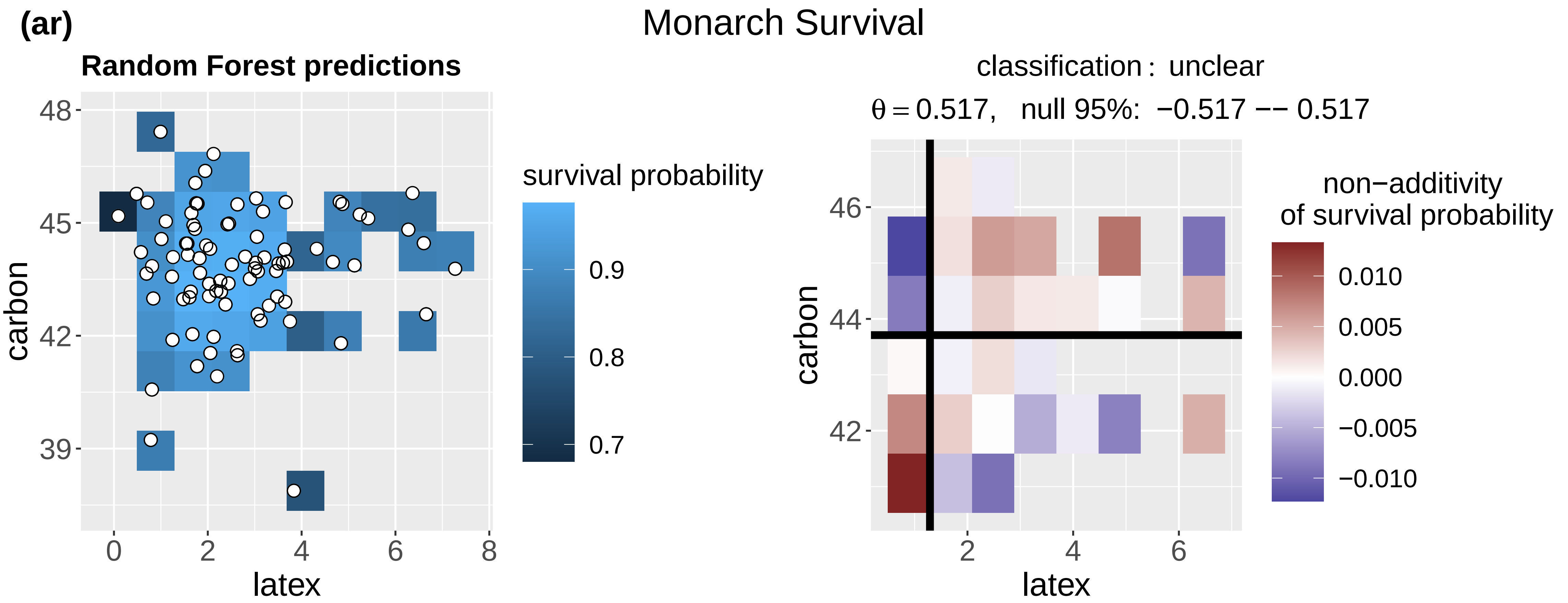
Monarch Survival
Random Forest predictions
carbon
48
45
42
39
0
2
4
6
8
latex
survival probability
0.9
0.8
0.7
classification : unclear
θ = 0.517, null 95%: −0.517 −− 0.517
carbon
46
44
42
2
4
6
latex
non−additivity
of survival probability
0.010
0.005
0.000
−0.005
−0.010

**(as)**

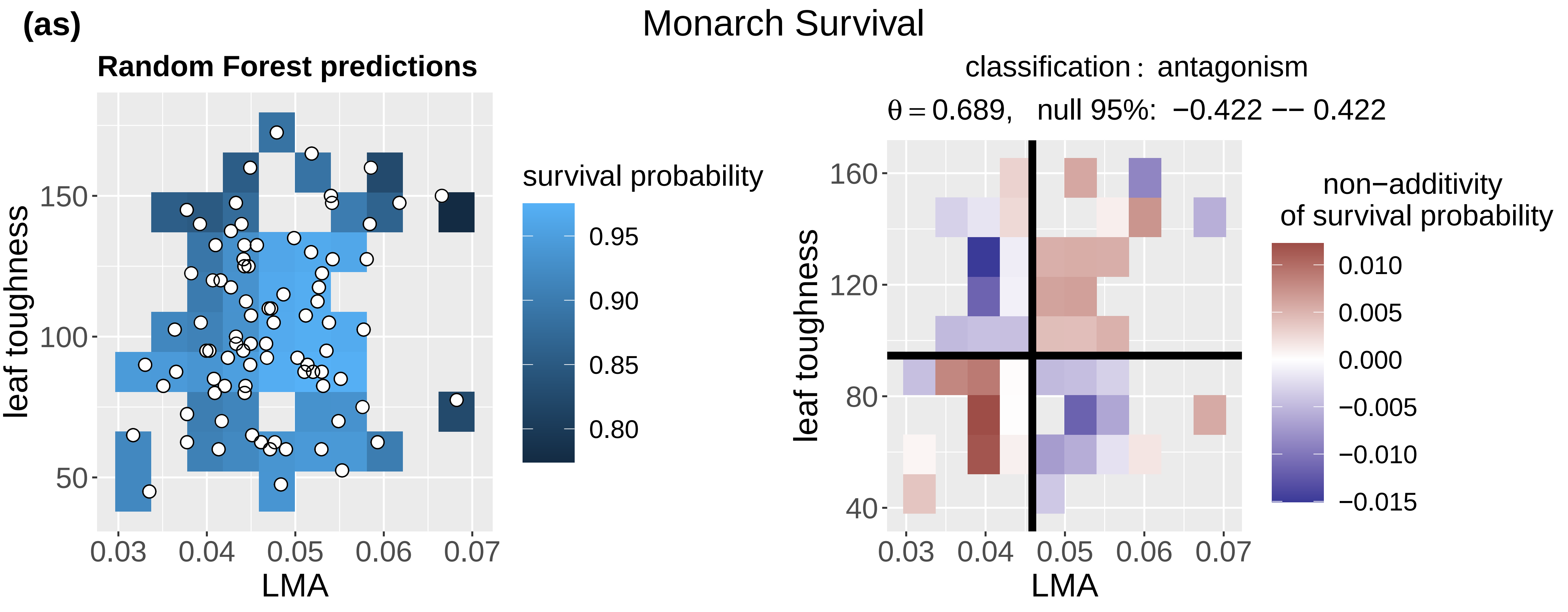
Monarch Survival
Random Forest predictions
leaf toughness
150
100
50
0.03
0.04
0.05
0.06
0.07
LMA
survival probability
0.95
0.90
0.85
0.80
classification : antagonism
θ = 0.689, null 95%: −0.422 −− 0.422
leaf toughness
160
120
80
40
0.03
0.04
0.05
0.06
0.07
LMA
non−additivity
of survival probability
0.010
0.005
0.000
−0.005
−0.010
−0.015

(at)

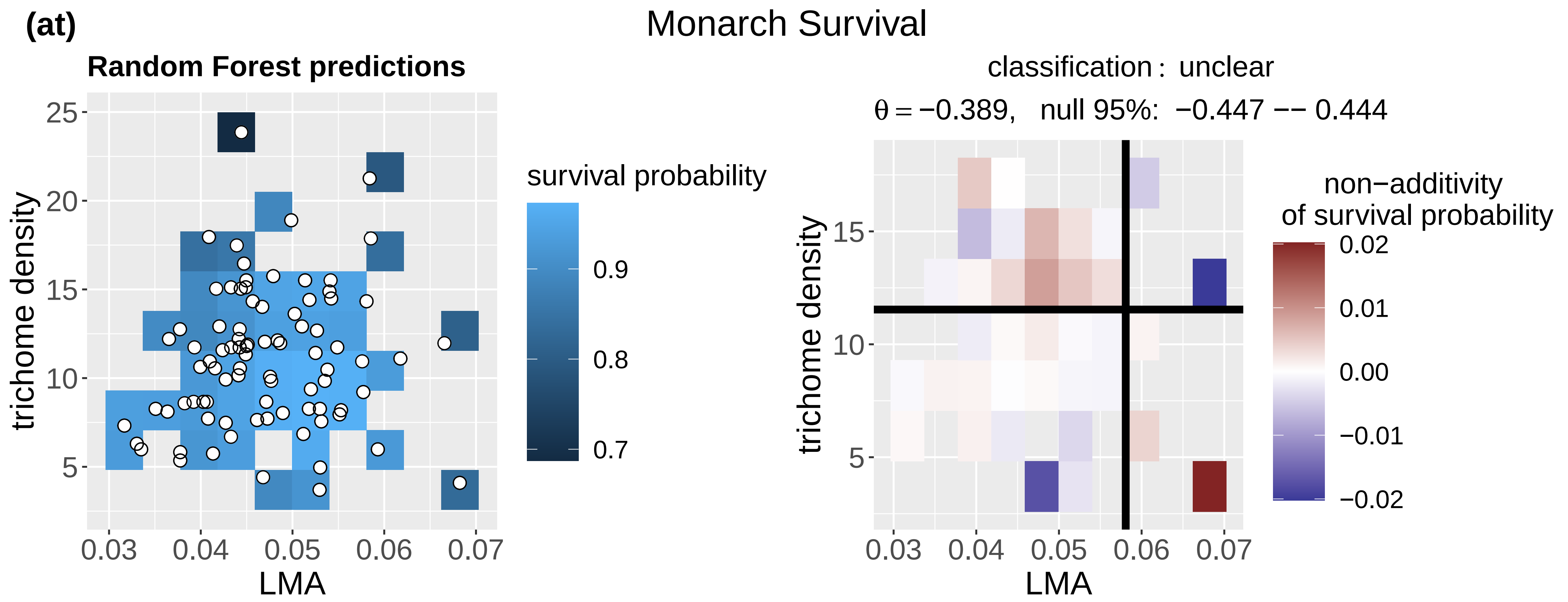
Monarch Survival
Random Forest predictions
survival probability
trichome density
LMA
classification : unclear
θ = −0.389, null 95%: −0.447 −− 0.444
non−additivity
of survival probability
trichome density
LMA

(au)

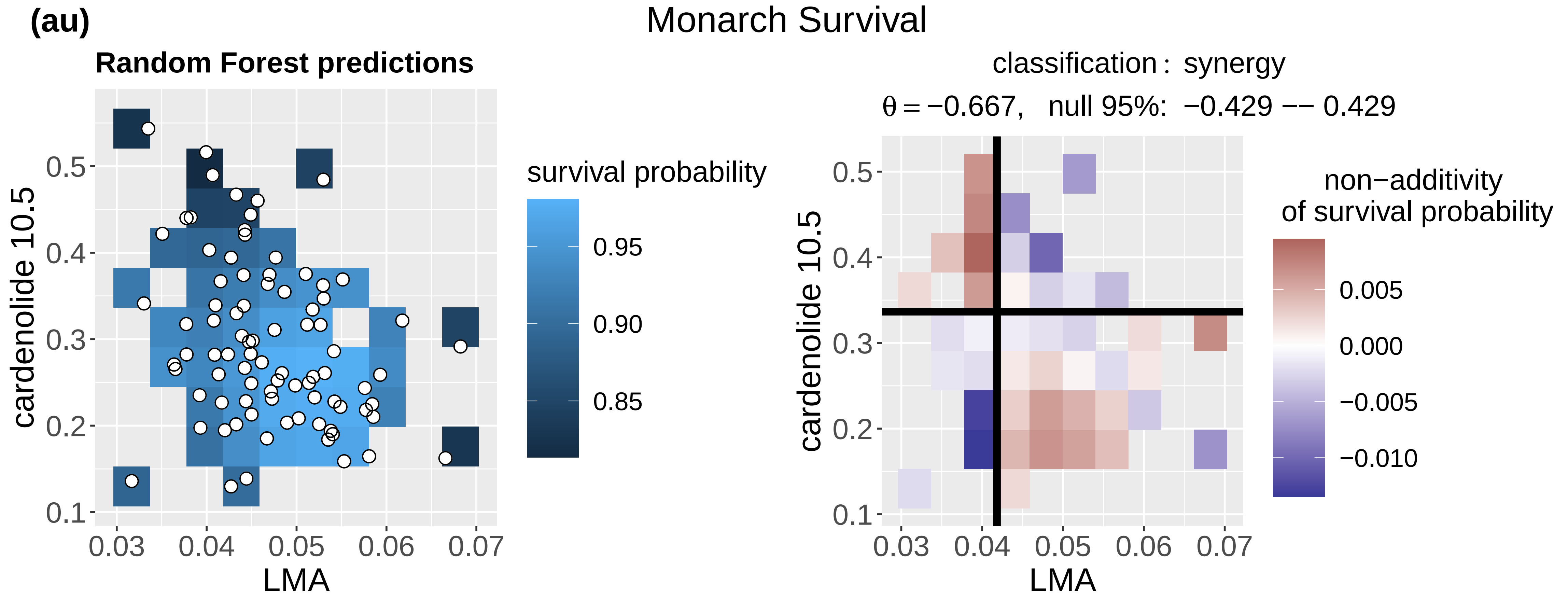
Monarch Survival
Random Forest predictions
survival probability
cardenolide 10.5
LMA
classification : synergy
θ = −0.667, null 95%: −0.429 −− 0.429
non−additivity
of survival probability
cardenolide 10.5
LMA

(av)

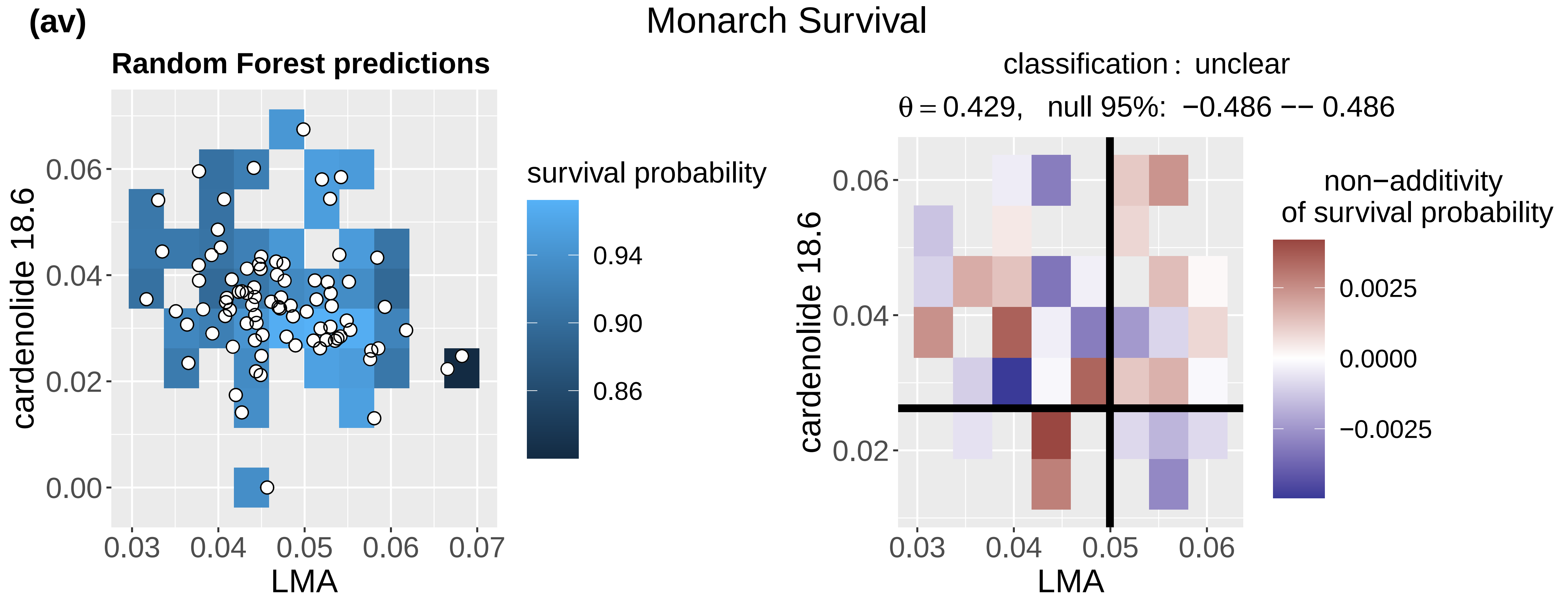
Monarch Survival
Random Forest predictions
survival probability
cardenolide 18.6
LMA
classification : unclear
θ = 0.429, null 95%: −0.486 −− 0.486
non−additivity
of survival probability
cardenolide 18.6
LMA

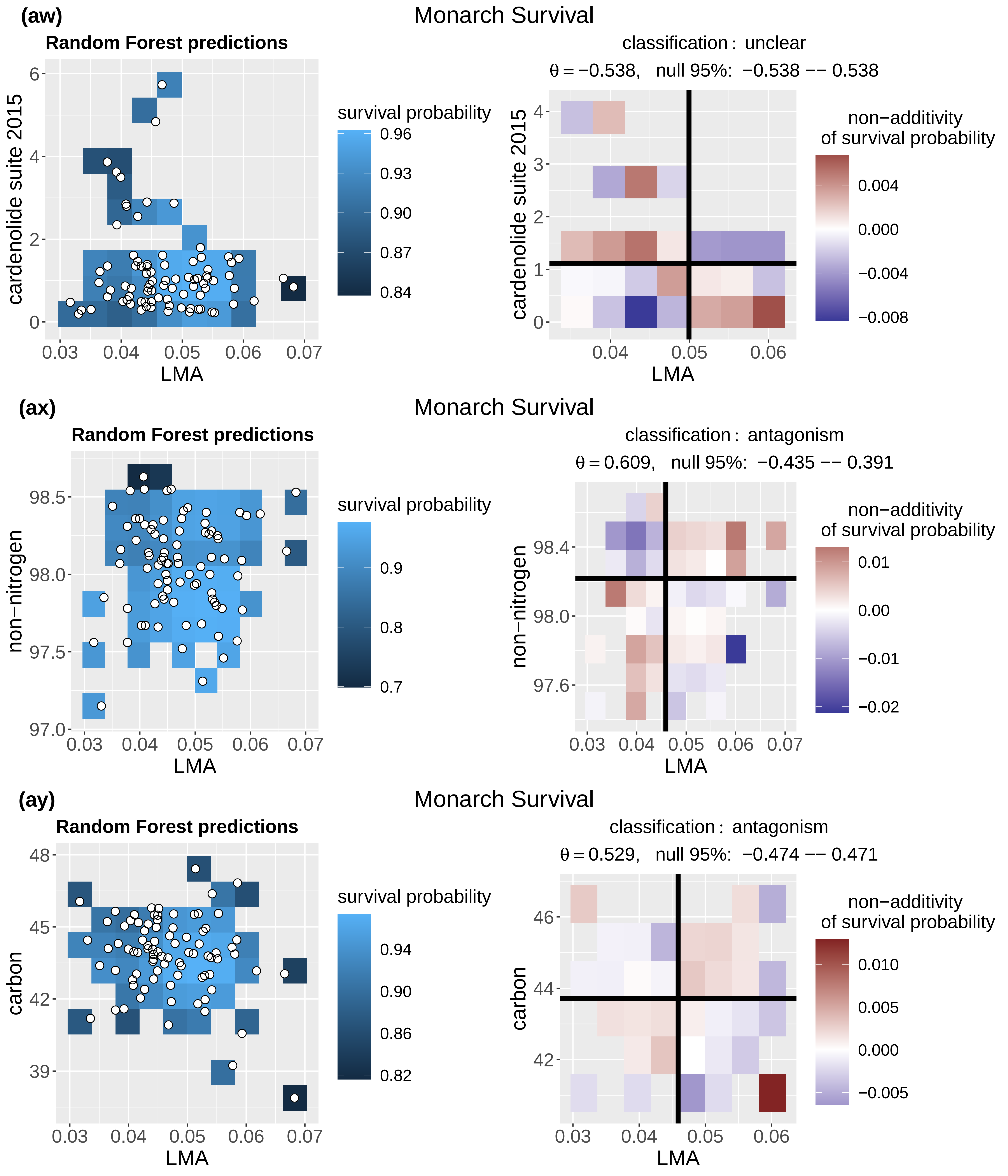
(aw)
Monarch Survival
Random Forest predictions
cardenolide suite 2015
LMA
survival probability
0.96
0.93
0.90
0.87
0.84
classification : unclear
θ = −0.538,   null 95%:  −0.538 −− 0.538
non−additivity
of survival probability
0.004
0.000
−0.004
−0.008
(ax)
Monarch Survival
Random Forest predictions
non−nitrogen
LMA
survival probability
0.9
0.8
0.7
classification : antagonism
θ = 0.609,   null 95%:  −0.435 −− 0.391
non−additivity
of survival probability
0.01
0.00
−0.01
−0.02
(ay)
Monarch Survival
Random Forest predictions
carbon
LMA
survival probability
0.94
0.90
0.86
0.82
classification : antagonism
θ = 0.529,   null 95%:  −0.474 −− 0.471
non−additivity
of survival probability
0.010
0.005
0.000
−0.005

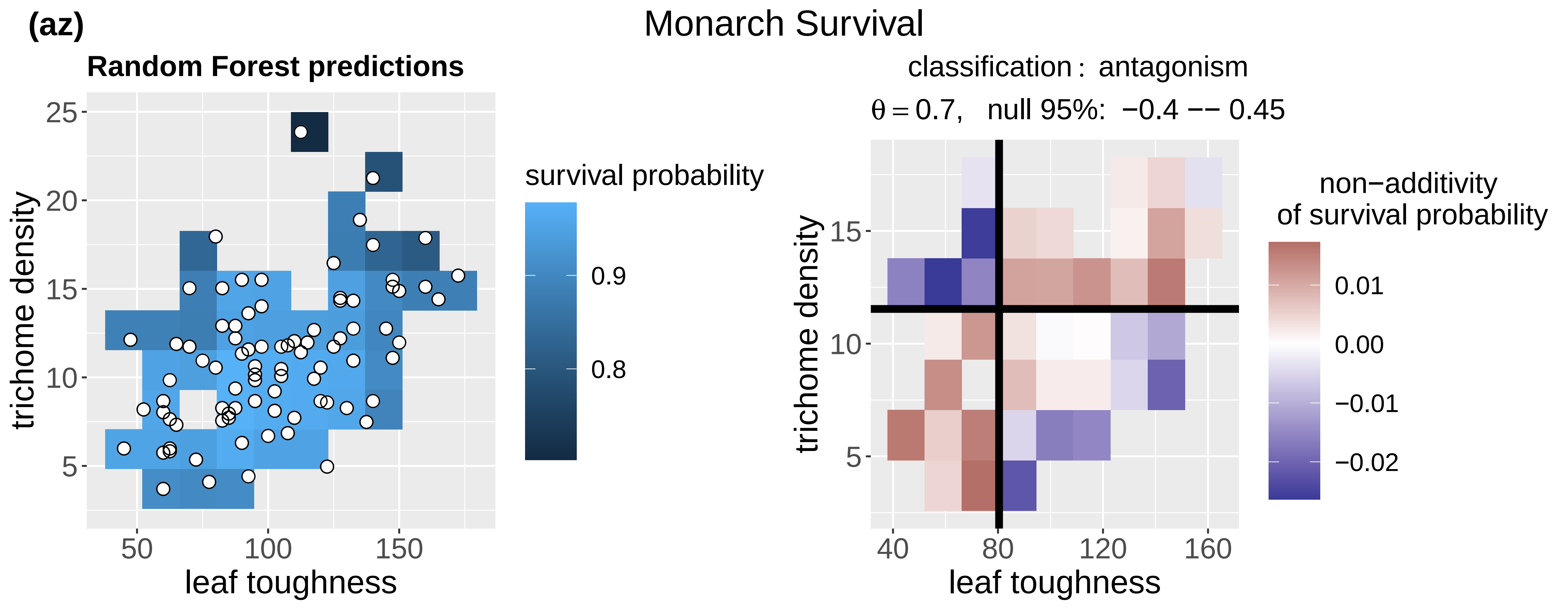
(az)
Monarch Survival
Random Forest predictions
trichome density
leaf toughness
survival probability
0.9
0.8
classification : antagonism
θ = 0.7, null 95%: −0.4 −− 0.45
non−additivity
of survival probability
0.01
0.00
−0.01
−0.02

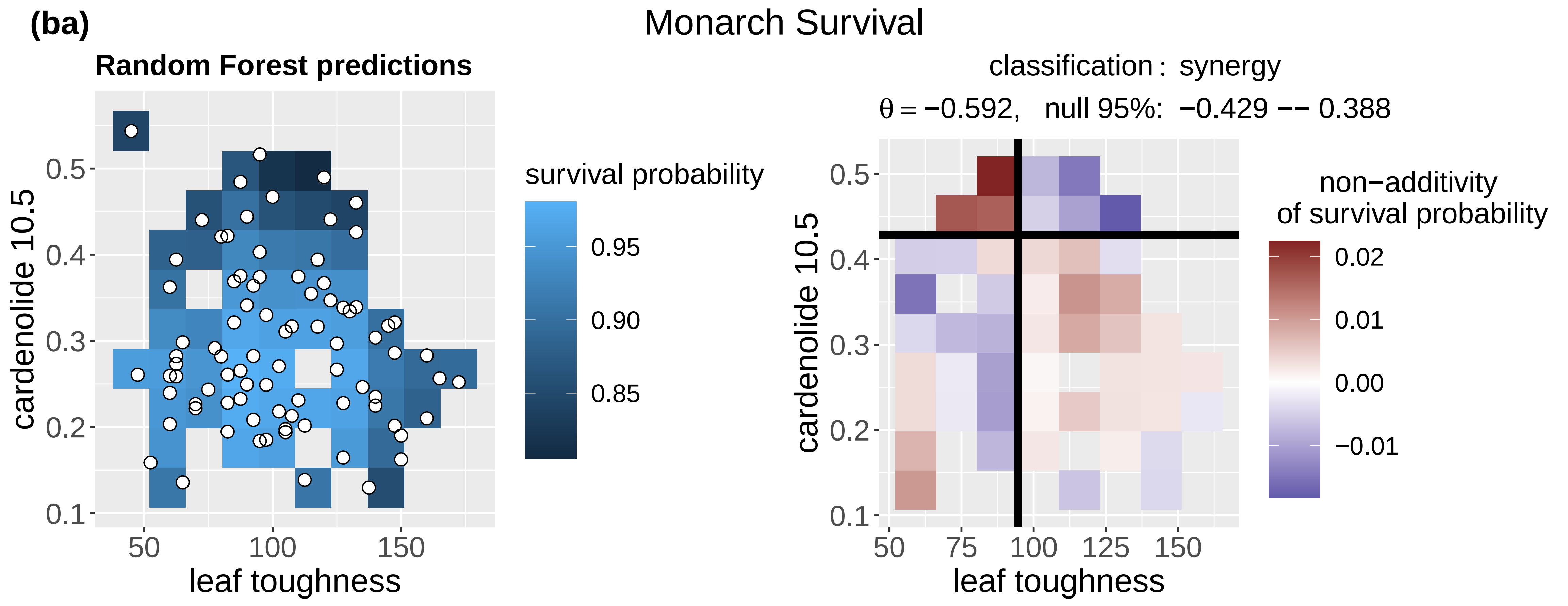
(ba)
Monarch Survival
Random Forest predictions
cardenolide 10.5
leaf toughness
survival probability
0.95
0.90
0.85
classification : synergy
θ = −0.592, null 95%: −0.429 −− 0.388
non−additivity
of survival probability
0.02
0.01
0.00
−0.01

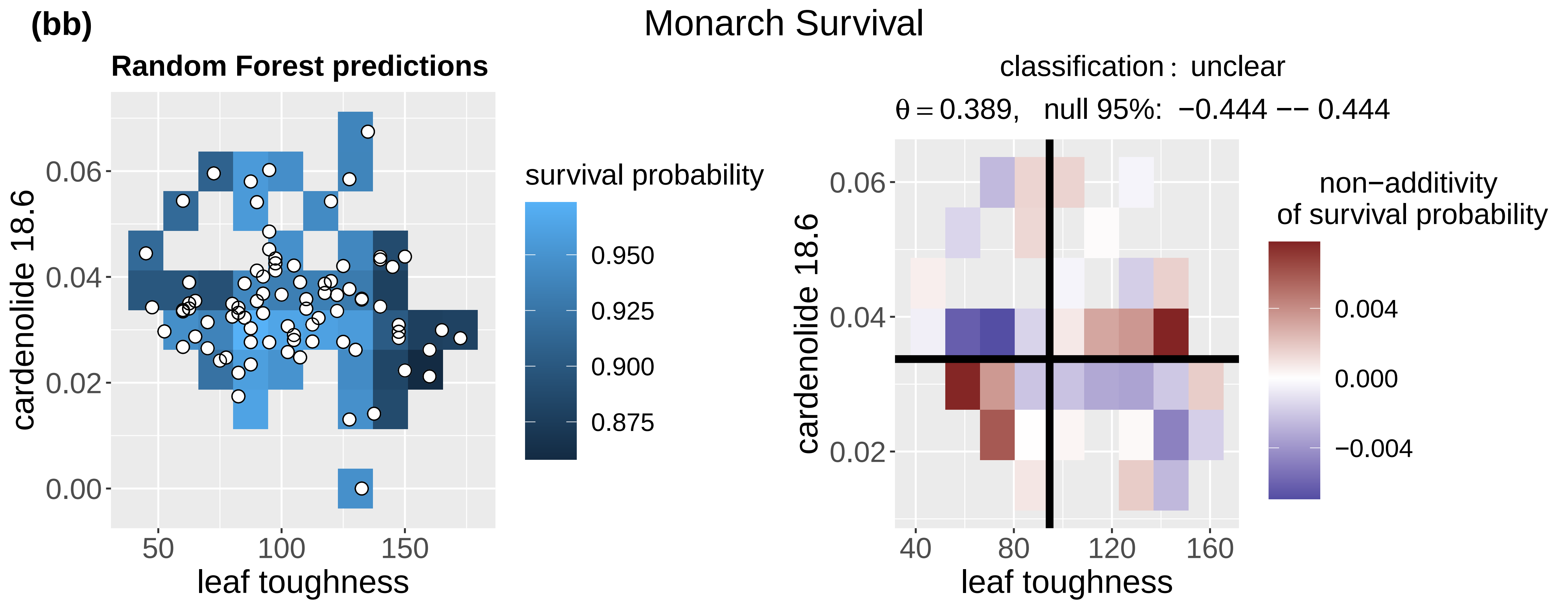
(bb)
Monarch Survival
Random Forest predictions
cardenolide 18.6
leaf toughness
survival probability
0.950
0.925
0.900
0.875
classification : unclear
θ = 0.389, null 95%: −0.444 −− 0.444
non−additivity
of survival probability
0.004
0.000
−0.004

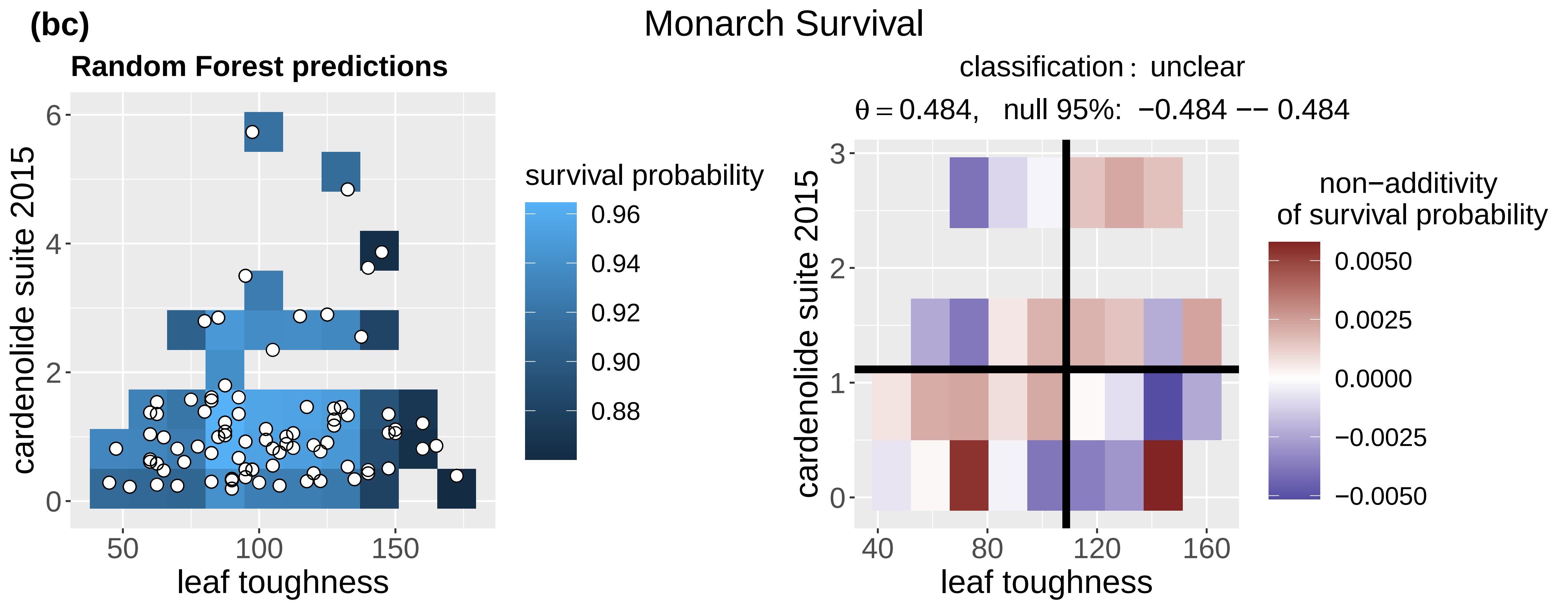
(bc)
Monarch Survival
Random Forest predictions
cardenolide suite 2015
0
2
4
6
50
100
150
leaf toughness
survival probability
0.96
0.94
0.92
0.90
0.88
classification : unclear
θ = 0.484, null 95%: −0.484 −− 0.484
cardenolide suite 2015
0
1
2
3
40
80
120
160
leaf toughness
non−additivity
of survival probability
0.0050
0.0025
0.0000
−0.0025
−0.0050

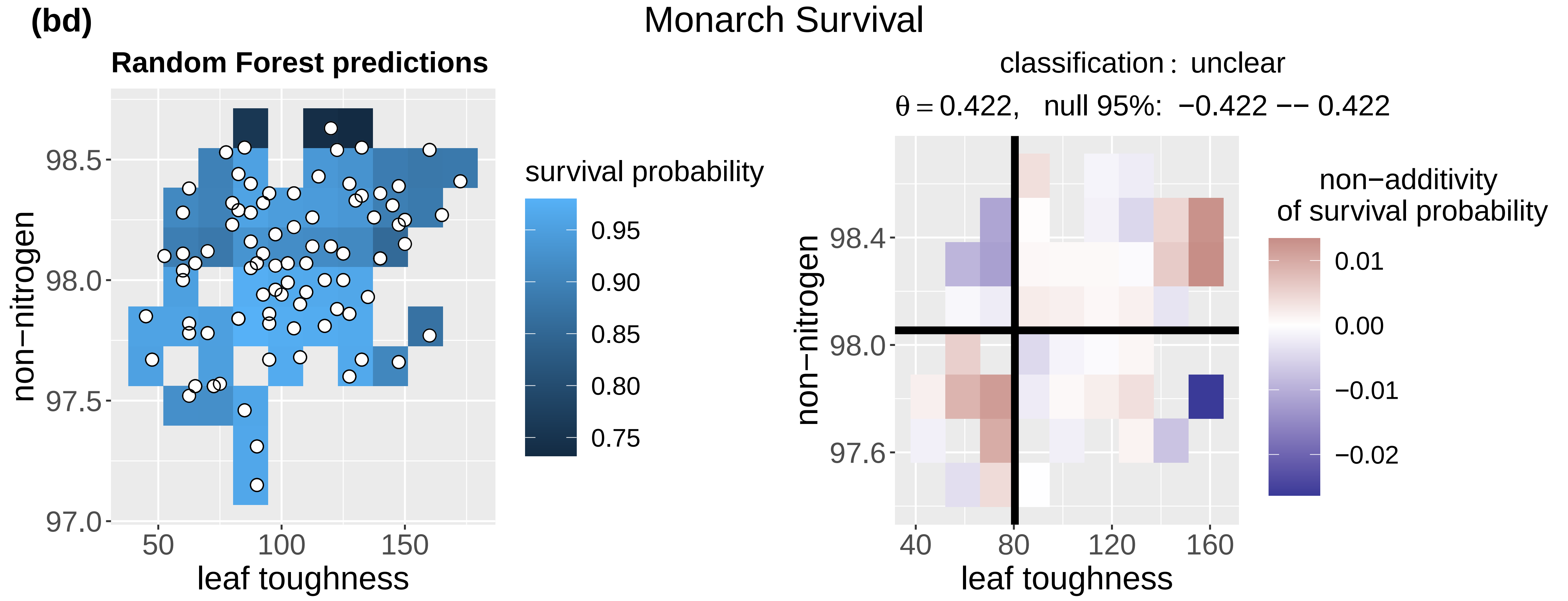
(bd)
Monarch Survival
Random Forest predictions
non−nitrogen
97.0
97.5
98.0
98.5
50
100
150
leaf toughness
survival probability
0.95
0.90
0.85
0.80
0.75
classification : unclear
θ = 0.422, null 95%: −0.422 −− 0.422
non−nitrogen
97.6
98.0
98.4
40
80
120
160
leaf toughness
non−additivity
of survival probability
0.01
0.00
−0.01
−0.02

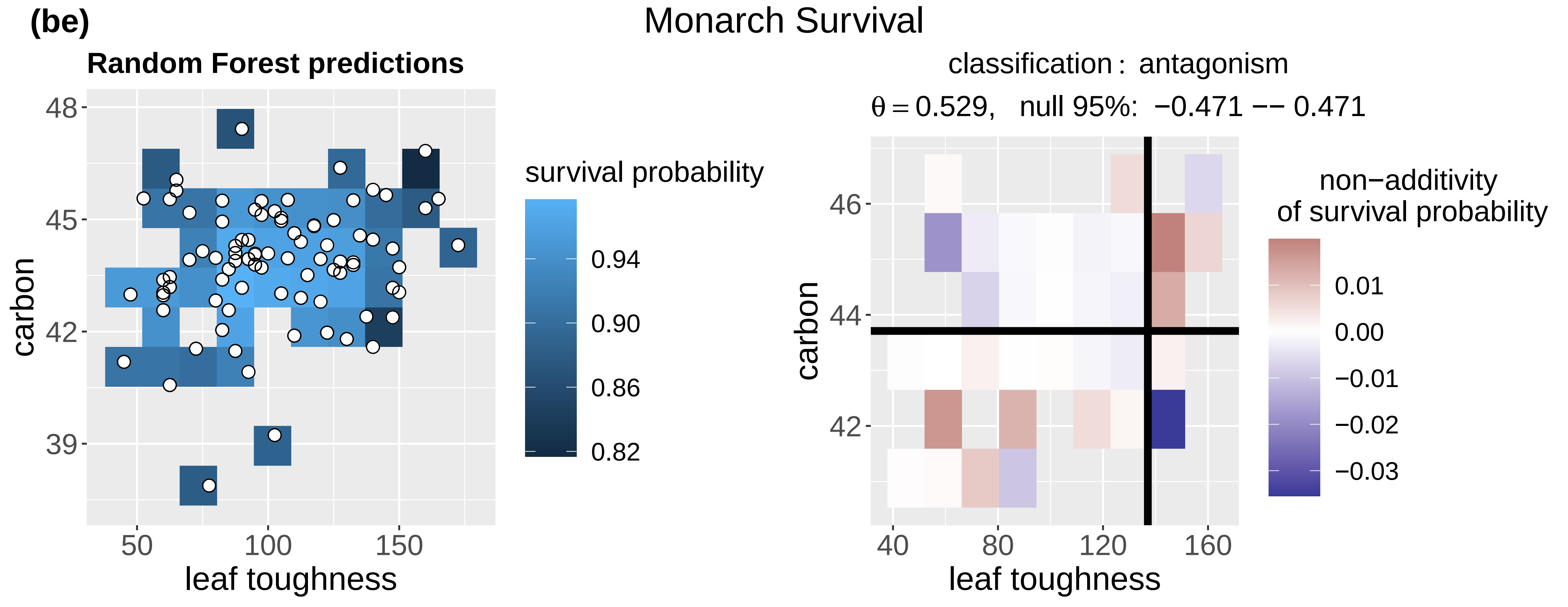
(be)
Monarch Survival
Random Forest predictions
carbon
39
42
45
48
50
100
150
leaf toughness
survival probability
0.94
0.90
0.86
0.82
classification : antagonism
θ = 0.529, null 95%: −0.471 −− 0.471
carbon
42
44
46
40
80
120
160
leaf toughness
non−additivity
of survival probability
0.01
0.00
−0.01
−0.02
−0.03

(bf)

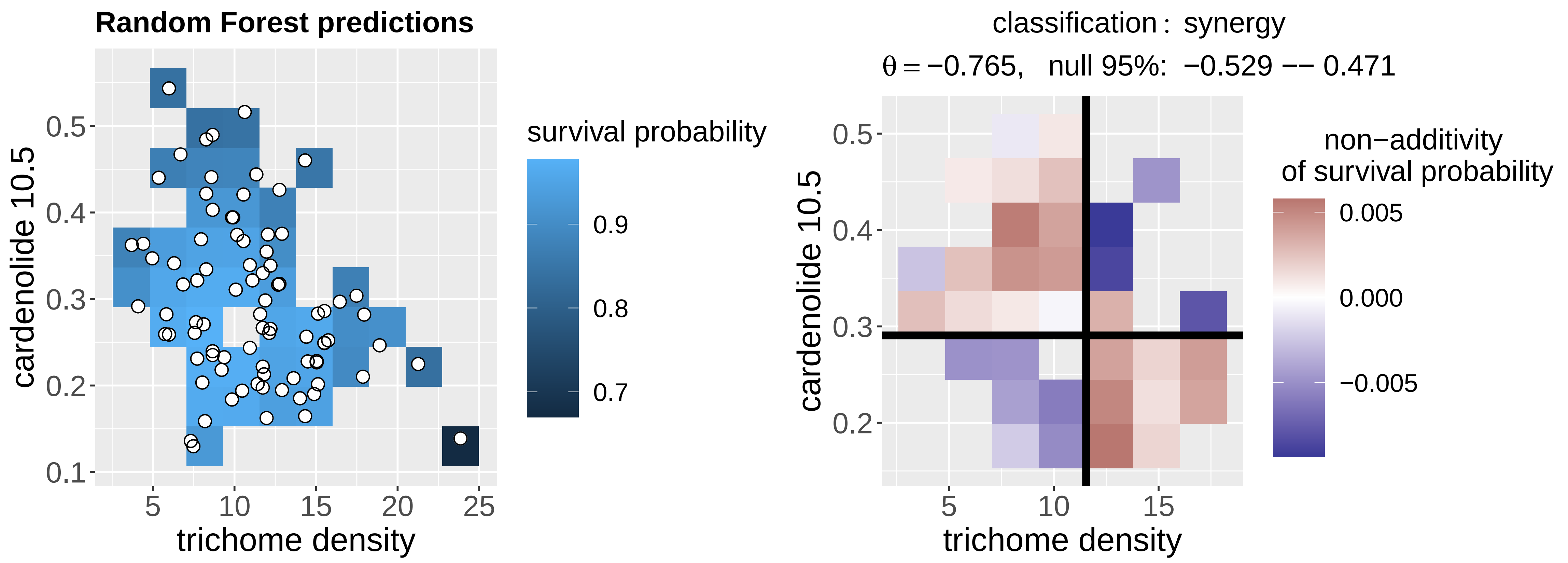
Monarch Survival
Random Forest predictions
cardenolide 10.5
0.1
0.2
0.3
0.4
0.5
5
10
15
20
25
trichome density
survival probability
0.9
0.8
0.7
classification : synergy
θ = −0.765, null 95%: −0.529 −− 0.471
cardenolide 10.5
0.2
0.3
0.4
0.5
5
10
15
trichome density
non−additivity
of survival probability
0.005
0.000
−0.005

(bg)

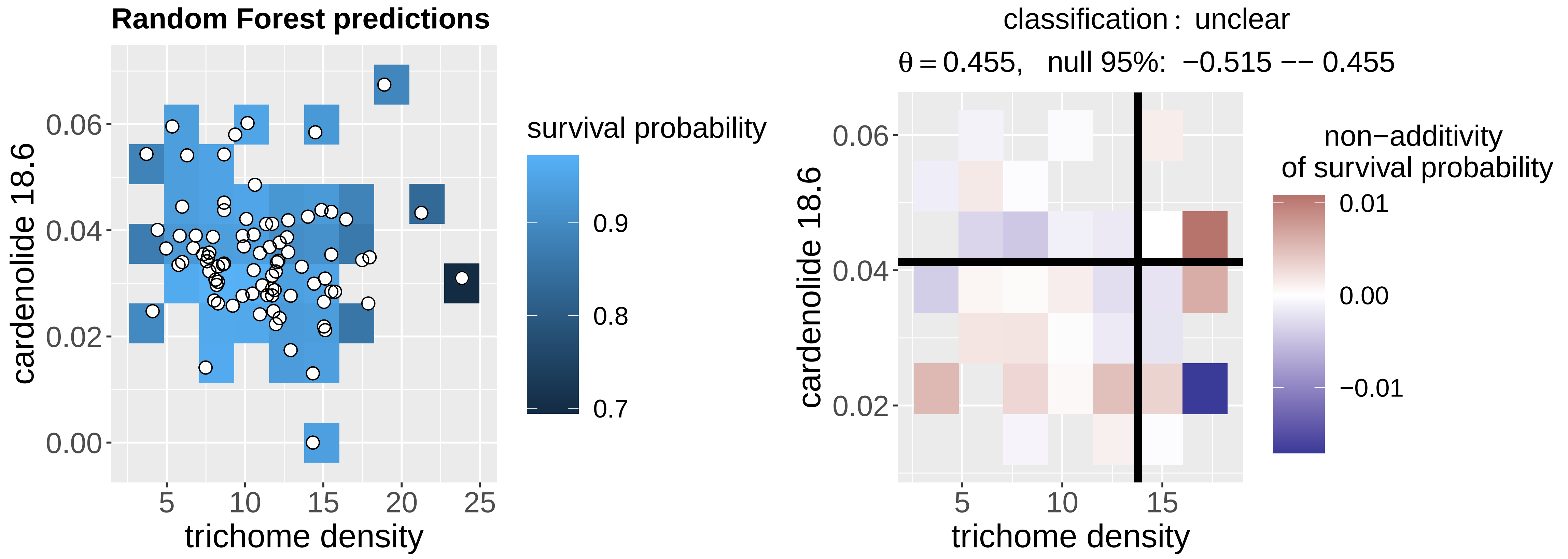
Monarch Survival
Random Forest predictions
cardenolide 18.6
0.00
0.02
0.04
0.06
5
10
15
20
25
trichome density
survival probability
0.9
0.8
0.7
classification : unclear
θ = 0.455, null 95%: −0.515 −− 0.455
cardenolide 18.6
0.02
0.04
0.06
5
10
15
trichome density
non−additivity
of survival probability
0.01
0.00
−0.01

(bh)

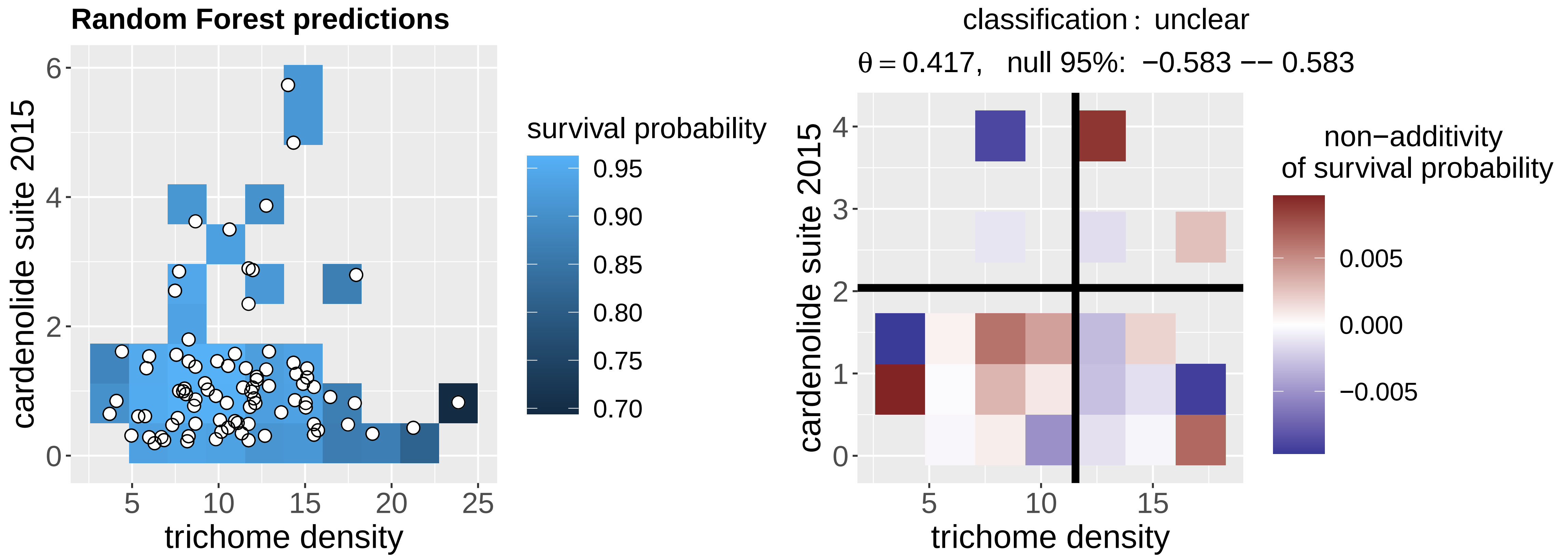
Monarch Survival
Random Forest predictions
cardenolide suite 2015
0
2
4
6
5
10
15
20
25
trichome density
survival probability
0.95
0.90
0.85
0.80
0.75
0.70
classification : unclear
θ = 0.417, null 95%: −0.583 −− 0.583
cardenolide suite 2015
0
1
2
3
4
5
10
15
trichome density
non−additivity
of survival probability
0.005
0.000
−0.005

**(bi)**

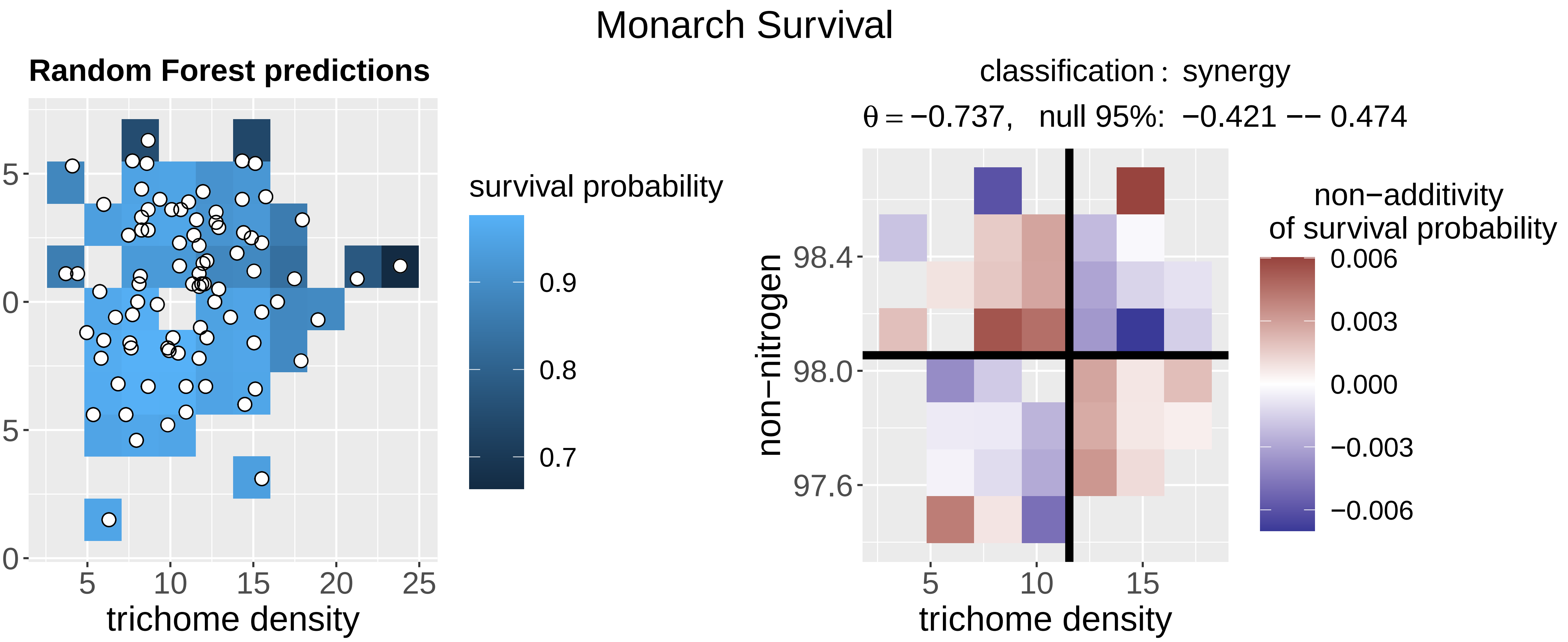
Monarch Survival
Random Forest predictions
non−nitrogen
trichome density
survival probability
0.9
0.8
0.7
classification : synergy
θ = −0.737, null 95%: −0.421 −− 0.474
non−additivity
of survival probability
0.006
0.003
0.000
−0.003
−0.006

**(bj)**

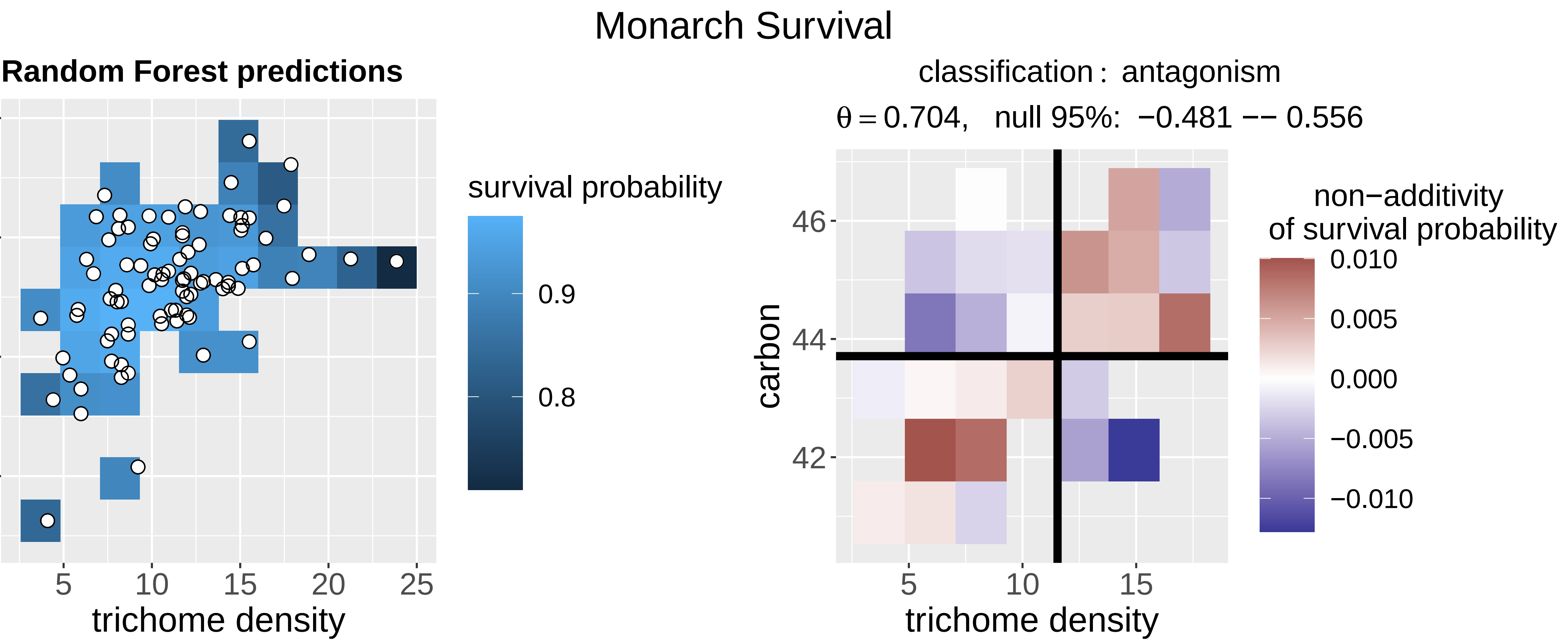
Monarch Survival
Random Forest predictions
carbon
trichome density
survival probability
0.9
0.8
classification : antagonism
θ = 0.704, null 95%: −0.481 −− 0.556
non−additivity
of survival probability
0.010
0.005
0.000
−0.005
−0.010

**(bk)**

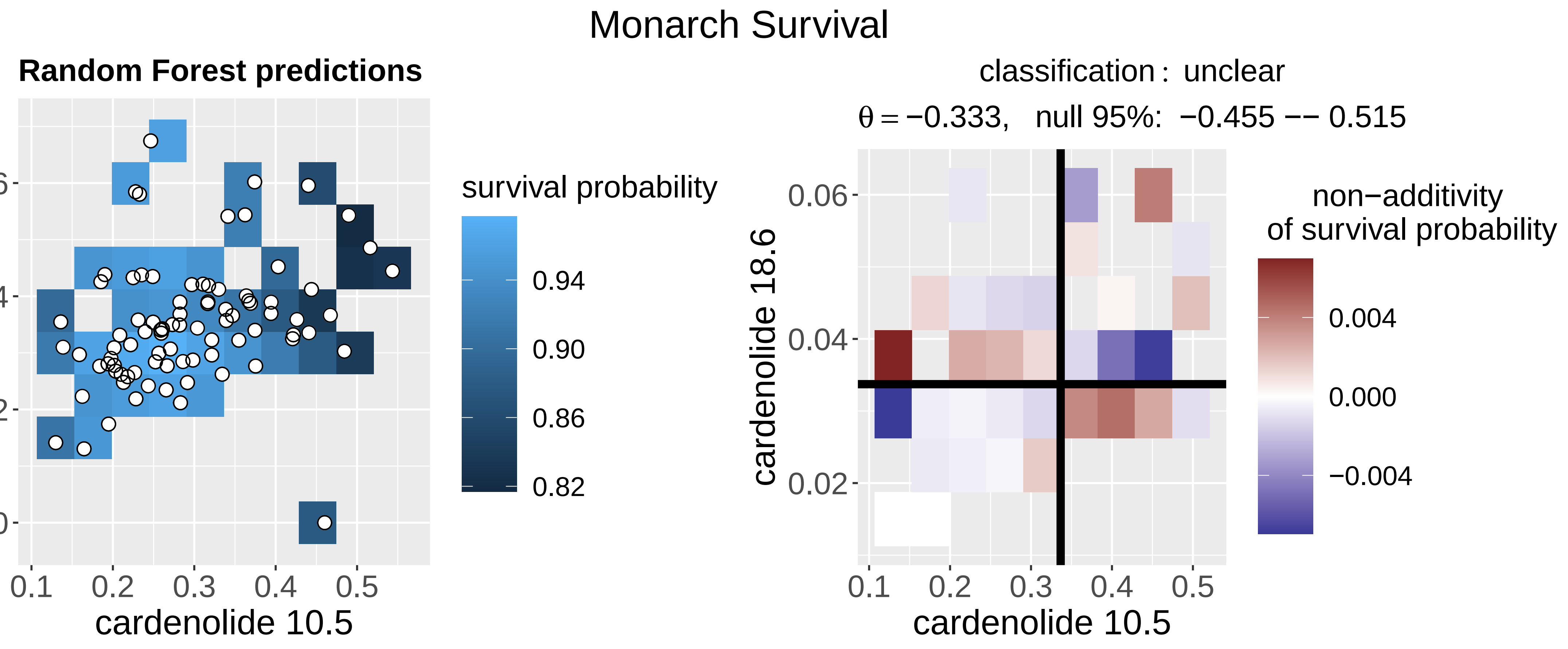
Monarch Survival
Random Forest predictions
cardenolide 18.6
cardenolide 10.5
survival probability
0.94
0.90
0.86
0.82
classification : unclear
θ = −0.333, null 95%: −0.455 −− 0.515
non−additivity
of survival probability
0.004
0.000
−0.004

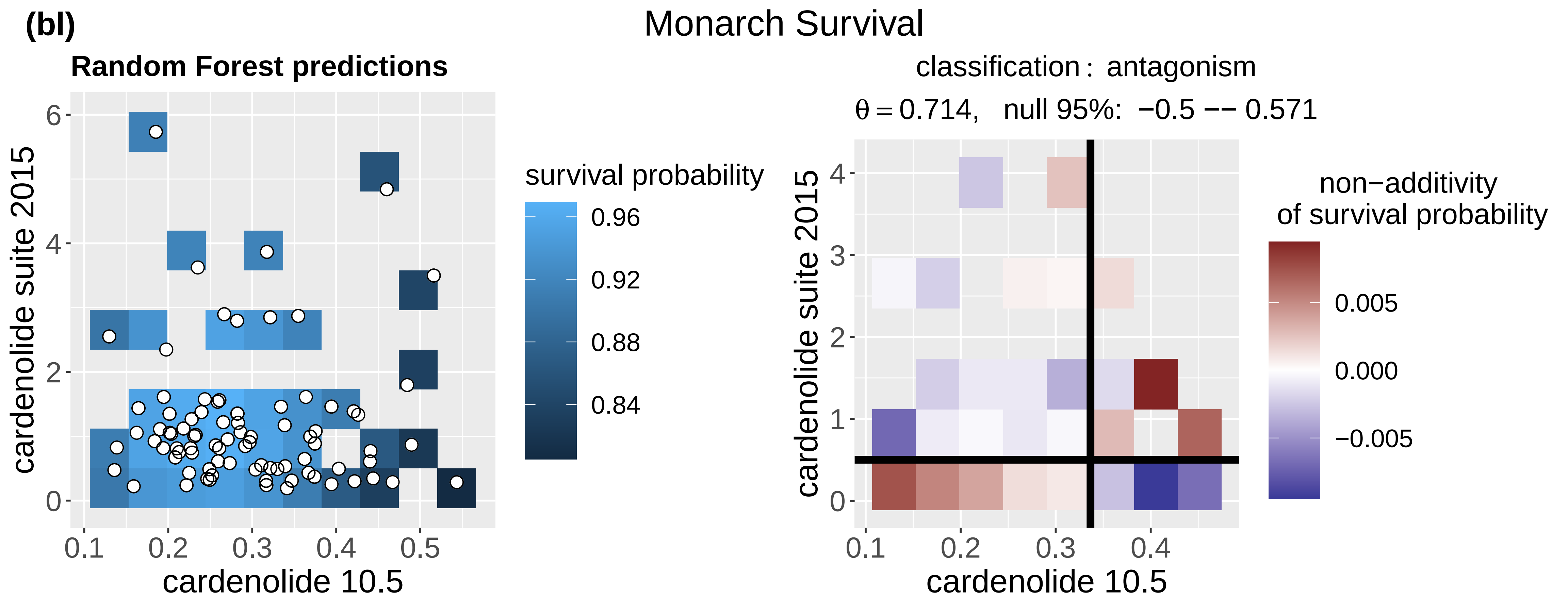
(bl)
Monarch Survival
Random Forest predictions
survival probability
0.96
0.92
0.88
0.84
cardenolide suite 2015
cardenolide 10.5
0.1 0.2 0.3 0.4 0.5
0 2 4 6
classification: antagonism
θ = 0.714, null 95%: −0.5 −− 0.571
non−additivity
of survival probability
0.005
0.000
−0.005
0 1 2 3 4
0.1 0.2 0.3 0.4

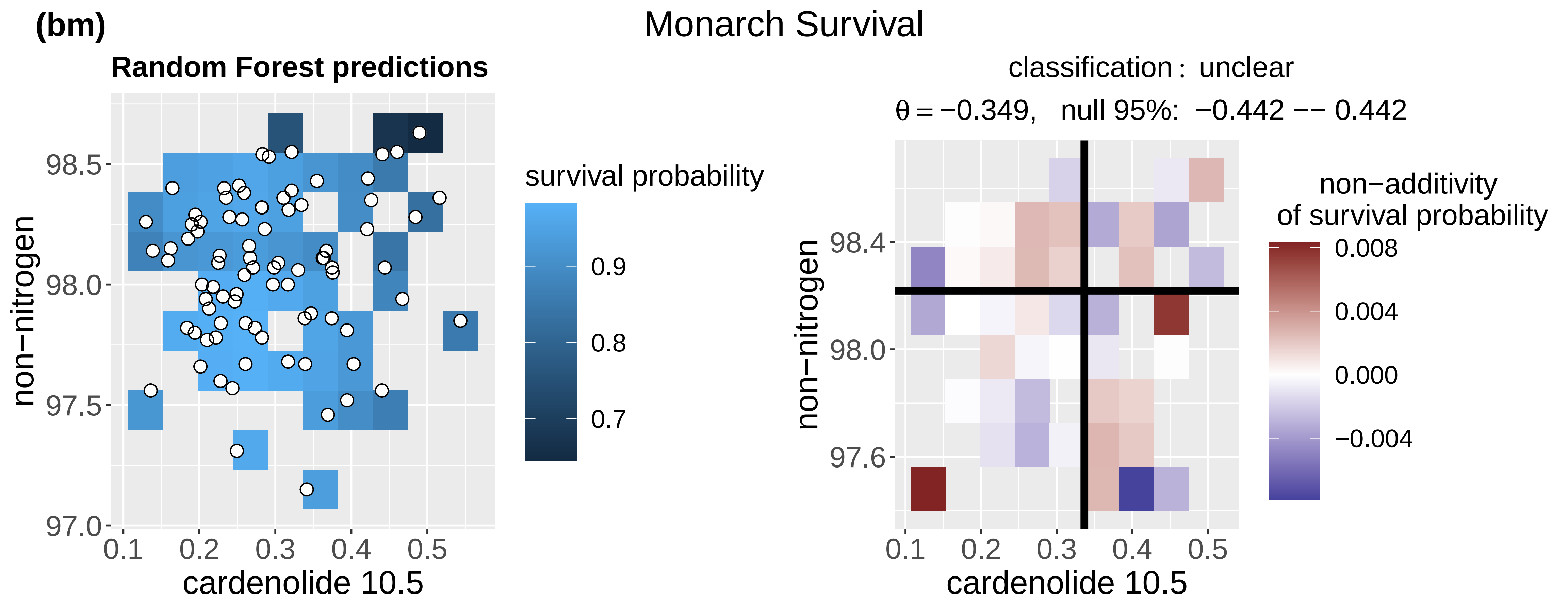
(bm)
Monarch Survival
Random Forest predictions
survival probability
0.9
0.8
0.7
non−nitrogen
cardenolide 10.5
97.0 97.5 98.0 98.5
0.1 0.2 0.3 0.4 0.5
classification: unclear
θ = −0.349, null 95%: −0.442 −− 0.442
non−additivity
of survival probability
0.008
0.004
0.000
−0.004
97.6 98.0 98.4

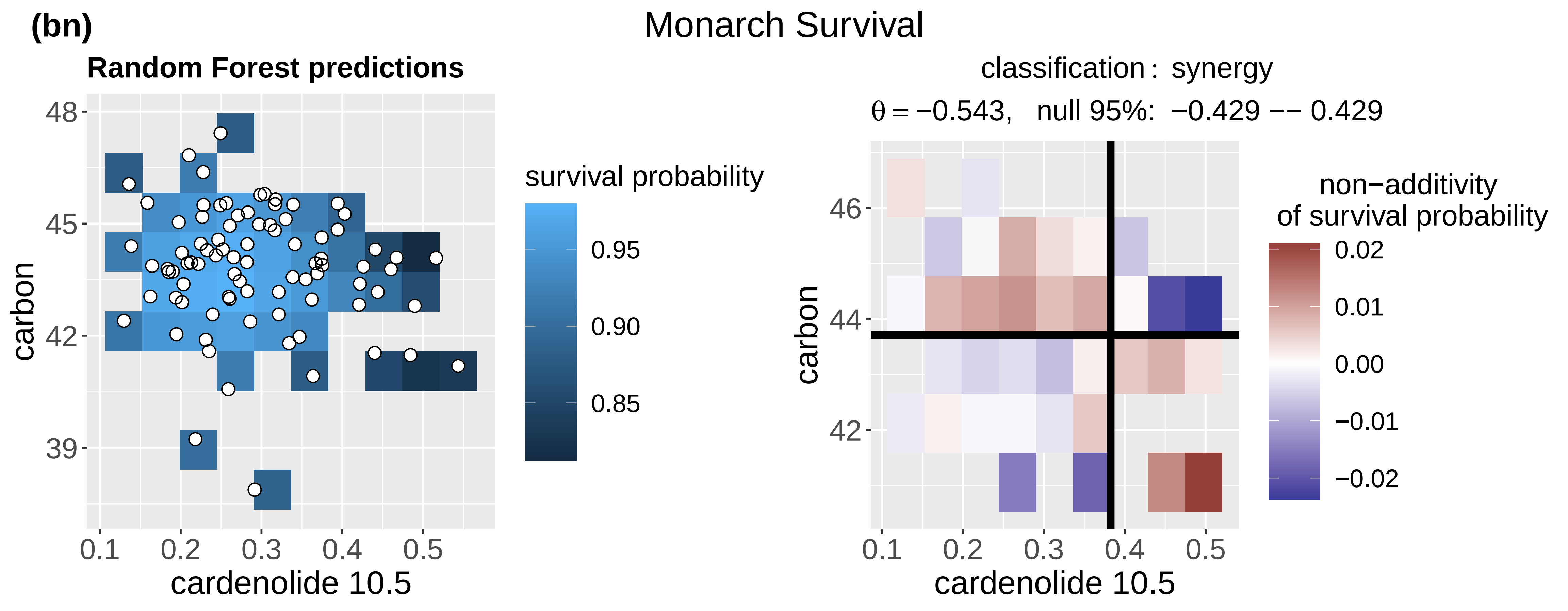
(bn)
Monarch Survival
Random Forest predictions
survival probability
0.95
0.90
0.85
carbon
cardenolide 10.5
39 42 45 48
0.1 0.2 0.3 0.4 0.5
classification: synergy
θ = −0.543, null 95%: −0.429 −− 0.429
non−additivity
of survival probability
0.02
0.01
0.00
−0.01
−0.02
42 44 46

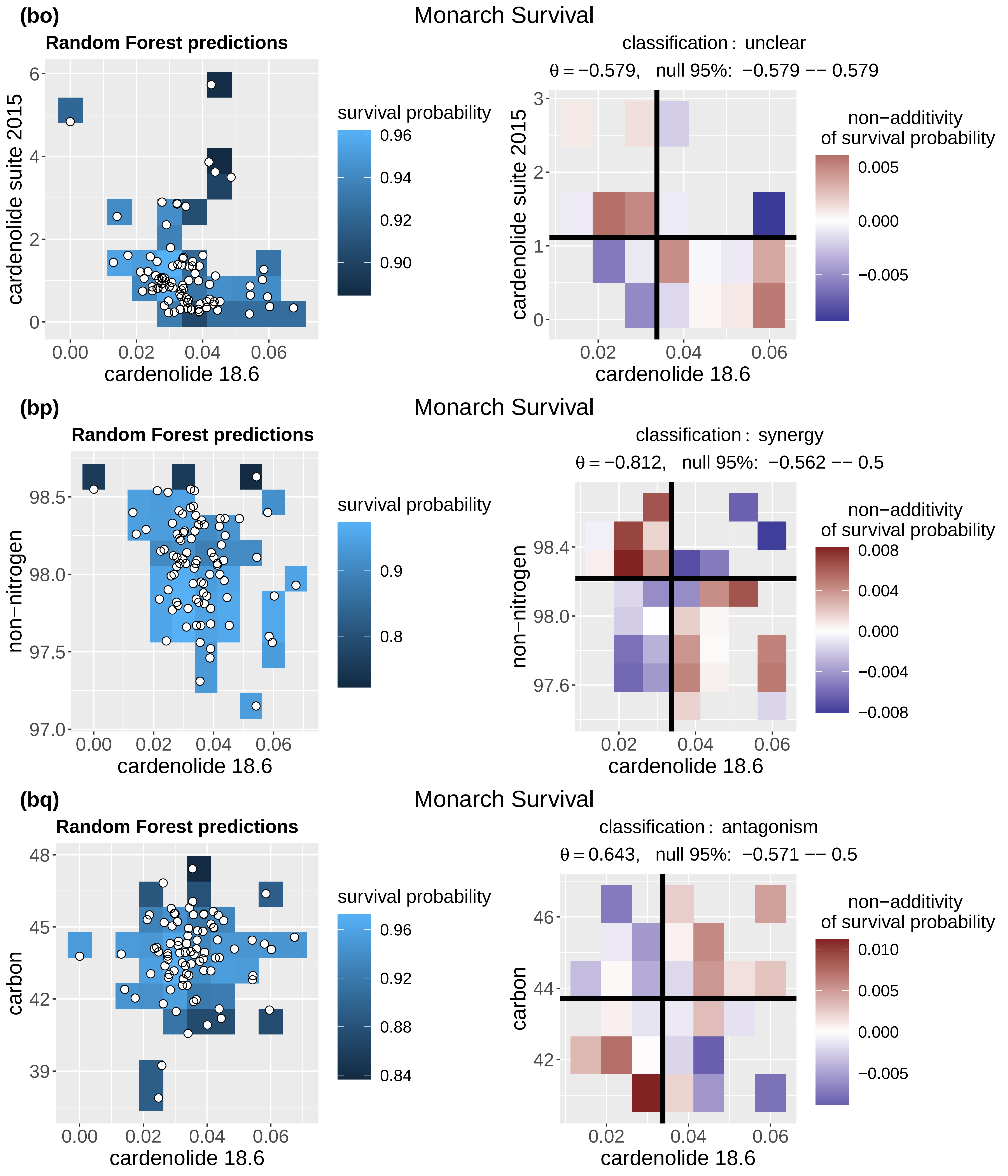
(bo)
Monarch Survival
Random Forest predictions
survival probability
0.96
0.94
0.92
0.90
cardenolide suite 2015
cardenolide 18.6
0.00 0.02 0.04 0.06
classification : unclear
θ = −0.579, null 95%: −0.579 −− 0.579
non−additivity
of survival probability
0.005
0.000
−0.005
(bp)
Monarch Survival
Random Forest predictions
survival probability
0.9
0.8
non−nitrogen
98.5
98.0
97.5
97.0
cardenolide 18.6
classification : synergy
θ = −0.812, null 95%: −0.562 −− 0.5
non−additivity
of survival probability
0.008
0.004
0.000
−0.004
−0.008
98.4
98.0
97.6
(bq)
Monarch Survival
Random Forest predictions
survival probability
0.96
0.92
0.88
0.84
carbon
48
45
42
39
cardenolide 18.6
classification : antagonism
θ = 0.643, null 95%: −0.571 −− 0.5
non−additivity
of survival probability
0.010
0.005
0.000
−0.005
46
44
42

(br)

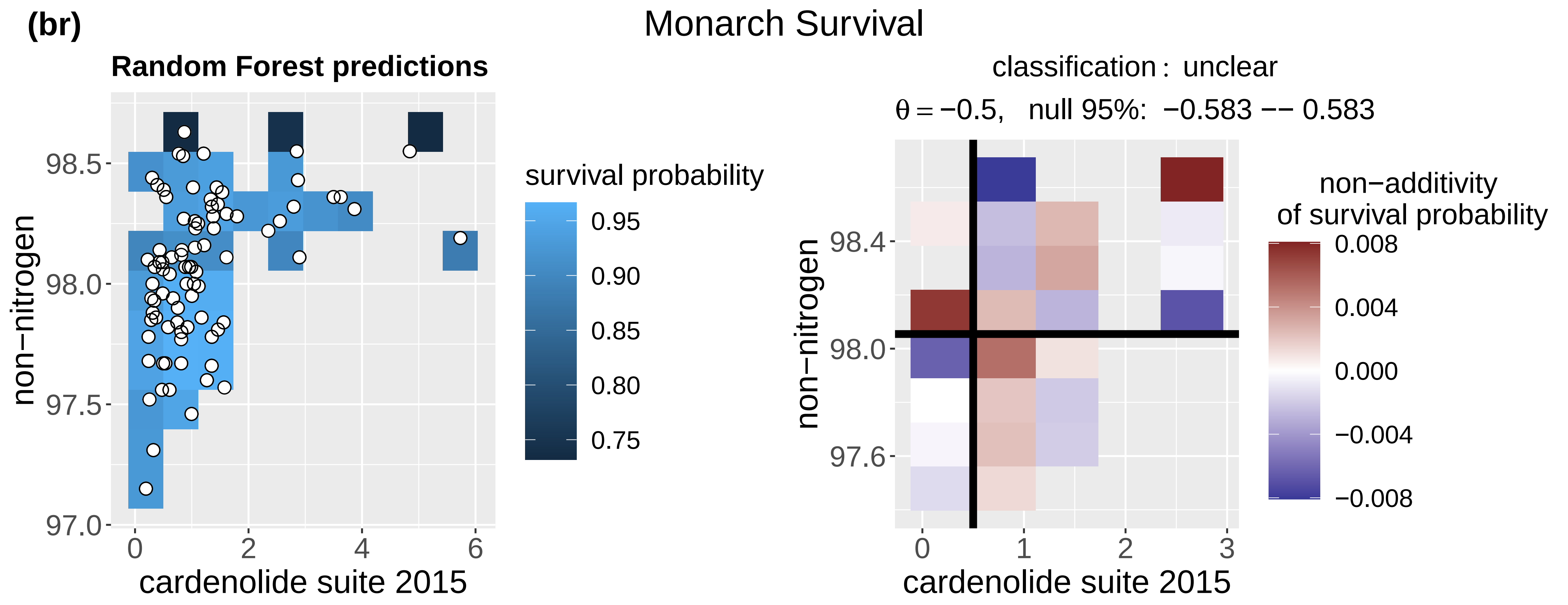
Monarch Survival
Random Forest predictions
survival probability
0.95
0.90
0.85
0.80
0.75
non−nitrogen
cardenolide suite 2015
classification : unclear
θ = −0.5,   null 95%:  −0.583 −− 0.583
non−additivity
of survival probability
0.008
0.004
0.000
−0.004
−0.008

(bs)

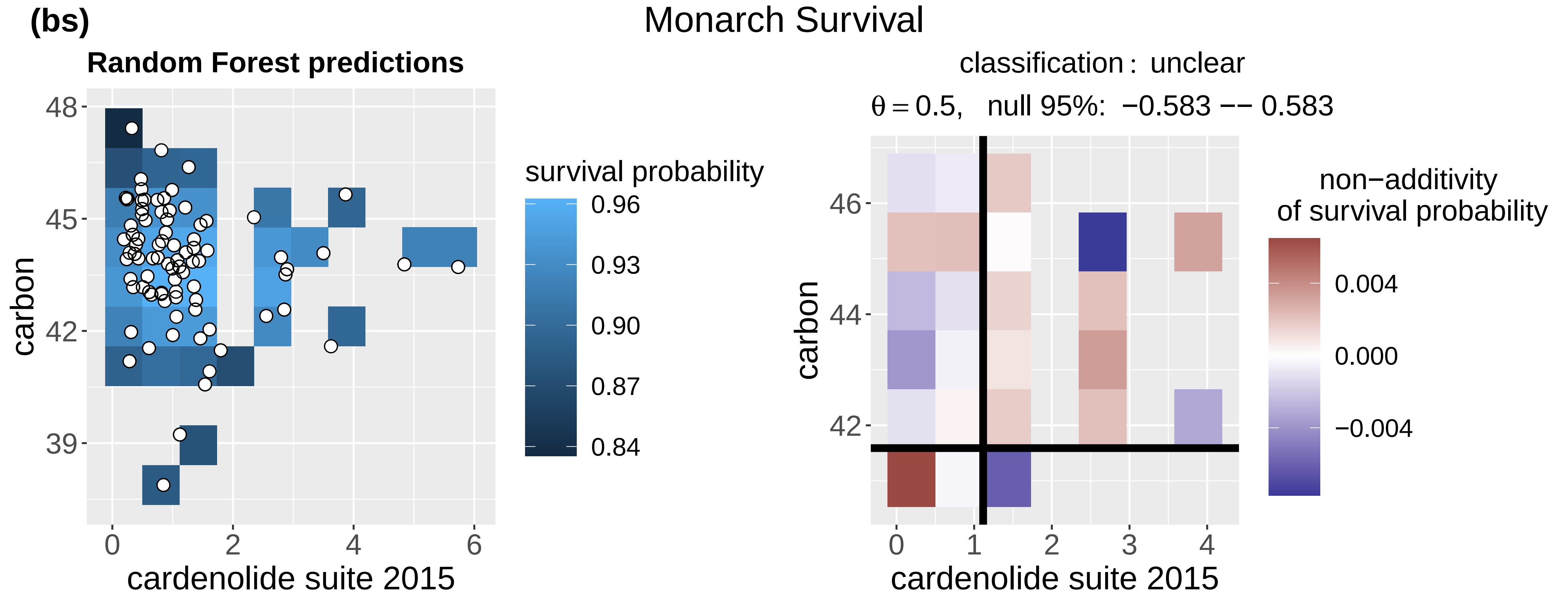
Monarch Survival
Random Forest predictions
survival probability
0.96
0.93
0.90
0.87
0.84
carbon
cardenolide suite 2015
classification : unclear
θ = 0.5,   null 95%:  −0.583 −− 0.583
non−additivity
of survival probability
0.004
0.000
−0.004

(bt)

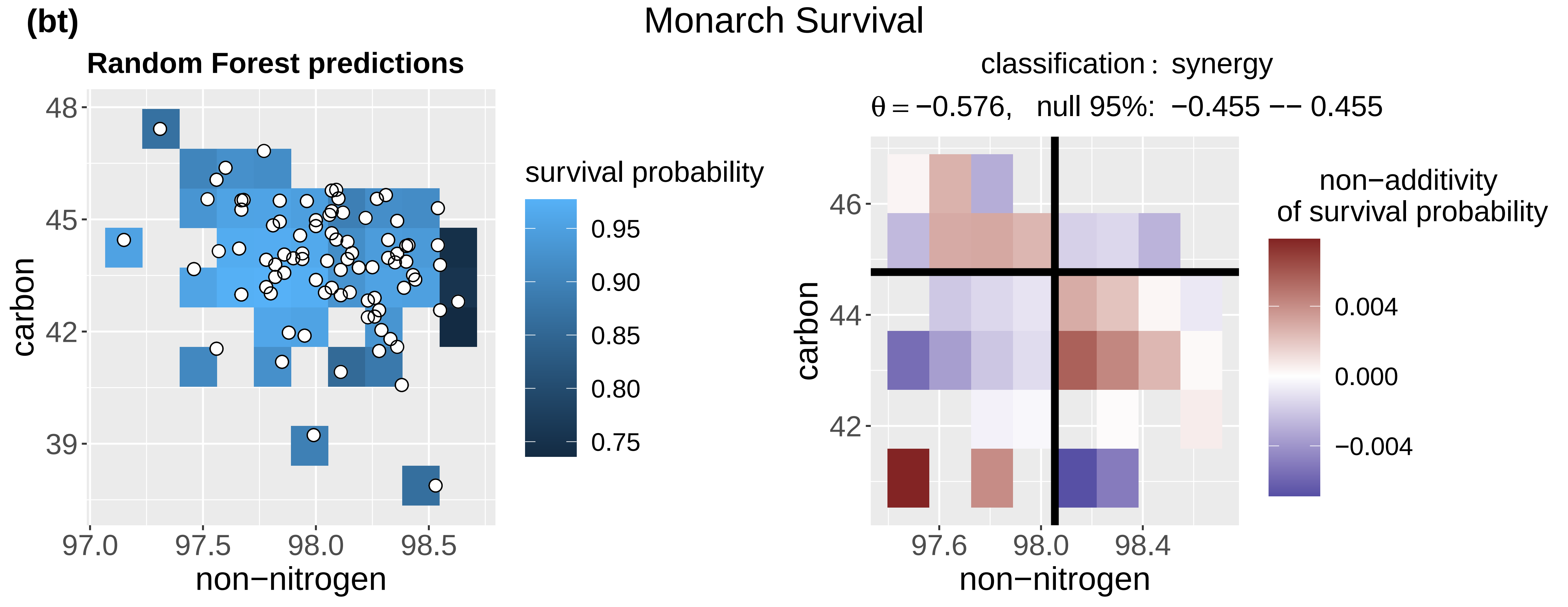
Monarch Survival
Random Forest predictions
survival probability
0.95
0.90
0.85
0.80
0.75
carbon
non−nitrogen
classification : synergy
θ = −0.576,   null 95%:  −0.455 −− 0.455
non−additivity
of survival probability
0.004
0.000
−0.004

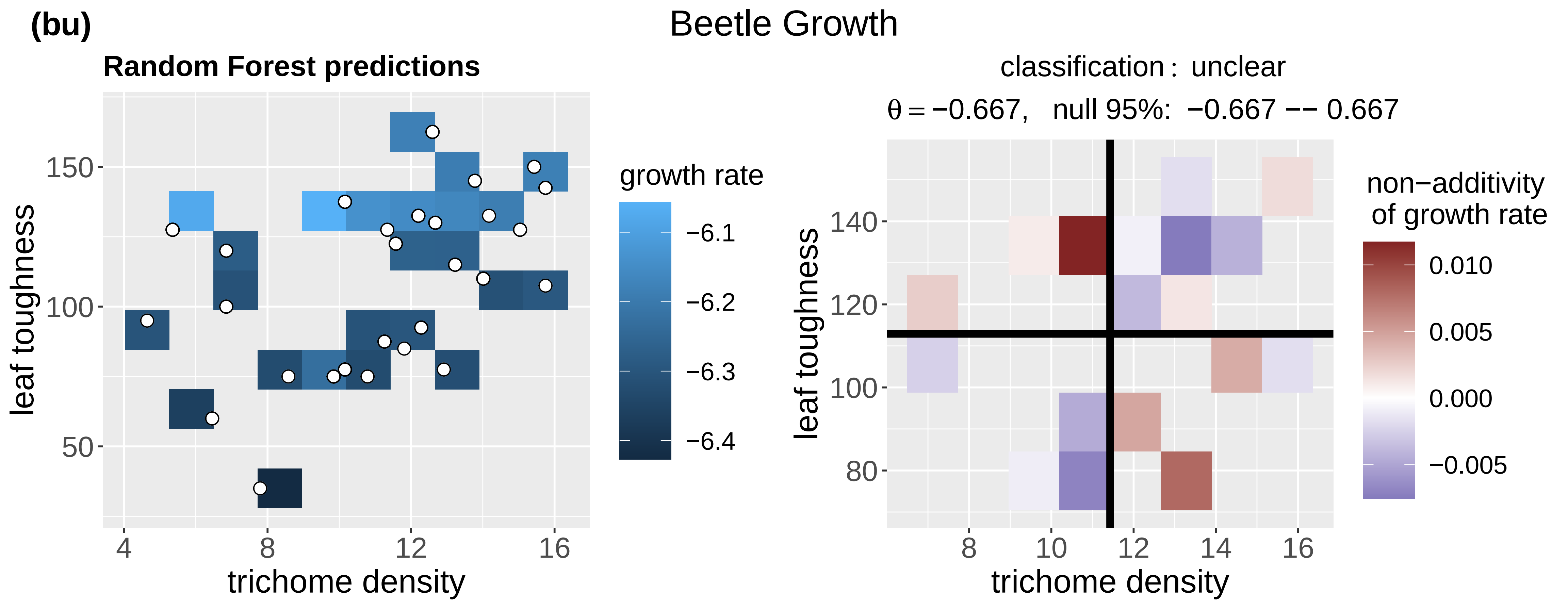
(bu)
Beetle Growth
Random Forest predictions
leaf toughness
trichome density
growth rate
−6.1
−6.2
−6.3
−6.4
classification : unclear
θ = −0.667, null 95%: −0.667 −− 0.667
non−additivity of growth rate
0.010
0.005
0.000
−0.005

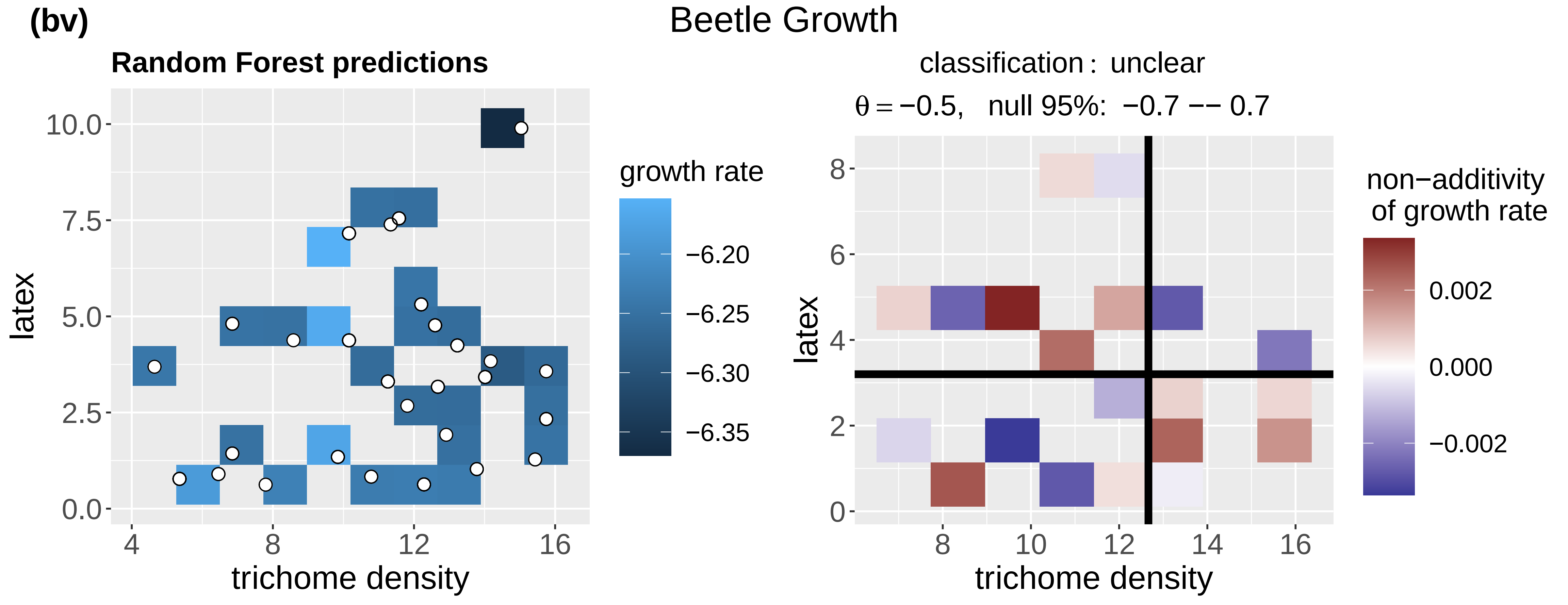
(bv)
Beetle Growth
Random Forest predictions
latex
trichome density
growth rate
−6.20
−6.25
−6.30
−6.35
classification : unclear
θ = −0.5, null 95%: −0.7 −− 0.7
non−additivity of growth rate
0.002
0.000
−0.002

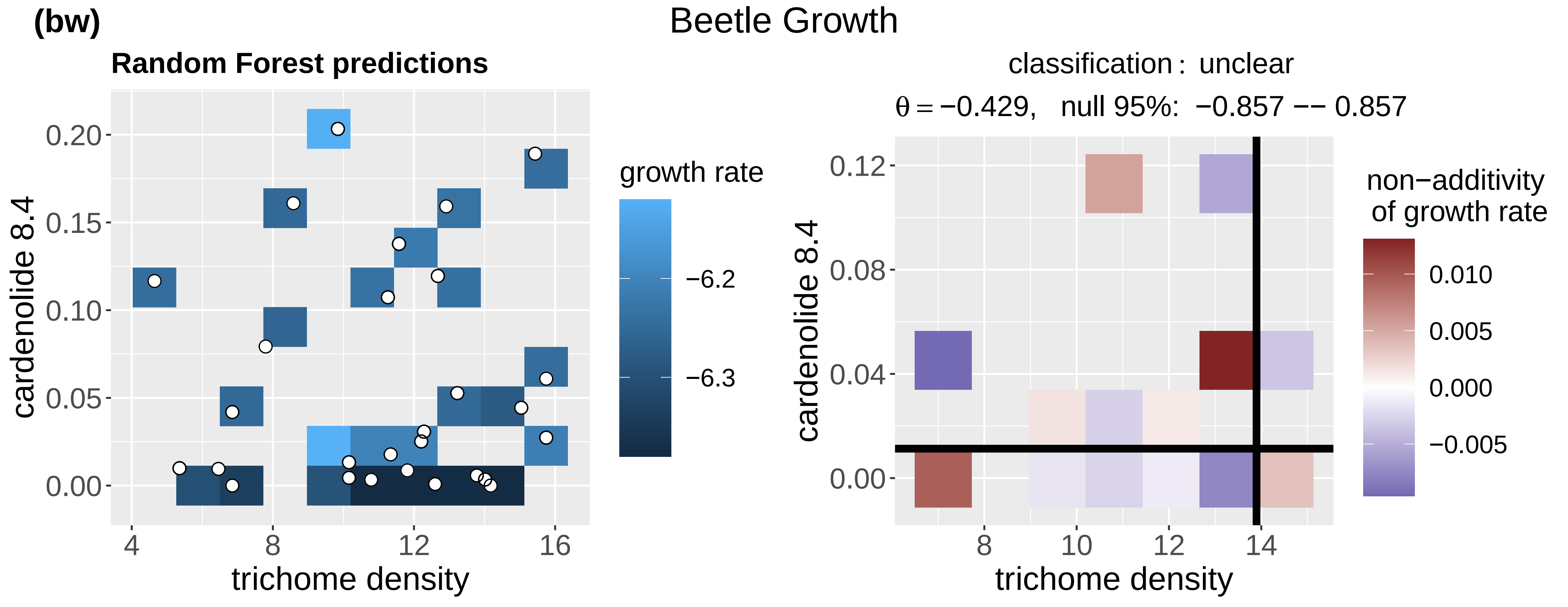
(bw)
Beetle Growth
Random Forest predictions
cardenolide 8.4
trichome density
growth rate
−6.2
−6.3
classification : unclear
θ = −0.429, null 95%: −0.857 −− 0.857
non−additivity of growth rate
0.010
0.005
0.000
−0.005

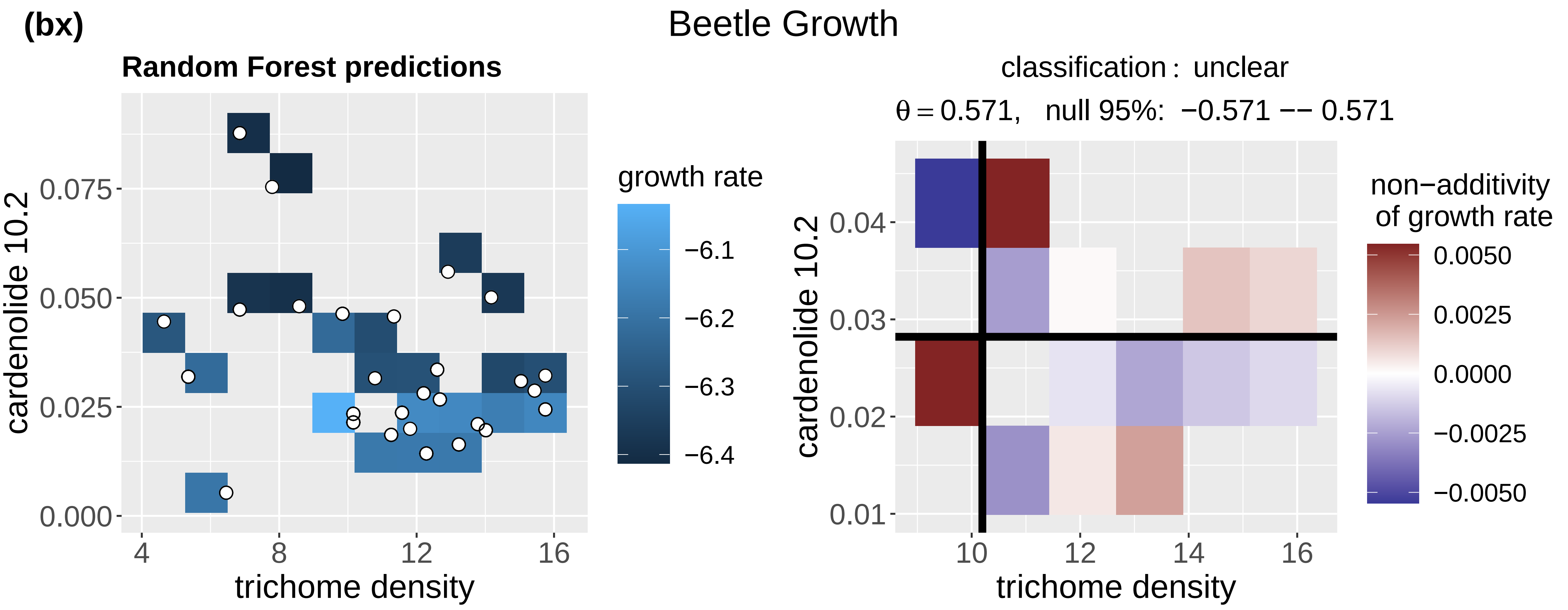
(bx)
Beetle Growth
Random Forest predictions
cardenolide 10.2
trichome density
growth rate
classification : unclear
θ = 0.571, null 95%: −0.571 −− 0.571
non−additivity of growth rate

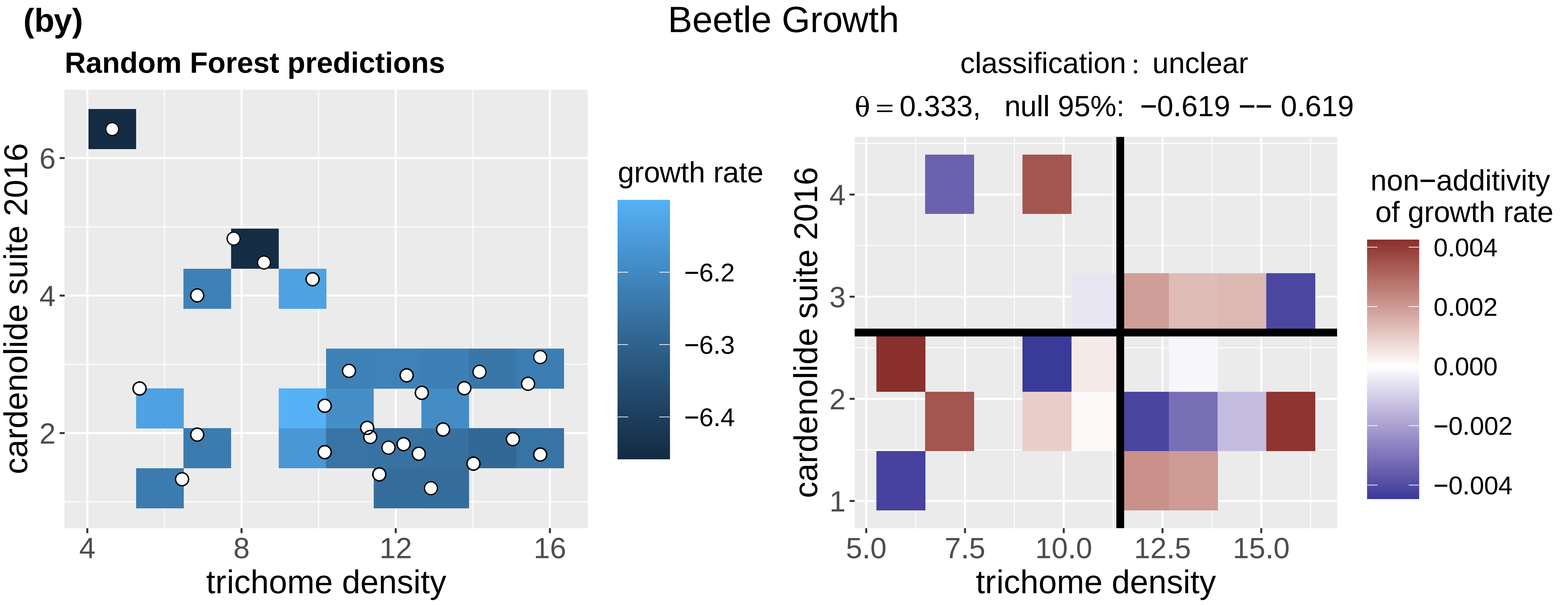
(by)
Beetle Growth
Random Forest predictions
cardenolide suite 2016
trichome density
growth rate
classification : unclear
θ = 0.333, null 95%: −0.619 −− 0.619
non−additivity of growth rate

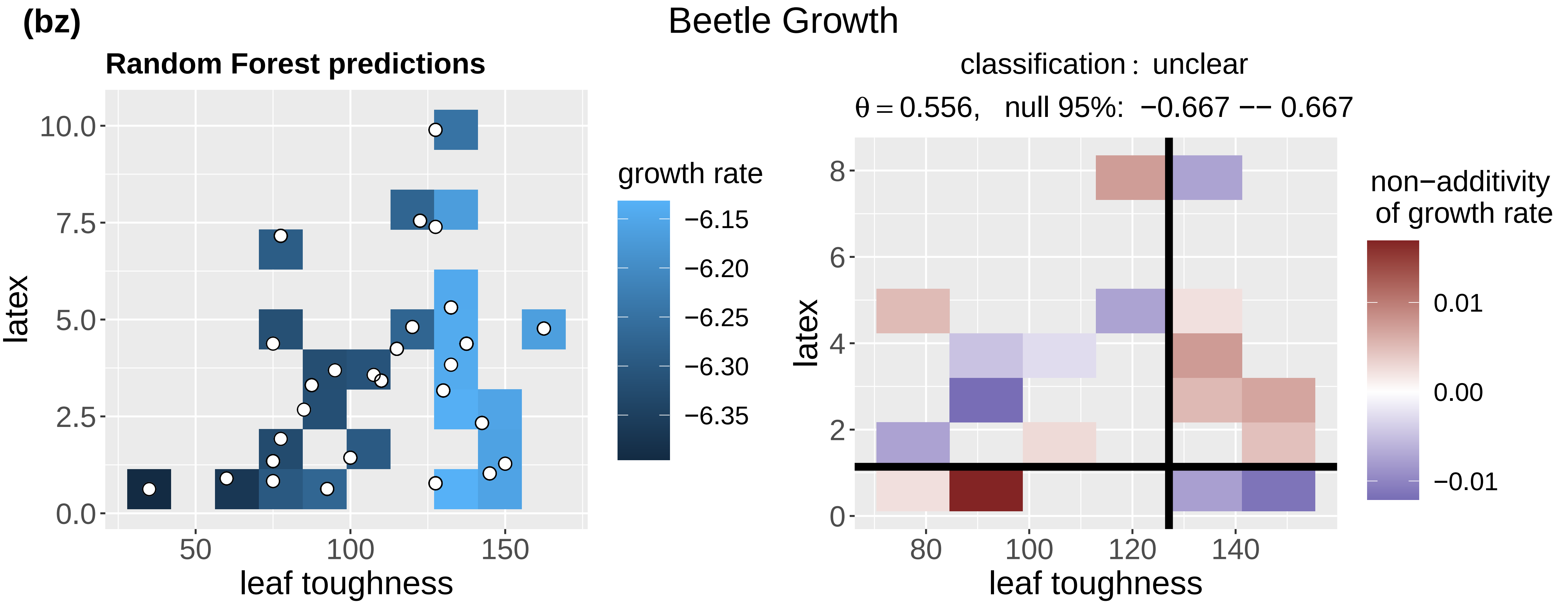
(bz)
Beetle Growth
Random Forest predictions
latex
leaf toughness
growth rate
classification : unclear
θ = 0.556, null 95%: −0.667 −− 0.667
non−additivity of growth rate

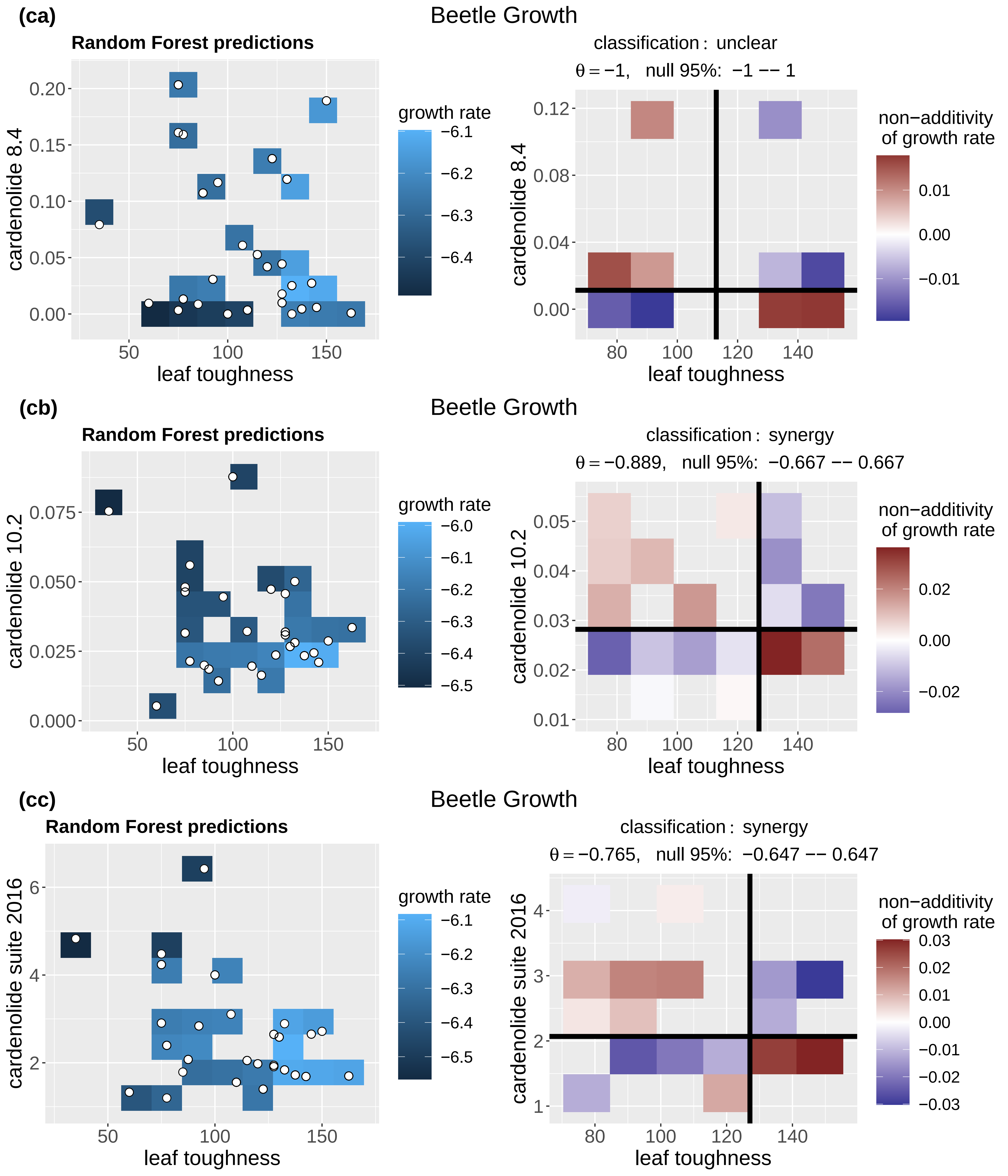
(ca)
Beetle Growth
Random Forest predictions
cardenolide 8.4
leaf toughness
growth rate
classification : unclear
θ = −1, null 95%: −1 −− 1
non−additivity of growth rate
(cb)
Beetle Growth
Random Forest predictions
cardenolide 10.2
leaf toughness
growth rate
classification : synergy
θ = −0.889, null 95%: −0.667 −− 0.667
non−additivity of growth rate
(cc)
Beetle Growth
Random Forest predictions
cardenolide suite 2016
leaf toughness
growth rate
classification : synergy
θ = −0.765, null 95%: −0.647 −− 0.647
non−additivity of growth rate

(cd)

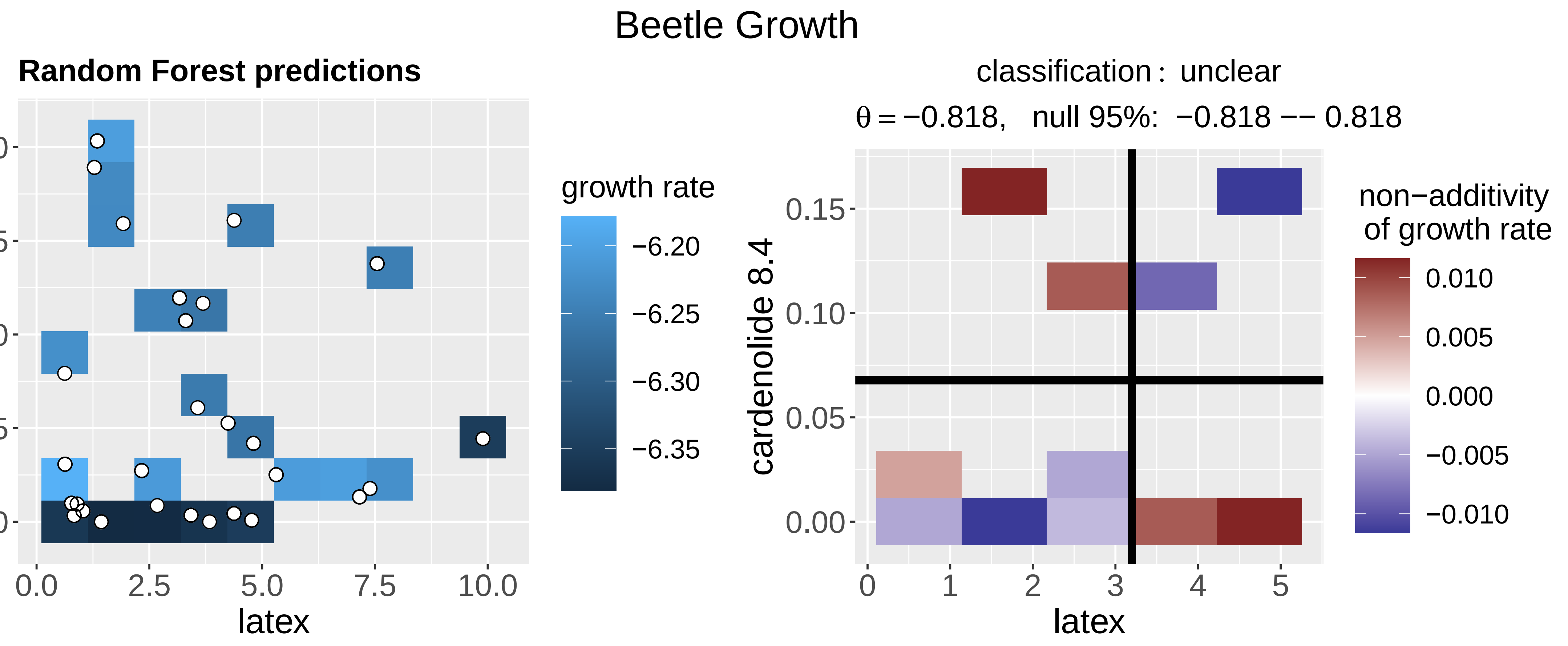
Beetle Growth
Random Forest predictions
cardenolide 8.4
latex
growth rate
classification : unclear
θ = −0.818, null 95%: −0.818 −− 0.818
non−additivity of growth rate

(ce)

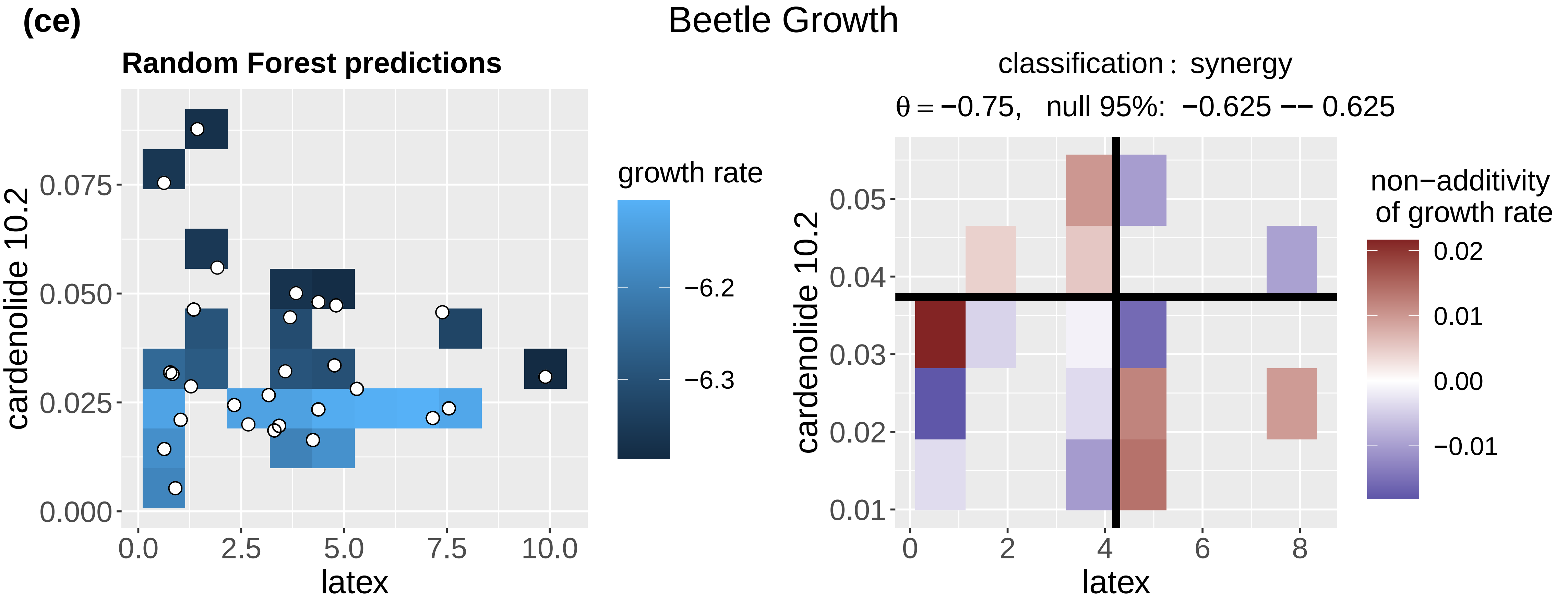
Beetle Growth
Random Forest predictions
cardenolide 10.2
latex
growth rate
classification : synergy
θ = −0.75, null 95%: −0.625 −− 0.625
non−additivity of growth rate

(cf)

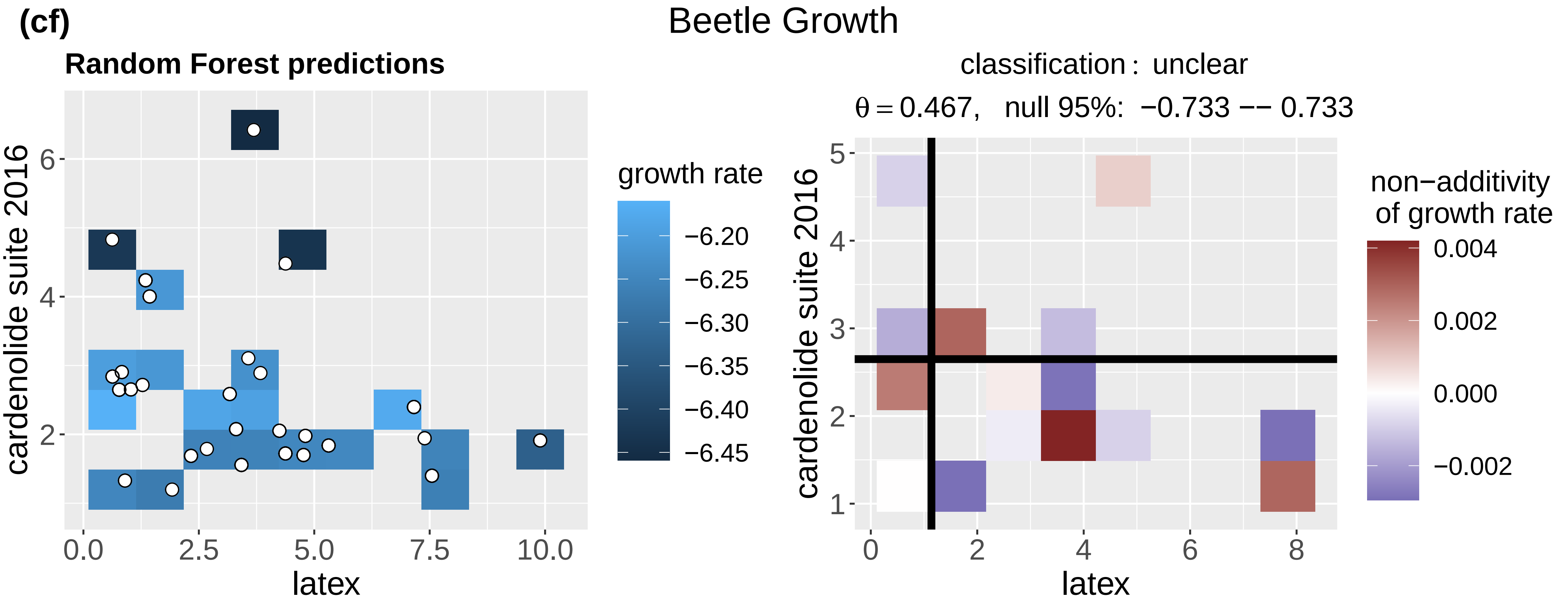
Beetle Growth
Random Forest predictions
cardenolide suite 2016
latex
growth rate
classification : unclear
θ = 0.467, null 95%: −0.733 −− 0.733
non−additivity of growth rate

(cg)

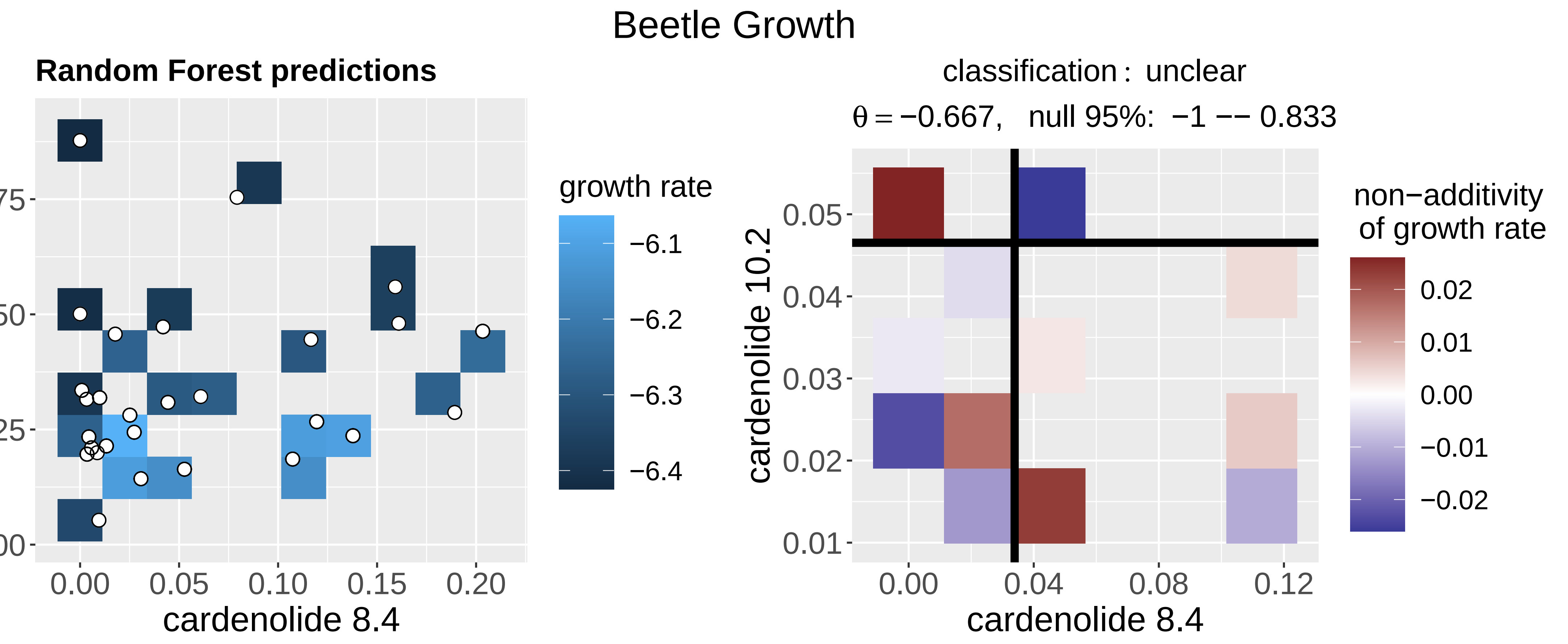
Beetle Growth
Random Forest predictions
cardenolide 10.2
cardenolide 8.4
growth rate
classification : unclear
θ = −0.667, null 95%: −1 −− 0.833
non−additivity of growth rate

(ch)

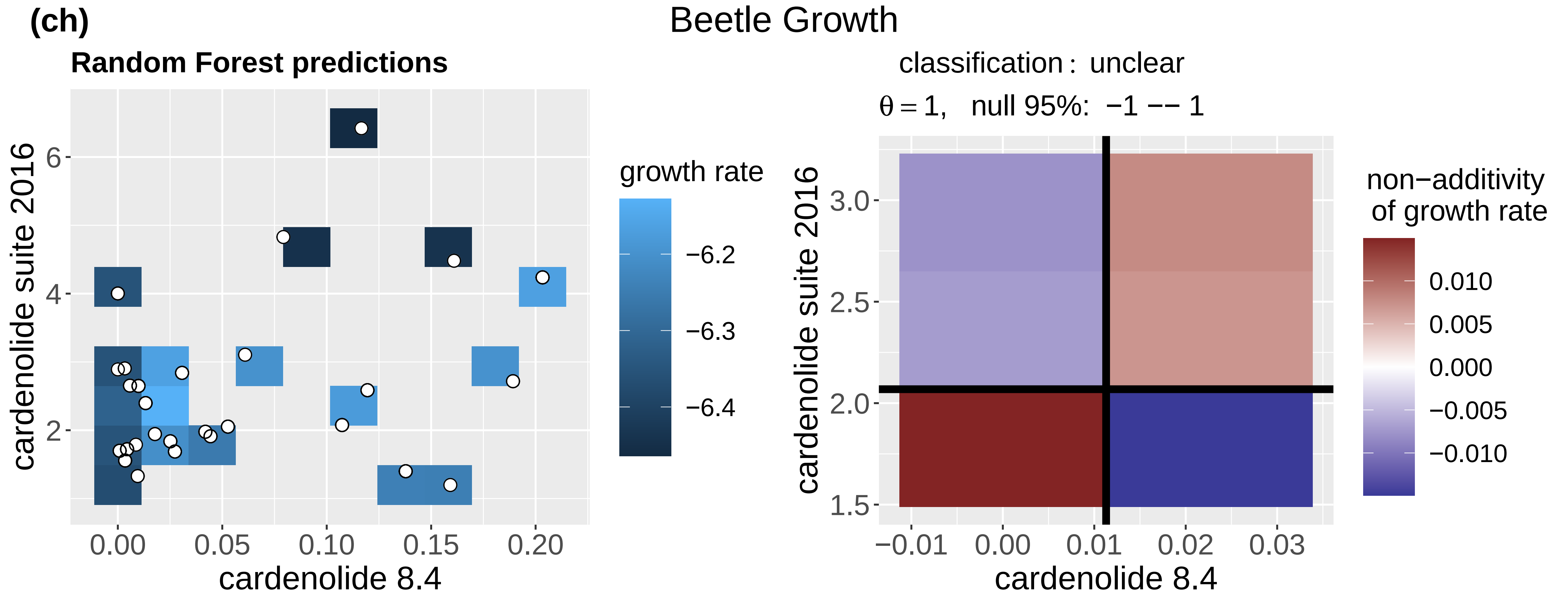
Beetle Growth
Random Forest predictions
cardenolide suite 2016
cardenolide 8.4
growth rate
classification : unclear
θ = 1, null 95%: −1 −− 1
non−additivity of growth rate

(ci)

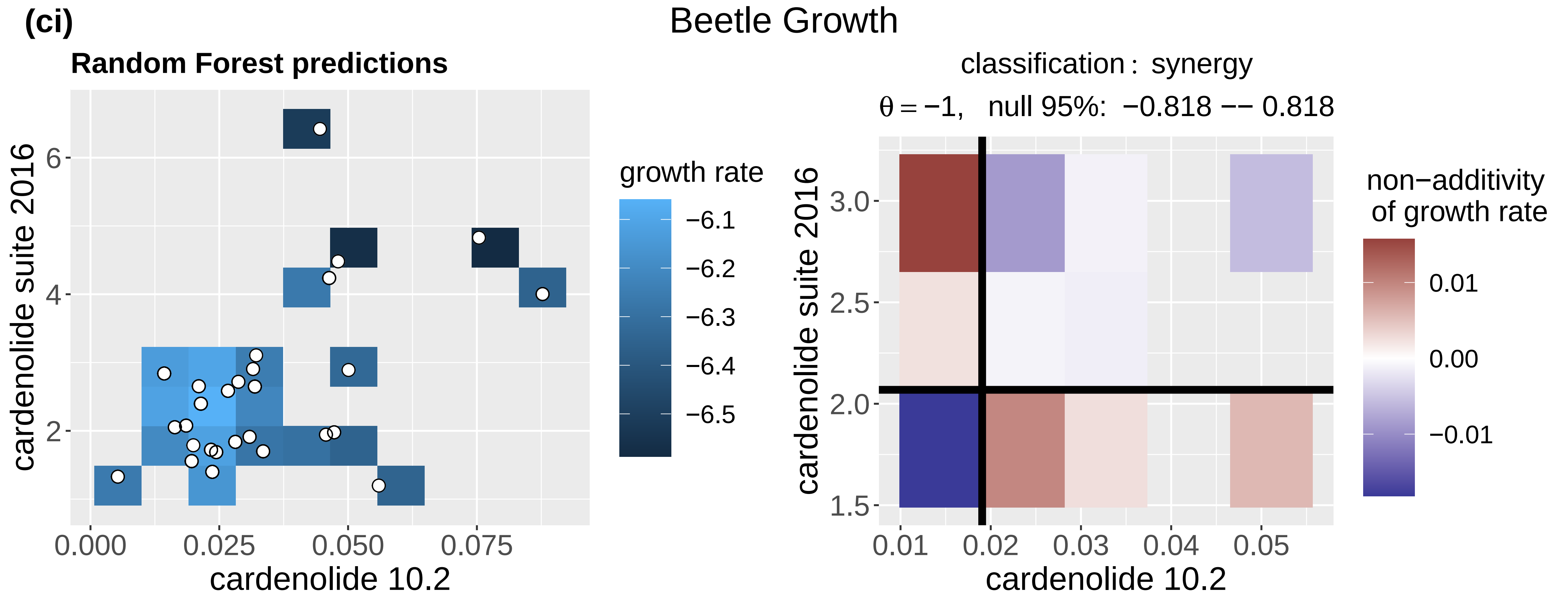
Beetle Growth
Random Forest predictions
cardenolide suite 2016
cardenolide 10.2
growth rate
classification : synergy
θ = −1, null 95%: −0.818 −− 0.818
non−additivity of growth rate

(cj) Beetle Survival

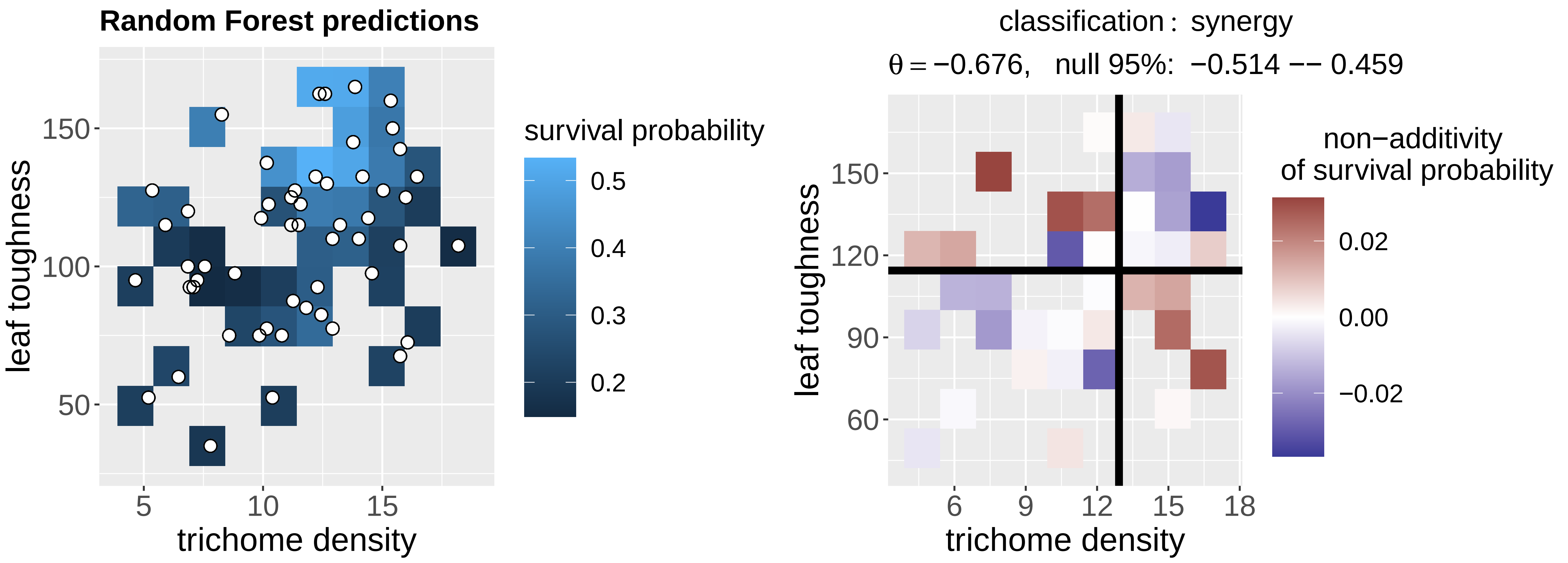


(ck) Beetle Survival

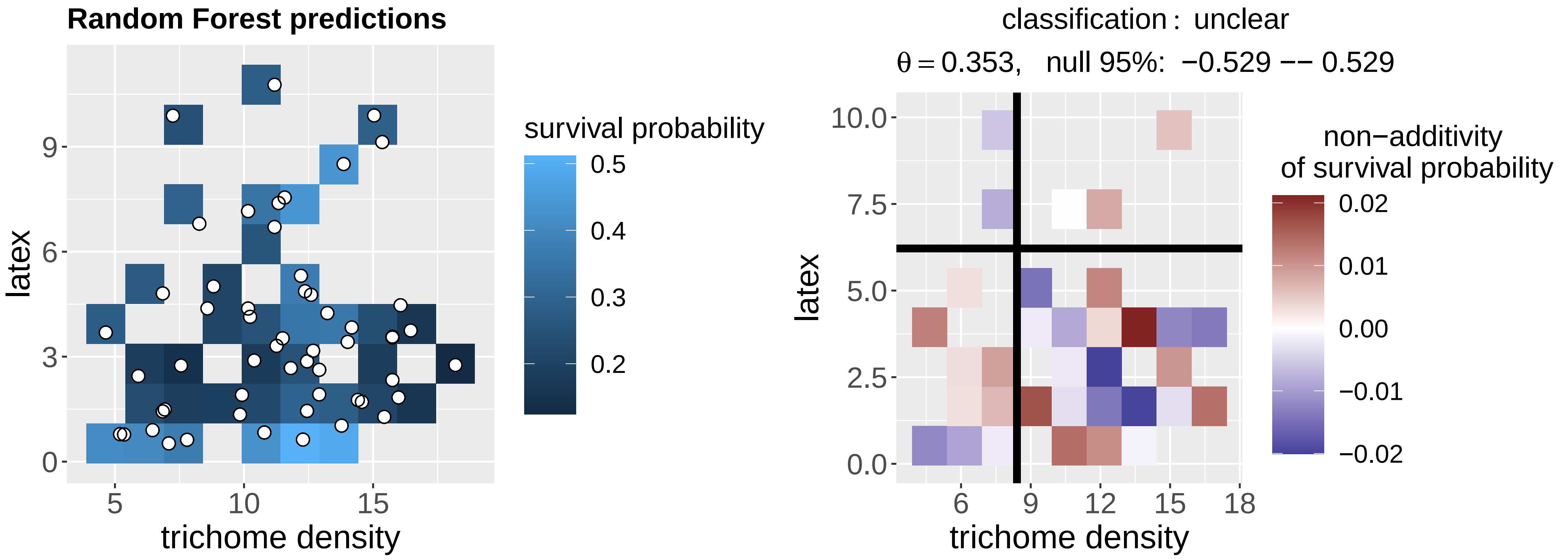


(cl) Beetle Survival

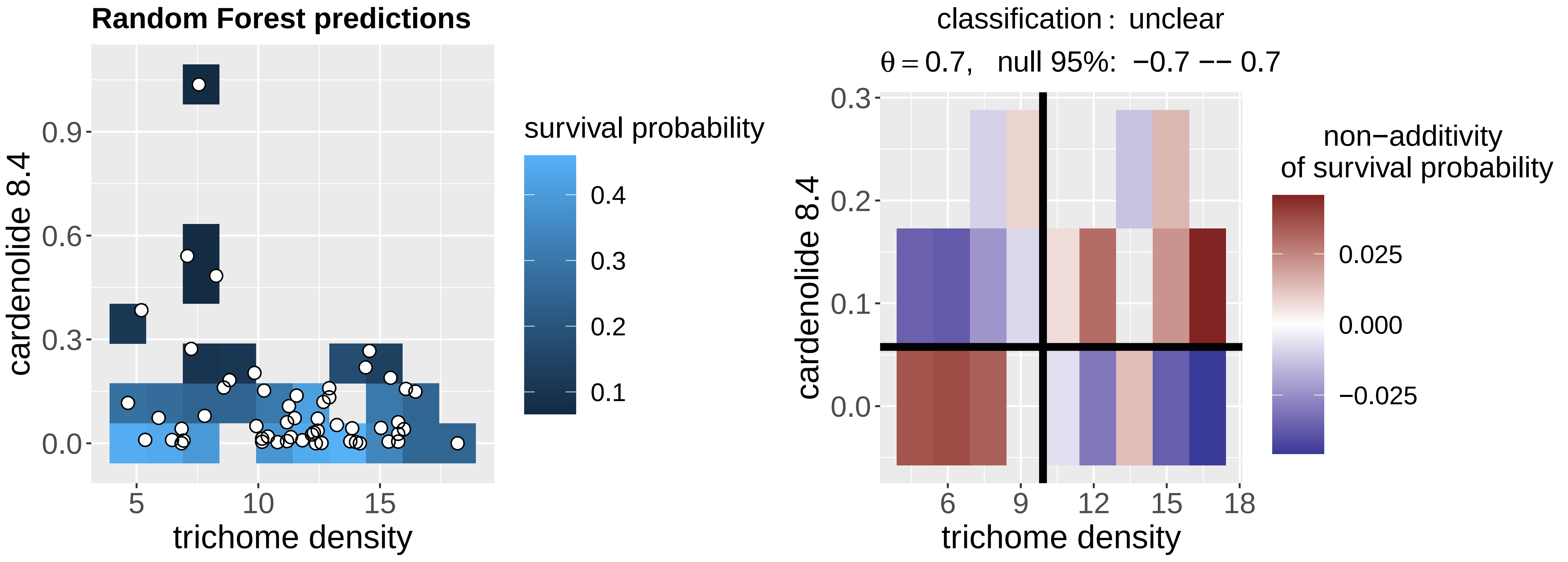

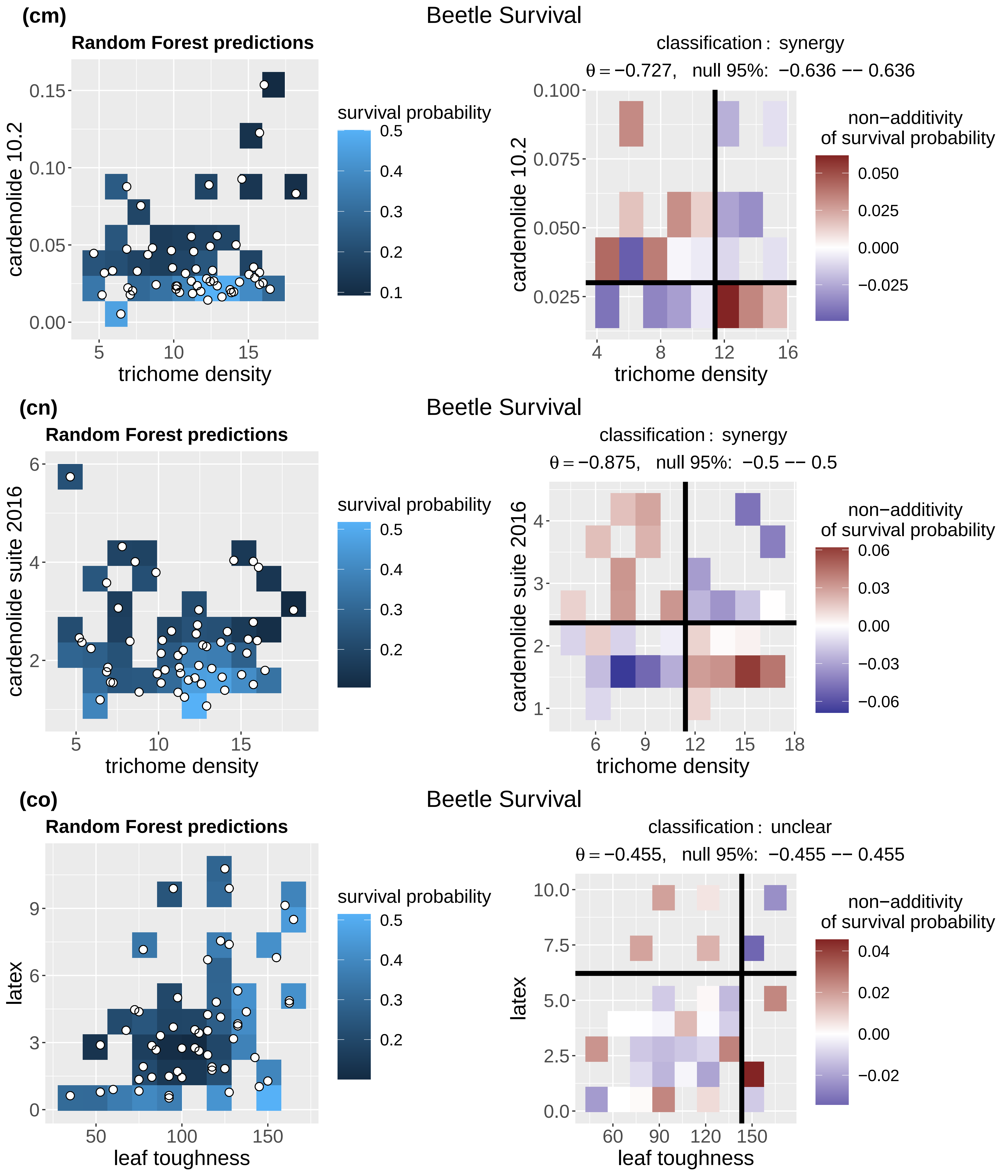
(cm)
Beetle Survival
Random Forest predictions
cardenolide 10.2
trichome density
survival probability
classification : synergy
θ = −0.727, null 95%: −0.636 −− 0.636
non−additivity of survival probability
(cn)
Beetle Survival
Random Forest predictions
cardenolide suite 2016
trichome density
survival probability
classification : synergy
θ = −0.875, null 95%: −0.5 −− 0.5
non−additivity of survival probability
(co)
Beetle Survival
Random Forest predictions
latex
leaf toughness
survival probability
classification : unclear
θ = −0.455, null 95%: −0.455 −− 0.455
non−additivity of survival probability

(cp)

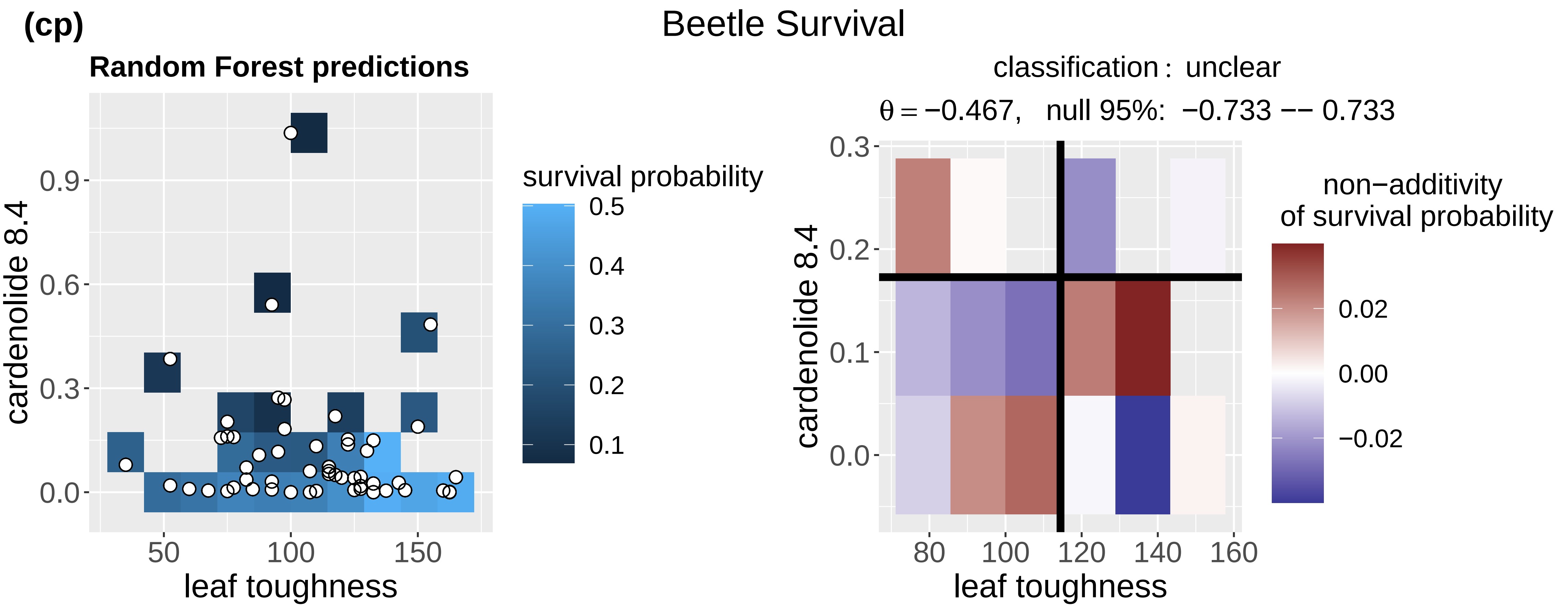
Beetle Survival
Random Forest predictions
cardenolide 8.4
leaf toughness
survival probability
classification: unclear
θ = −0.467, null 95%: −0.733 −− 0.733
non−additivity
of survival probability
leaf toughness
cardenolide 8.4

(cq)

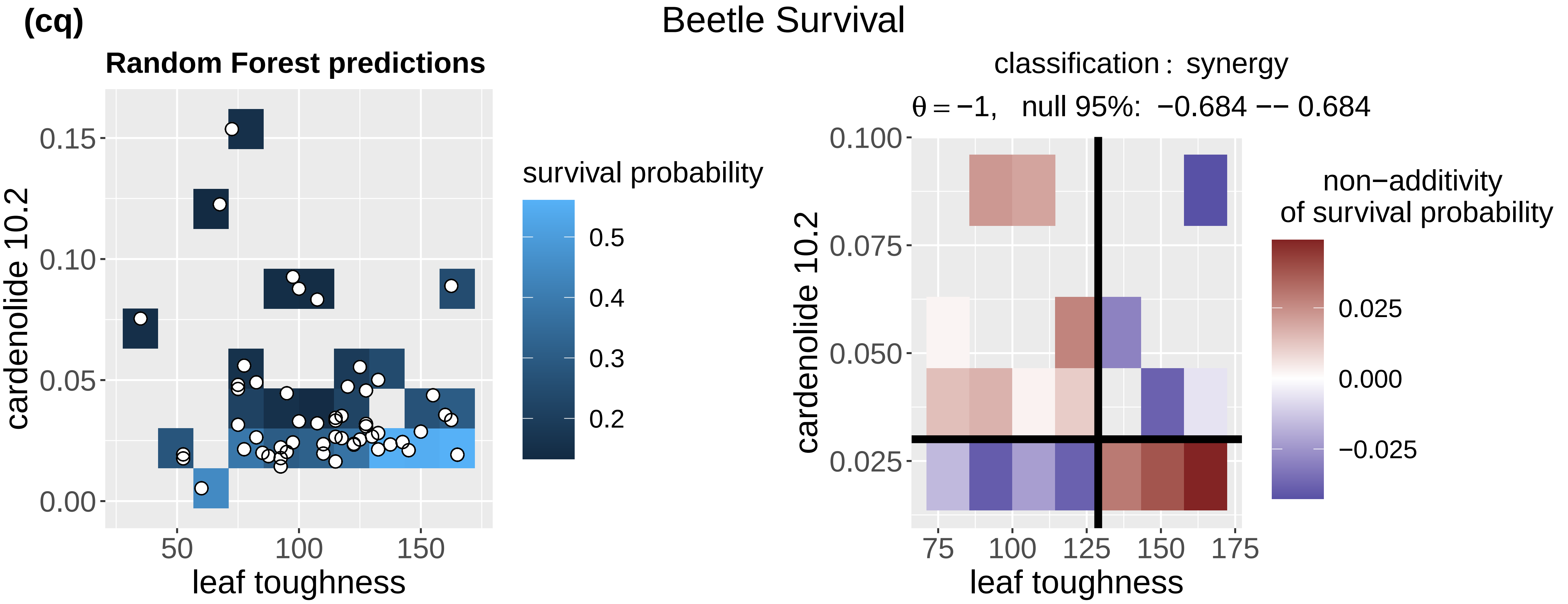
Beetle Survival
Random Forest predictions
cardenolide 10.2
leaf toughness
survival probability
classification: synergy
θ = −1, null 95%: −0.684 −− 0.684
non−additivity
of survival probability
leaf toughness
cardenolide 10.2

(cr)

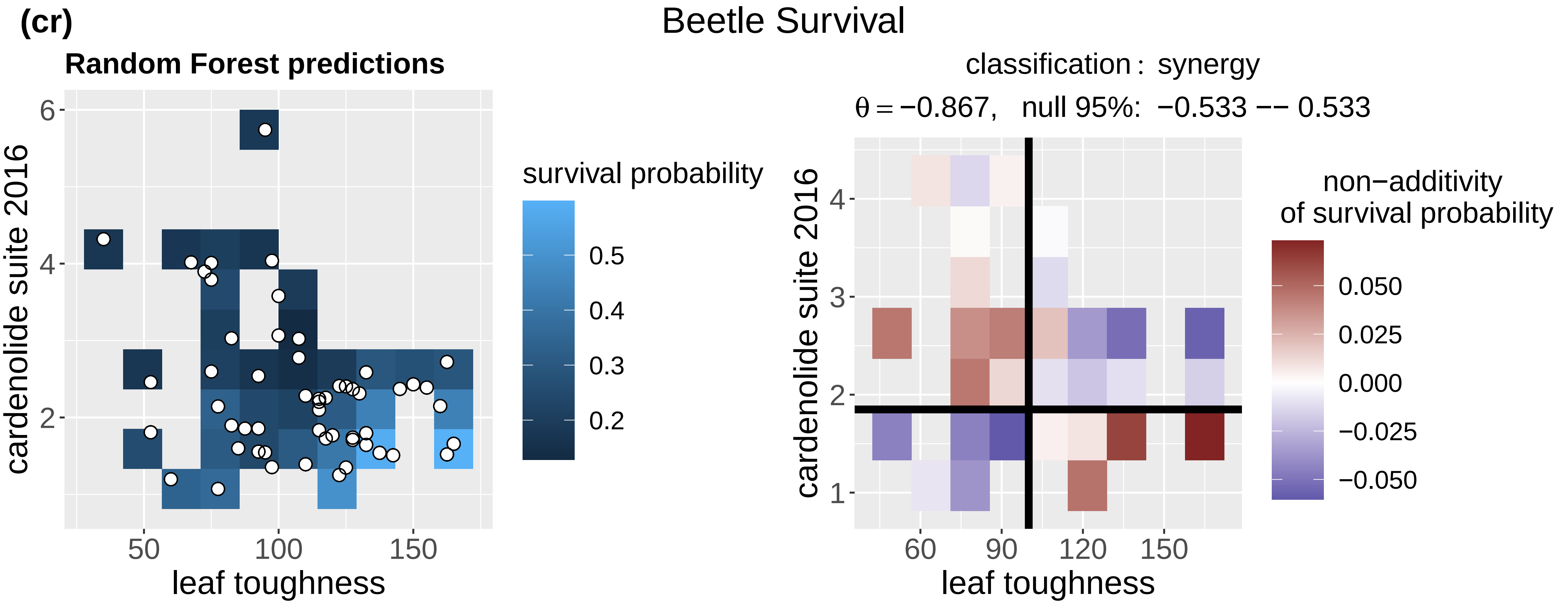
Beetle Survival
Random Forest predictions
cardenolide suite 2016
leaf toughness
survival probability
classification: synergy
θ = −0.867, null 95%: −0.533 −− 0.533
non−additivity
of survival probability
leaf toughness
cardenolide suite 2016

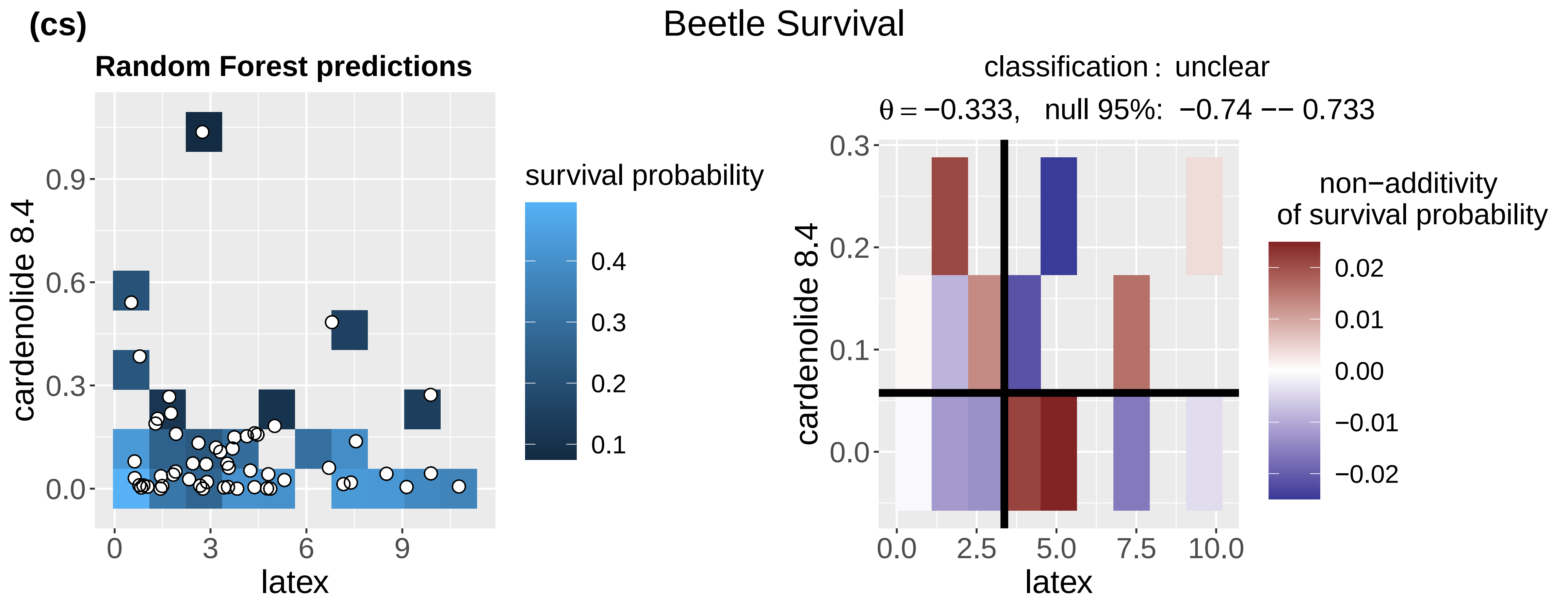
(cs)
Beetle Survival
Random Forest predictions
cardenolide 8.4
0.0
0.3
0.6
0.9
latex
0
3
6
9
survival probability
0.4
0.3
0.2
0.1
classification : unclear
θ = −0.333, null 95%: −0.74 −− 0.733
0.3
0.2
0.1
0.0
0.0
2.5
5.0
7.5
10.0
non−additivity
of survival probability
0.02
0.01
0.00
−0.01
−0.02

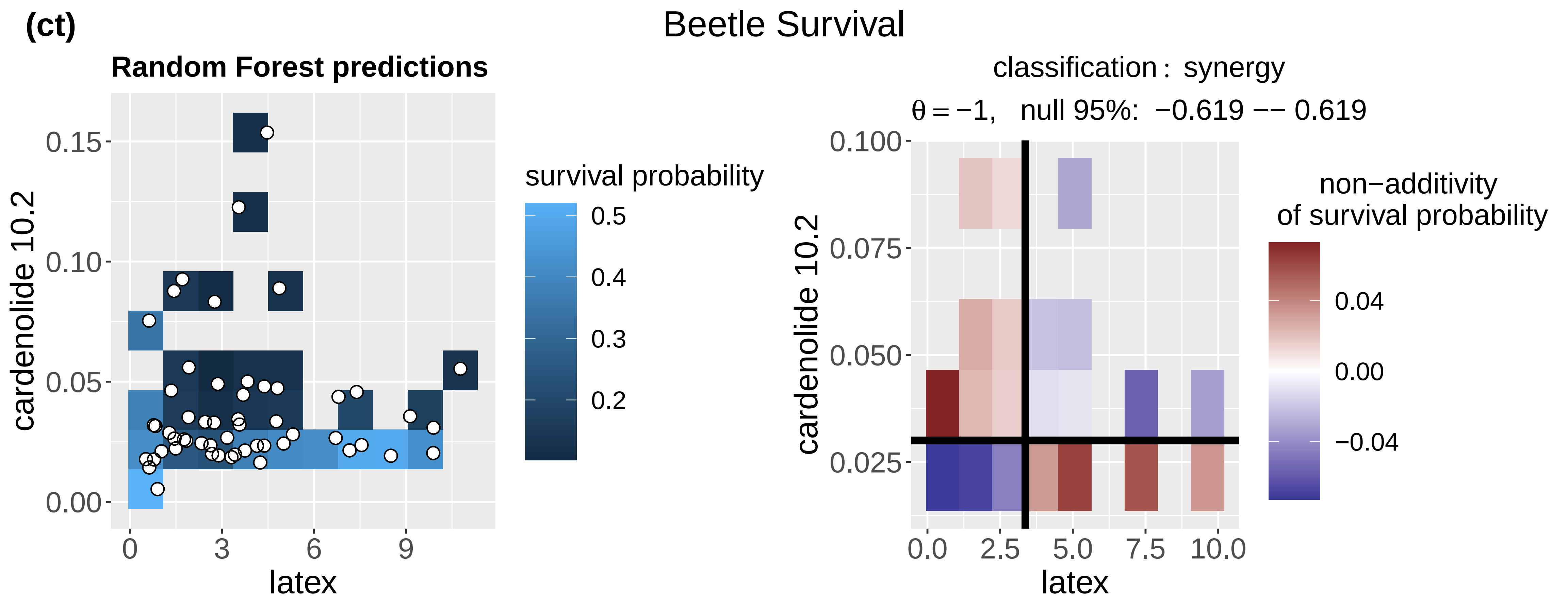
(ct)
Beetle Survival
Random Forest predictions
cardenolide 10.2
0.00
0.05
0.10
0.15
latex
0
3
6
9
survival probability
0.5
0.4
0.3
0.2
classification : synergy
θ = −1, null 95%: −0.619 −− 0.619
0.100
0.075
0.050
0.025
0.0
2.5
5.0
7.5
10.0
non−additivity
of survival probability
0.04
0.00
−0.04

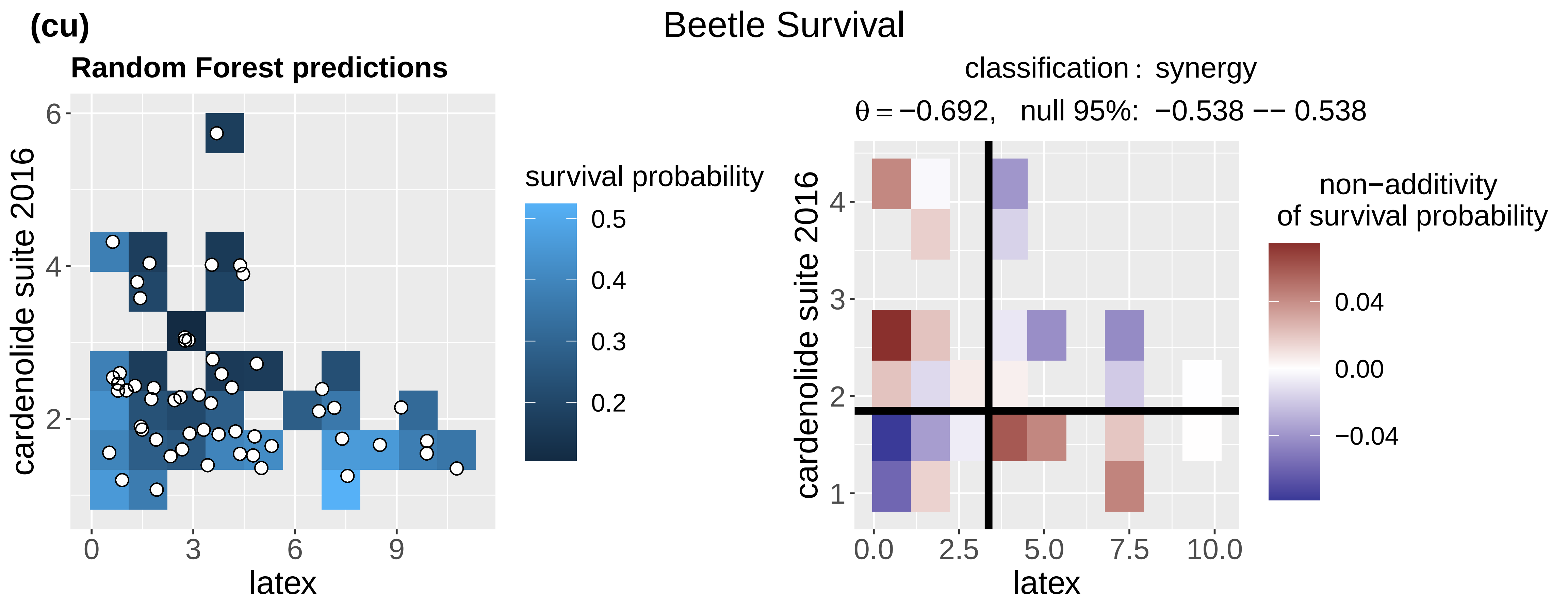
(cu)
Beetle Survival
Random Forest predictions
cardenolide suite 2016
2
4
6
latex
0
3
6
9
survival probability
0.5
0.4
0.3
0.2
classification : synergy
θ = −0.692, null 95%: −0.538 −− 0.538
1
2
3
4
0.0
2.5
5.0
7.5
10.0
non−additivity
of survival probability
0.04
0.00
−0.04

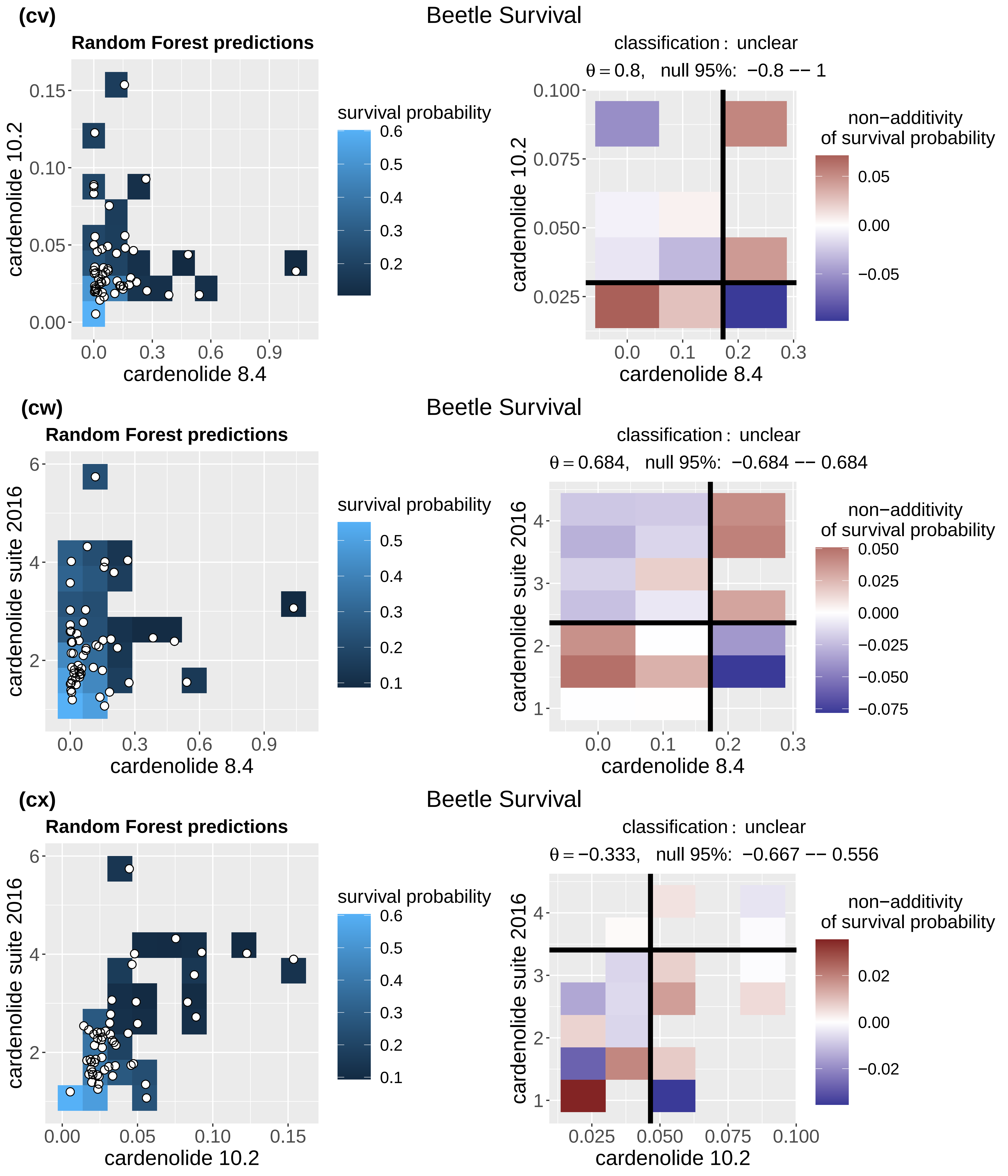
(cv)
Beetle Survival
Random Forest predictions
cardenolide 10.2
cardenolide 8.4
survival probability
classification : unclear
θ = 0.8, null 95%: −0.8 −− 1
non−additivity
of survival probability
(cw)
Beetle Survival
Random Forest predictions
cardenolide suite 2016
cardenolide 8.4
survival probability
classification : unclear
θ = 0.684, null 95%: −0.684 −− 0.684
non−additivity
of survival probability
(cx)
Beetle Survival
Random Forest predictions
cardenolide suite 2016
cardenolide 10.2
survival probability
classification : unclear
θ = −0.333, null 95%: −0.667 −− 0.556
non−additivity
of survival probability
